# Circadian rhythm in the atrioventricular node - in electrophysiology, contractility, metabolism, intracellular signalling and extracellular matrix

**DOI:** 10.64898/2026.11.10.765573

**Authors:** Sunil Jit R.J. Logantha, Claire Wilson, Sooraj V. Nair, Pierre Fontanaud, Eleonora Torre, Roman Tikhomirov, Sami Al-Othman, Mathijs van Schie, Rama Ijaz, Anatoliy Shmygol, Gilles Bru-Mercier, Yanwen Wang, Michael Coleman, George Hart, Munir Hussain, Hugh Piggins, Morten B. Thomsen, Shin Inada, Frank C. Howarth, Natasja de Groot, Vanja Pekovic-Vaughan, Pietro Mesirca, Matteo Mangoni, Halina Dobrzynski, Alicia D’Souza, Mark R. Boyett

**Affiliations:** School of Medicine and Dentistry, University of Lancashire, Preston, UK; School of Medicine, American University of the Caribbean, Cupecoy, St. Maarten; Division of Medical Sciences, University of Chester, Chester, UK; National Heart and Lung Institute, Imperial College London, London, UK; Institut de Génomique Fonctionnelle, Université de Montpellier, CNRS, INSERM, Montpellier, France; Division of Cardiovascular Sciences, University of Manchester, Manchester, UK; Erasmus Medical Center, Dr. Molewaterplein 40, 3015 GD Rotterdam, The Netherlands; Institute of Life Course and Medical Sciences, Faculty of Health and Life Sciences, University of Liverpool, Liverpool, UK; College of Medicine & Health Sciences, United Arab Emirates University, Al Ain, Abu Dhabi, United Arab Emirates; School of Biological Sciences, University of Leeds, Leeds, UK; Faculty of Life Sciences, University of Bradford, Bradford, UK; School of Physiology, Pharmacology & Neuroscience, University of Bristol, Bristol, UK; Department of Biomedical Sciences, University of Copenhagen, Copenhagen, Denmark; Morinomiya University of Medical Sciences, Osaka, Japan; Department of Anatomy, Jagiellonian University Medical College, Kopernika 12, 31-034, Krakow, Poland

**Keywords:** cardiac circadian rhythms, atrioventricular node, PR interval, ion channels, ionic current, metabolism, contractile proteins, extracellular matrix

## Abstract

The atrioventricular (AV) node shows a circadian rhythm – during sleep the PR interval is longer (because conduction velocity is decreased) and AV block is more common. We show the rhythm is similar in mouse and human, and using mice we have investigated the cause. There was still a rhythm in *Kcnj5* knockout mice ruling out a major role of vagal tone (the vagus regulates the node via *Kcnj5*). Instead, we show the presence of a functioning circadian clock in the node (e.g. by detection of rhythmic clock transcripts). There was ∼43-52% reduction in the rhythm of the PR interval on inhibition of the clock in a *Bmal1* knockout mouse. There was a significant rhythm in >300 intracellular signalling molecules presumably driven by the clock and perhaps extrinsic signals. Downstream of the clock and signalling molecules, there was a rhythm in many transcripts and proteins (of 4,186 proteins, ∼15% had a rhythm), responsible for example for contractile machinery, metabolism, extracellular matrix, and importantly ion channels. Ultimately, the PR interval is determined by ion channels, and there were rhythms in ion channels and ionic currents. A key determinant of conduction velocity is the upstroke velocity of the Ca^2+^-dependent nodal action potential and this was significantly reduced during sleep – this can be explained by reduced expression of Ca^2+^ channel subunits and a smaller L-type Ca^2+^ current at this time and this was confirmed by biophysically-detailed computer modelling. In conclusion, there is day-night molecular remodelling of the AV node and this is largely responsible for the circadian rhythm in the PR interval.

## INTRODUCTION

Many if not all physiological systems, including the heart, show a day-night rhythm tailoring the systems to the different demands on the organism during the day and night.^1^ Here, day-night rhythms are referred to as circadian because the term is commonly used, whereas strictly a day-night rhythm should only be considered circadian if it persists in constant darkness. Many aspects of the heart show a circadian rhythm including electrical activity, contractility, metabolism and intracellular signalling^1^ and it is likely that other aspects of the heart also show a circadian rhythm. In terms of electrical activity, there is known to be a circadian rhythm in heart rate, the PR interval, the duration of the QRS complex, the QT interval and the susceptibility of the heart to different arrhythmias.^2–6^ During the awake period (day in the case of the human and night in the case of the nocturnal mouse) the heart rate is higher to meet the greater demands on the circulation, and conduction through the atrioventricular (AV) node and His-Purkinje system is faster (explaining the shortened PR interval and QRS duration at this time) presumably because less time is available between heart beats.^2^ The circadian rhythm of the PR interval is of pathological importance: the slowing of AV node conduction during the sleep period means that the heart is more vulnerable to AV block. In the human, it is known that AV block primarily occurs at night.^7,8^ Park *et al*.^9^ have recently published a study of “nocturnal atrioventricular block” and reported that 21% (14/67) of patients required a pacemaker implantation during a median follow-up period of 2.4 years. In the rat, it has also been shown that AV block primarily occurs during the sleep period.^10^ For approaching 100 years, the circadian rhythm in the heart rate has been attributed to a day-night rhythm in autonomic tone (principally vagal tone) to the heart.^11,12^ The circadian rhythm in the PR interval has also been attributed to this.^13^ All circadian rhythms are ultimately the result of a master circadian clock in the suprachiasmatic nucleus in the hypothalamus, but local circadian clocks (under the control of the master clock) are also present in peripheral tissues.^14^ We have recently shown that there is a functioning local circadian clock in the mammalian sinus node, the pacemaker of the heart, and the circadian rhythm in the heart rate persists after complete autonomic blockade or in the isolated denervated sinus node.^4^ Therefore, the circadian rhythm in the heart rate is in part at least intrinsic to the mammalian heart. Interestingly, there is also a circadian rhythm in the heart rate, which is intrinsic to the heart, in teleosts and crustaceans.^15,16^ We showed that in the mammalian sinus node there is a circadian rhythm in the expression of key pacemaker ion channels, including the funny channel, HCN4, and block of HCN4 abolishes the circadian rhythm in the heart rate.^4^ In contrast to this, based on the effects of pharmacological autonomic blockade, Hayter et al.^17^ concluded that the circadian rhythm in the PR interval is the exclusively the result of changes in autonomic tone.

We now report a functioning local circadian clock in the AV node as well as a circadian rhythm in the expression of key ion channel transcripts, proteins and ionic currents. The circadian rhythm in the PR interval is reduced by ∼50% on incapacitating the local circadian clock. Finally, there is a circadian rhythm in many other aspects of the AV node: contractile proteins, metabolism, extracellular matrix, signalling pathways, protein kinases, transcription factors and micro-RNAs.

## METHODS

Continuous three-channel bipolar Holter ECG recordings from 10 human subjects were analysed and the PR interval (from the onset of the P-wave to the end of the QRS complex) as well as the RR interval were measured. The majority of other experiments were carried out on wild-type or genetically-modified mice with national and local ethical approval. The electrocardiogram (ECG) was measured *in vivo* in the conscious freely moving mouse using telemetry. Intracellular action potentials were recorded from the AV node with sharp microelectrodes. Transcripts and micro-RNAs in the AV node were measured using quantitative PCR (qPCR), and proteins in the AV node were measured using proteomics. Functioning of the circadian clock in the isolated AV node was measured using bioluminescent PER1 and PER2 reporters. Ionic currents in isolated AV node myocytes were measured with the patch clamp technique. Biophysically-detailed computer modelling was used to assess the importance of the observed circadian rhythm in the L-type Ca^2+^ current. Time is normally reported as zeitgeber time (ZT) in hours; ZT0 corresponds to lights-on at the beginning of the light period (sleep period) and ZT12 corresponds to lights-off at the beginning of the dark period (active period). Data are reported as means±SEMs; statistical tests used to assess differences are given in the text. In the text, unique gene names are generally given (in italics with an initial capital) regardless of whether the gene, transcript or protein is being referred to. Because gene names can be unfamiliar, the widely used protein names (in an upright font and, commonly, but not necessarily, in capitals) may be given. If the gene name is most commonly used, the protein name is omitted for brevity. Full details of the methods used are available in the Data Supplement.

## RESULTS

### Circadian rhythm of the AV node and role of the acetylcholine (ACh)-activated K^+^ channel

Continuous three-channel bipolar Holter ECG recordings were obtained from 10 human subjects. The central finding was a circadian rhythm in the PR interval and a consistent nocturnal prolongation of the PR interval: the PR interval increased at night in 10/10 patients; the patient-level median difference was +19.5 [14.6–36.1] ms (range +2.0 to +64.5 ms), with both the exact sign test and Wilcoxon signed-rank test yielding P=0.0020. There was also a circadian rhythm in the RR interval: the RR interval increased at night in 8/10 patients. The median night-day change was +164.1 [141.6– 226.6] ms; the Wilcoxon signed-rank test was P=0.0098, whereas the exact sign test was P=0.1094, reflecting two patients with small decreases in the RR interval at night rather than increases seen in eight patients. The time course of the circadian rhythm in the median (and interquartile range) PR interval for the 10 subjects is shown in Fig. 1A (corresponding data from the individual subjects are shown in Supplementary Fig. S1A). There was marked decline in the PR interval during the morning. Although the PR interval increased subsequently, it remained below the 24 h average (dotted line in Fig. 1A) during the day. Fig. 1A also shows the time course of the circadian rhythm in the RR interval and the time course is broadly similar to that of the PR interval. As discussed in the Introduction, both the night-time increases in the PR and RR intervals have been attributed to a night-time increase in vagal tone^11–13,17^ and the broadly similar time courses of the circadian rhythms (Fig. 1A) are arguably consistent with this hypothesis. However, arguably inconsistent with the hypothesis, the magnitude of the nocturnal PR interval change was not significantly associated with the magnitude of the nocturnal RR change across patients (Fig. S1B; Spearman ρ=0.351, P=0.321): patients with the largest nocturnal PR interval change were not necessarily those with the largest RR change. For example, subject 1395918 had a slightly shorter RR at night (−23.4 ms) while the PR interval still increased (+19.5 ms); subject 2353552 showed the largest nocturnal PR interval increase (+64.5 ms) despite only a moderate RR increase (+144.5 ms).

**Fig. 1.**
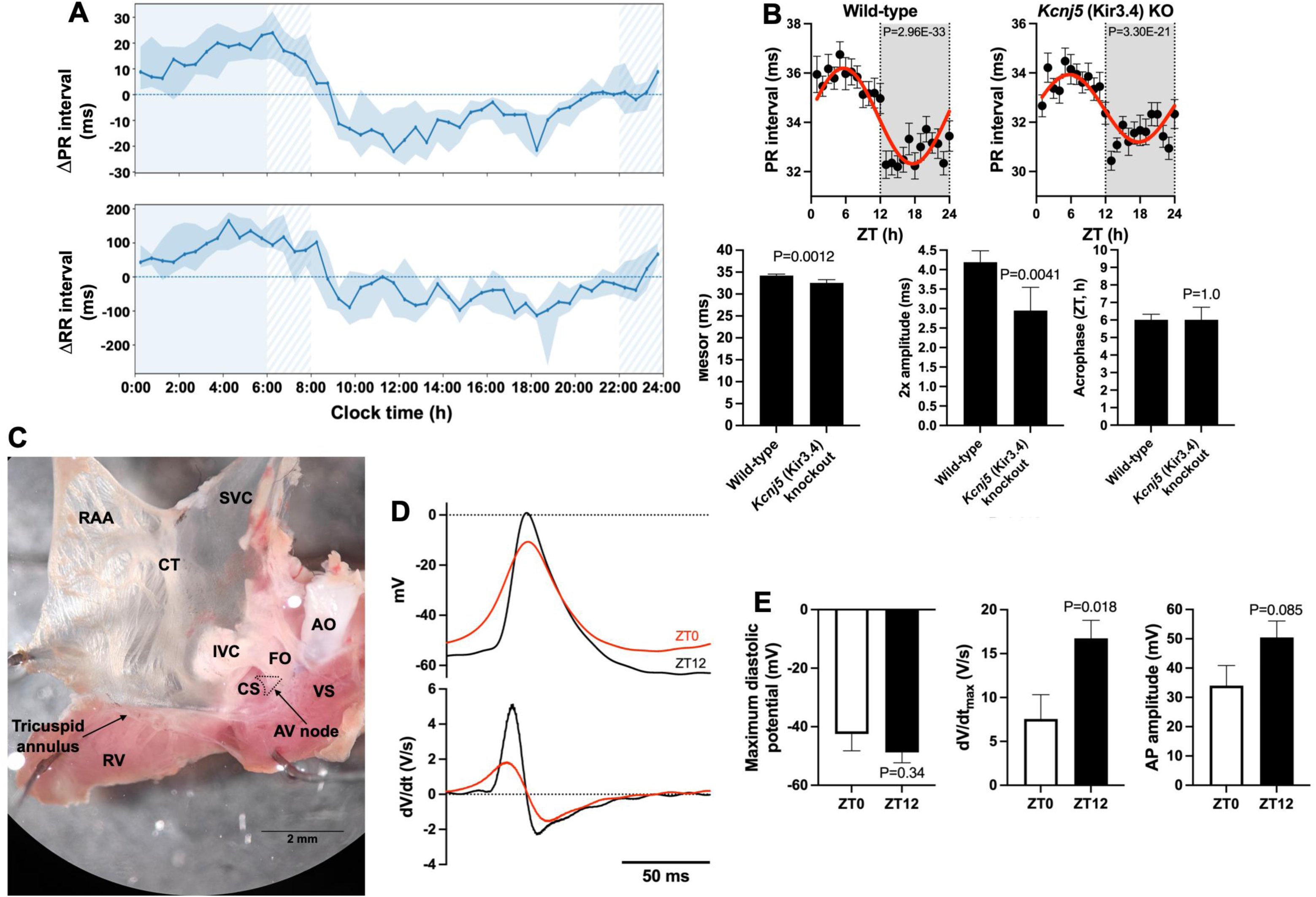
Circadian rhythm in the PR interval in the human and mouse. **A,** mean (and interquartile range – shaded) of PR and RR intervals of 10 human subjects over 24 h. To account for differences in intervals among subjects, for each subject, the intervals were calculated with reference to the median over the 24 h. On the time axis, the shaded area indicates night and the hatched area indicates the transition between night and day. **B**, circadian rhythm in the PR interval in wild-type and *Kcnj5* (Kir3.4) mice. Top panel, mean±SEM PR interval at 1 h intervals measured using telemetry from conscious freely moving wild-type (n=21) and *Kcnj5* (Kir3.4) knockout (n=23) mice over 24 h. Hours of darkness shaded; P value from cosinor analysis (of pooled data from all mice) for significance of a day-night rhythm given; data fitted with a cosine wave (shown as a red line) by a least squares fitting method. Bottom panel, mean±SEM mesor, 2xamplitude and acrophase of the circadian rhythm in the PR interval in the wild-type and *Kcnj5* (Kir3.4) knockout mice (based on cosinor analysis of individual mice). By convention, amplitude is the difference between the mesor and the peak (or nadir); 2x amplitude is therefore the difference between the peak and nadir. The difference between the wild-type and knockout mice was tested using an unpaired t test and the P value is shown. **C**, endocardial surface of a mouse isolated right atrial preparation. AO, aorta; CS, coronary sinus; CT, crista terminalis; FO, fossa ovalis; IVC, inferior vena cava; RAA, right atrial appendage; RV, right ventricle; SVC, superior vena cava; VS, ventricular septum. **D**, examples of intracellular APs recorded from the isolated AV node of the mouse at ZT0 and 12. **E**, mean (±SEM) maximum diastolic potential, maximum upstroke velocity of the AP, and amplitude of the AP at ZT0 (n=6) and ZT12 (n=8). P values shown (two-tailed unpaired t test).

Experiments on mice provided further evidence against the vagal tone hypothesis. Fig. 1B shows the mean PR interval in conscious freely moving wild-type mice (n=21) as measured by telemetry over 24 h. In this study, cosinor analysis^18,19^ was used to determine whether a measurement shows a significant circadian rhythm; if significant (P<0.05), in figures the data are fitted with a cosine wave in red. The mean PR interval in mice shows a significant rhythm and, as in the human, the PR interval was shortest (i.e. AV node conduction was fastest) during the awake period (night in case of mouse). The action of the vagus on the AV node has been shown to be the result of the activation of the ACh-activated K^+^ current carried by the heteromeric ACh-activated K^+^ channel made of up of Kir3.1 (also known as GIRK1; gene, *Kcnj3*) and Kir3.4 (also known as GIRK4; gene, *Kcnj5*) subunits: Drici *et al*.^20^ reported that tertiapin-Q, a peptide which selectively blocks the ACh-activated K^+^ channel, prevents dose-dependent ACh-induced AV blocks. Contrary to the current explanation of the circadian rhythm in the PR interval, Fig. 1A shows that a significant circadian rhythm remained in Kir3.4 knockout mice (n=23), although cosinor analysis showed that the amplitude of the circadian rhythm was reduced to 70.4% of that in the wild-type mice (Fig. 1B). Cosinor analysis also showed that the mesor (mean PR interval) and acrophase (time of peak PR interval) were little or not affected in the Kir3.4 knockout mice (Fig. 1B). This suggests that, although vagal tone may have a minor role (although not necessarily so – see Supplementary Discussion), other factors are also involved. If vagal tone is not the explanation, what is responsible for the circadian rhythm in the PR interval? Intracellular action potentials (APs) were recorded from the area of the AV node in isolated right atrial preparations (Fig. 1C). As expected, the AP morphology was heterogeneous,^21^ and APs were divided into nodal, transitional and atrial types (Fig. S2A). Fig. 1D shows example nodal APs and Fig. 1E the mean value of the maximum diastolic potential, the maximum upstroke velocity of the AP (dV/dt_max_), and the AP amplitude at ZT0 and ZT12 for nodal APs (see Fig. S2B for AP duration). At ZT12, the start of the active period, there was a significant increase in dV/dt_max_ and a similar trend in the AP amplitude (Fig. 1D,E). dV/dt_max_ and AP amplitude are key determinants of the conduction velocity, and increases at ZT12 are expected to increase the conduction velocity (and therefore explain the shortening of the PR interval during the active period). The AV node AP is Ca^2+^-dependent, unlike the Na^+^-dependent AP of the working myocardium. This explanation of the circadian rhythm in the PR interval is further explored below.

### Evidence of a functioning circadian clock in the AV node

Local circadian clocks are known to be present in many and perhaps all peripheral tissues. Using qPCR, the expression of 96 transcripts including key circadian clock transcripts was measured in the mouse AV node at four time points over 24 h. Expression was measured at ZT0, 6, 12 and 18. Fig. 2A shows a schematic diagram of the circadian clock and the mean±SEM expression of circadian clock transcripts over 24 h. There was a significant circadian rhythm in all transcripts with one exception. We have previously reported almost identical expression of circadian clock transcripts in the mouse sinus node (but data were obtained using RNAseq rather than qPCR):^3^ Fig. 2B shows the acrophase (ZT at which peak expression occurred) and the amplitude (here defined as the day-night difference in expression as a percentage of the maximum expression) of the circadian rhythm of the circadian clock transcripts in the mouse AV node plotted against those in the mouse sinus node. In Fig. 2B each point corresponds to a particular transcript and the red lines have a slope of one (lines of identity); the data fall along the line of identity in both cases. PER2 is an important clock gene (Fig. 2A) and expression of PER2 was measured by recording bioluminescence from mice expressing a PER2 reporter (PER2::LUCIFERASE); recordings were made from the isolated AV node and also atrial muscle (as a positive control). A representative bioluminescence trace from the AV node is shown in Fig. 2C (for comparison a representative trace from atrial muscle is shown in Fig. S3A). The AV node had a robust PER2::LUCIFERASE rhythm over five cycles with a period (peak-to-peak time) of 25.7 h and a projected acrophase of 12.6 h (Fig. S3A). This is comparable to the *Per2* transcript *in vivo*: *in vivo* the period of *Per2* was 24 h and the acrophase was 12.5 h (Fig. 2A). In Fig. 2C the circadian rhythm waned after five cycles and the rhythm was reinstated using 0.1 mM dexamethasone, a synthetic glucocorticoid, which is well known to induce, synchronise, and phase-shift circadian gene expression in peripheral tissues.^22^ This led to an increase in amplitude compared to initial values and phase resetting (Figs. 2C and S3A). Using a mouse expressing a PER1 reporter (PER1::LUCIFERASE), expression of PER1 was also measured (Fig. S3B); the data are similar to the data for PER2 (Fig. 2C). These data confirm the presence of a functioning circadian clock in the AV node. The involvement of the local circadian clock in the AV node was tested independently in two centres by recording the circadian rhythm in the PR interval in cardiomyocyte-specific *Bmal1* knockout mice using telemetry. *Bmal1* is a key circadian clock gene (Fig. 2A) and knockout of *Bmal1* is known to incapacitate the circadian clock.^23^ Fig. 2D shows data from the University of Copenhagen from nine wild-type and 11 CBK (cardiomyocyte-specific *Bmal1* knockout) mice. A significant circadian rhythm in the PR interval persisted in the *Bmal1* knockout mice (Fig. 2D), but it was significantly reduced in amplitude to 68.7% (Fig. S4A). Similar data were obtained at the University of Manchester (Fig. S4B,C). This shows that the local circadian clock in the AV node is involved in the circadian rhythm in the PR interval.

**Fig. 2.**
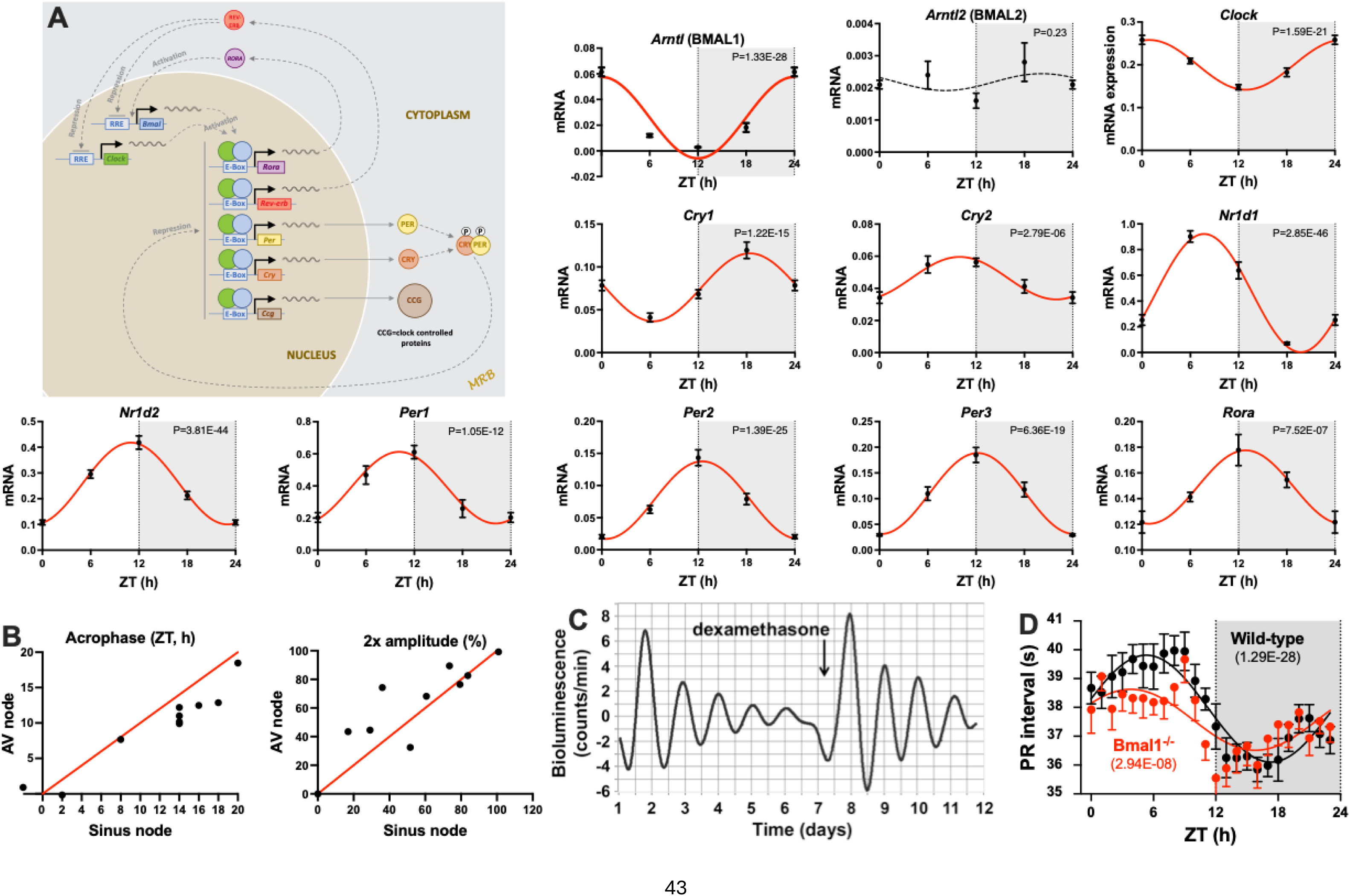
Circadian clock in the AV node. **A,** abundance of circadian clock transcripts measured at four time points over 24 h. The inset shows a schematic diagram of the circadian clock from Wang *et al*.^3^ In this and similar figures: gene name given (common name may also be given); mean±SEM transcript, protein or micro-RNA abundance shown at four ZT time points over 24 h (24 h data repeat of 0 h data); hours of darkness shaded; P value from cosinor analysis for significance of a day-night rhythm given; transcripts, proteins or micro-RNAs with a significant circadian rhythm (P<0.05) fitted with a cosine wave (shown as a red line) by a least squares fitting method; non-significant transcripts, proteins or micro-RNAs may be fitted with a cosine wave shown as a black dashed line. In all figures showing data for transcripts, as in this case, n=6-11 mice. **B**, acrophase and 2x amplitude of circadian clock transcripts in the AV node in this study (shown in A) plotted against the acrophase and 2x amplitude of the same transcripts in the sinus node (data from Wang *et al*.^3^). The acrophase is the ZT at peak expression. By convention, the amplitude is the difference between the mesor and the peak (or nadir) and therefore 2x amplitude is the day-night variation (here expressed as a percentage of the maximum). Lines of identity are shown in red. **C**, evidence of a functioning circadian clock in the AV node. PER1 activity (reported by luciferase bioluminescence) in the isolated AV node from a Per1::LUC background shown. PER1 activity varied in a circadian manner. When the circadian rhythm had run down, it was reinstated by 0.1 mM dexamethasone, a synthetic corticosteroid. The bioluminescence trace is baseline-subtracted. **D,** Knockout of the circadian clock in the heart blunts the circadian rhythm in the PR interval. Mean±SEM PR interval measured using telemetry from nine wild-type and 11 cardiomyocyte-specific *Bmal1* knockout (CBK) mice shown (data taken from the study of Gottlieb et al.^98^).

### Circadian rhythm in ion channel transcripts

332 ion channels are listed in the HGNC database. Expression of 44 transcripts for key cardiac ion channels known to be important in AV node pacemaking and conduction were also measured using qPCR (Supplementary Table S1). 22 (50%) showed significant circadian rhythms (Figs. 3, 4 and S5; Supplementary Table S1). HCN channels are known to be expressed in the AV node and be important for both pacemaking and conduction.^24,25^ The expression of three *Hcn* transcripts was measured and expression was *Hcn1*>*Hcn4*>*Hcn2* (Fig. 3). The two most abundant isoforms, *Hcn1* and *Hcn4*, showed significant circadian rhythms and peak expression occurred during the active period (Fig. 3). A Cl^-^ channel, CLC-2 (*Clcn2*), behaves in a functionally similar way to the HCN channels and has been shown to play a role in sinus node pacemaking^26^ – *Clcn2* showed a significant circadian rhythm and peak expression occurred at the start of the active period (Fig. 3). Although the cardiac Na^+^ channel, Nav1.5 (*Scn5a*), is absent in the compact AV node,^27^ it is present in the peripheral regions^28^ and is known to be important for AV node conduction.^29–31^ Both *Scn5a* and the transcript for another Na^+^ channel, *Scn1a* (Nav1.1), showed a significant circadian rhythm and peaked at the start of the active period (Fig. 3). The L-type Ca^2+^ channel carried by Cav1.2 and Cav1.3 channels is responsible for the upstroke of the AV node action potential in the absence of the Na^+^ current, *I*_Na_.^32^ *Cacna1c* (Cav1.2) and *Cacna1d* (Cav1.3) showed significant circadian rhythms and peaked during the active period (Fig. 3). Evidence suggests that the accessory L-type Ca^2+^ channel subunits, Cavα2δ1 (*Cacna2d1*) and Cavα2δ2 (*Cacna2d2*), facilitate Ca^2+^ entry via the L-type Ca^2+^ channel – for example, deletion of the Cavα2δ1 subunit decreases the L-type Ca^2+^ current, *I*_Ca,L_.^33,34^ It is therefore potentially important that both *Cacna2d1* and *Cacna2d2* approached significance (0.05<P<0.1) in terms of a circadian rhythm and peaked at approximately the same time as *Cacna1c* and *Cacna1d* (Fig. 3). Cav3.1 (*Cacna1g*) plays an important role in AV node conduction and knockout of Cav3.1 slows AV node conduction^35^ and it is therefore surprising that *Cacna1g* showed a significant circadian rhythm and reached a nadir at the start of the active period (Fig. 3).

**Fig. 3.**
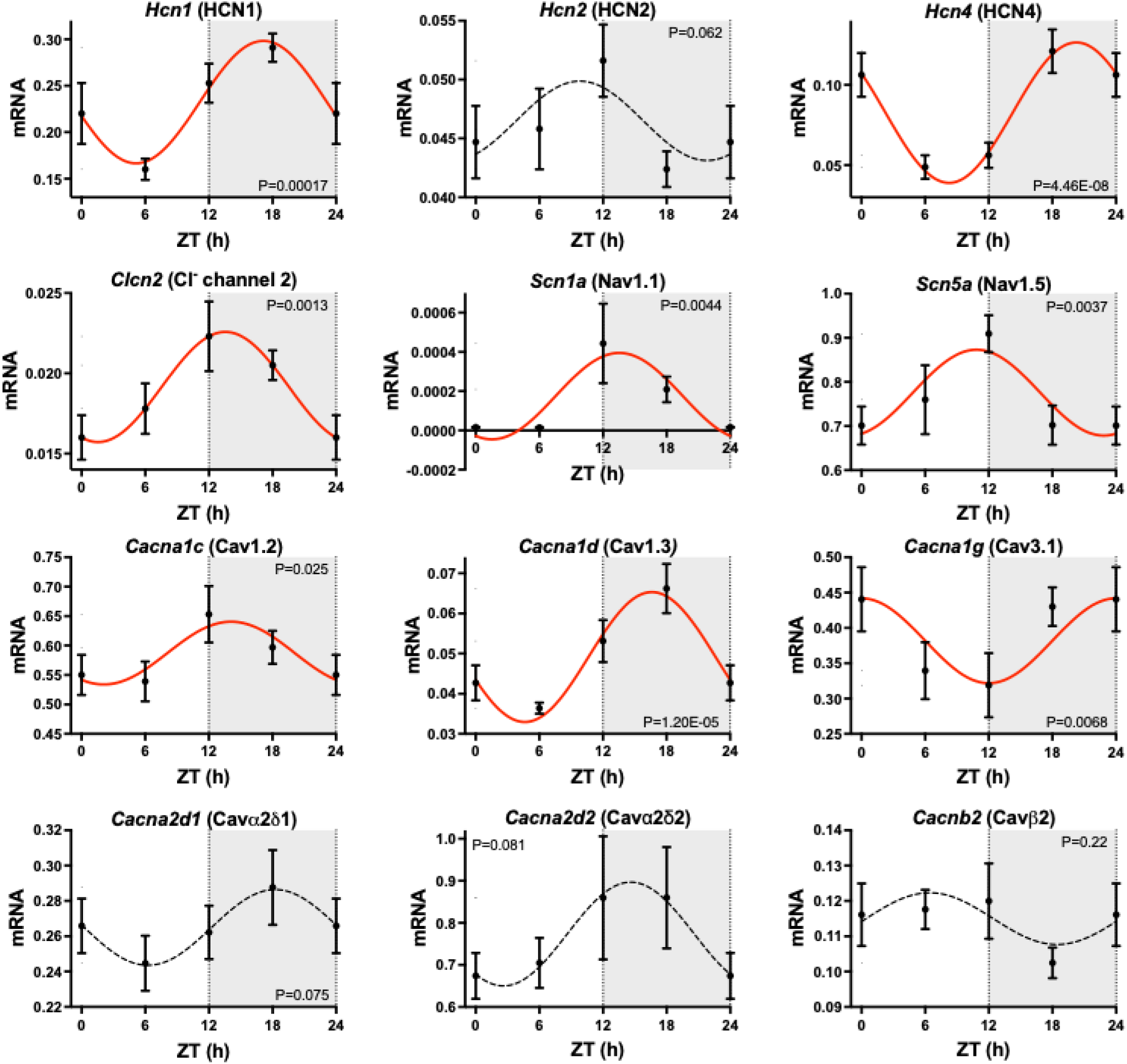
Circadian rhythm in ion channel transcripts responsible for inward currents in the AV node. Data for HCN, Cl^-^, Na^+^ and Ca^2+^ channels shown. See Fig. 2 legend for a general description of the figure.

The transient outward K^+^ current (*I*_to_) contributes to action potential repolarization. Some AV node myocytes are reported to express *I*_to_,^36^ and three transcripts, *Kcna4* (Kv1.4), *Kcnd2* (Kv4.2) and *Kcnd3* (Kv4.3), for transient outward K^+^ channel subunits were detected (Fig. 4B). *Kcna4* (Kv1.4) and *Kcnd2* (Kv4.2) showed significant circadian rhythms but were out of phase with each other and one peaked during the active period, whereas the other reached a nadir during the active period (Fig. 4B). Mice lacking both the fast (Kv4-dependent) and the slow (Kv1.4-dependent) components of *I_to_* display second degree AV blocks.^37^ Expression of transient outward K^+^ channel transcripts was low compared to expression of delayed rectifier K^+^ channel transcripts (Fig. 4C). Delayed rectifier K^+^ current contributes to action potential repolarization and it is likely that it helps set the maximum diastolic potential immediately after the action potential as it does in the sinus node.^38^ Transcripts for three delayed rectifier K^+^ channels, *Kcna5* (Kv1.5), *Kcnh2* (ERG) and *Kcnq1* (KvLQT1), responsible for the ultra-rapid, fast and slow delayed rectifier K^+^ currents (*I*_K,ur_, *I*_K,r_ and *I*_K,s_), were detected (Fig. 4C). All three transcripts including *Kcnh2* (ERG), the most abundant, showed significant circadian rhythms and peaked at the beginning of the active period (Fig. 4C). The background inward rectifier K^+^ current (*I*_K,1_) helps to set the membrane potential during diastole. Two transcripts responsible for *I*_K,1_ were measured, *Kcnj2* (Kir2.1) and *Kcnj12* (Kir2.2), and both showed significant circadian rhythms and peaked during the active period (Fig. 4D). As already discussed, vagal control of AV node conduction and the PR interval involves the activation of the ACh-activated K^+^ current (*I*_K,ACh_)^39^ and the transcripts responsible for the two K^+^ channel subunits making up the ACh-activated K^+^ channel, *Kcnj3* (Kir3.1) and *Kcnj5* (Kir3.4), both showed significant circadian rhythms and were greater during the active period (Fig. 4E); the implications of this for the circadian rhythm in the PR interval are considered in the Supplementary Discussion. SK channels play an important role in the AV node – downregulation, knockout or overexpression of SK2 (most abundant SK channel in AV node; Fig. 4F) results in a prolongation of the PR interval, complete AV block or a shortening of the PR interval, respectively.^40^ *Kcnn3* responsible for SK3 showed a significant circadian rhythm and *Kcnn1* and *Kcnn2* responsible for SK1 and SK2 approached significance (0.05<P<0.1) in terms of a circadian rhythm; expression of all three transcripts peaked during the active period, but at different times (Fig. 4F).

**Fig. 4.**
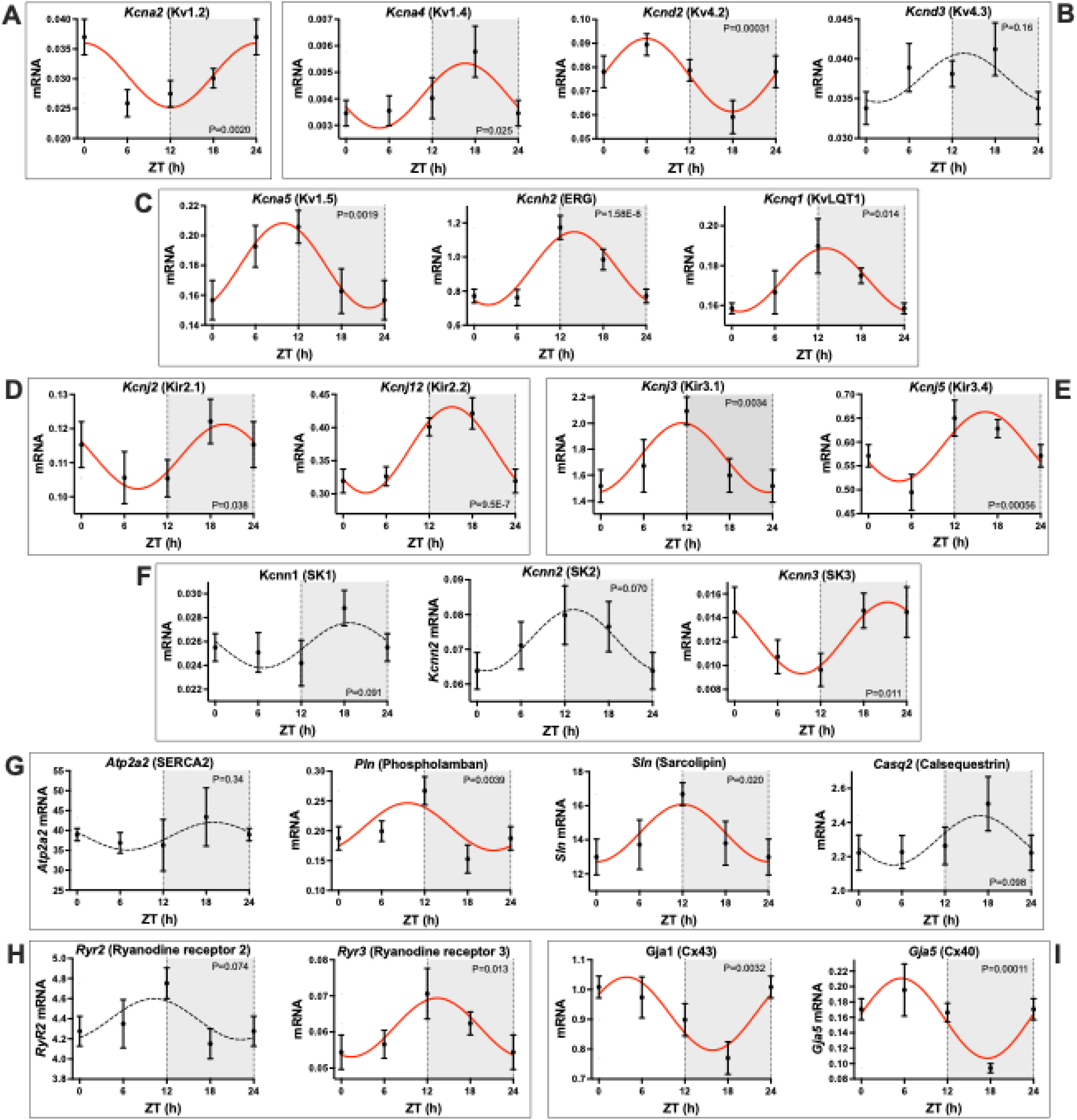
Circadian rhythm in ion channel transcripts responsible for outward currents and other transcripts related to electrical activity in the AV node. **A**, *Kcna2* (Kv1.2). **B**, transient outward K^+^ channel transcripts. **C**, delayed rectifier K^+^ channel transcripts. **D**, background inward rectifier K^+^ channel transcripts. **E**, ACh-activated K^+^ channel transcripts. **F**, small-conductance Ca^2+^-activated K^+^ channel transcripts. **G**, Ca^2+^ clock transcripts. **H**, ryanodine receptor channel transcripts. **I**, gap junction channel transcripts. See Fig. 2 legend for a general description of the figure.

The so-called Ca^2+^-clock is known to be an important pacemaker mechanism in the sinus node, but there is also evidence that it is involved in AV node conduction – ryanodine slows AV node conduction in the rat.^41^ Various transcripts related to the Ca^2+^ clock, *Pln* (phospholamban), *Sln* (sarcolipin), *Casq2* (calsequestrin 2) as well as *Ryr2* (ryanodine receptor 2), showed a significant circadian rhythm (or a trend – 0.05<P<0.1) and peaked sometime during the active period (Fig. 4G,H). Interestingly, *Ryr3* (ryanodine receptor 3) has been shown to be expressed in the sinus node and right atrial myocytes;^42^ *Ryr3* was detected in the AV node and showed a significant circadian rhythm similar to the trend shown by *Ryr2* (Fig. 4H). Gap junctions are responsible for electrical coupling between myocytes and are important determinants of the conduction velocity.^43^ Transcripts for the two most abundant gap junction channels, *Gja1* and *Gja5* responsible for Cx43 and Cx40, both showed significant circadian rhythms but unexpectedly reached a nadir during the active period (Fig. 4I).

### Circadian rhythm in ion channel proteins

Protein expression does not necessarily follow transcript expression, and proteomics and mass spectrometry were used to measure protein expression in the mouse AV node at four time points over 24 h (at ZT0, 6, 12 and 18). 4,186 proteins were detected – it is likely that this is not the whole proteome (see Supplementary Discussion for consideration of the sensitivity and resolution of proteomics). Of the 4,186 proteins, 616 (14.7%) showed a significant circadian rhythm. Of the list of 332 ion channels in mammals, 39 ion channel proteins were detected – this may be limited in part due to the sensitivity of proteomics, because some ion channels known to be present in the mouse AV node and detected by qPCR were not detected at the protein level. Of the 39 ion channel proteins detected, seven (∼18%) showed a significant (P<0.05) circadian rhythm and a further eight (∼21%) showed a trend towards significance (0.05<P<0.1). Fig. 5 shows protein expression of ion channels with a significant circadian rhythm as well as some other key ion channels. In the case of *Scn5a* (Nav1.5), *Cacna1c* (Cav1.2), C*acna2d1* (Cavα2δ1), C*acna2d2* (Cavα2δ2), *Kcnh2* (ERG), *Kcnj3* (Kir3.1), *Kcnj5* (Kir3.4), *Ryr2* (ryanodine receptor 2), *Ryr3* (ryanodine receptor 3) and *Gja5* (Cx40), expression at the protein level was consistent (qualitatively if not quantitatively) with expression at the transcript level (cf. Figs. 3- 5). However, in the case of HCN4 (*Hcn4*), there is a striking mismatch: whereas in the case of the transcript there was a marked and highly significant circadian rhythm, in the case of the protein there appeared to be no rhythm (cf. Figs. 3 and 5). In addition, there is a mismatch in the expression of transcript and protein in the case of Cx43 (*Gja1*) (cf. Figs. 4 and 5). Data for ion channel subunits responsible for the ATP-sensitive K^+^ channel are shown in Fig. S5; only *Abcc8* (SUR1) showed a significant rhythm.

**Fig. 5.**
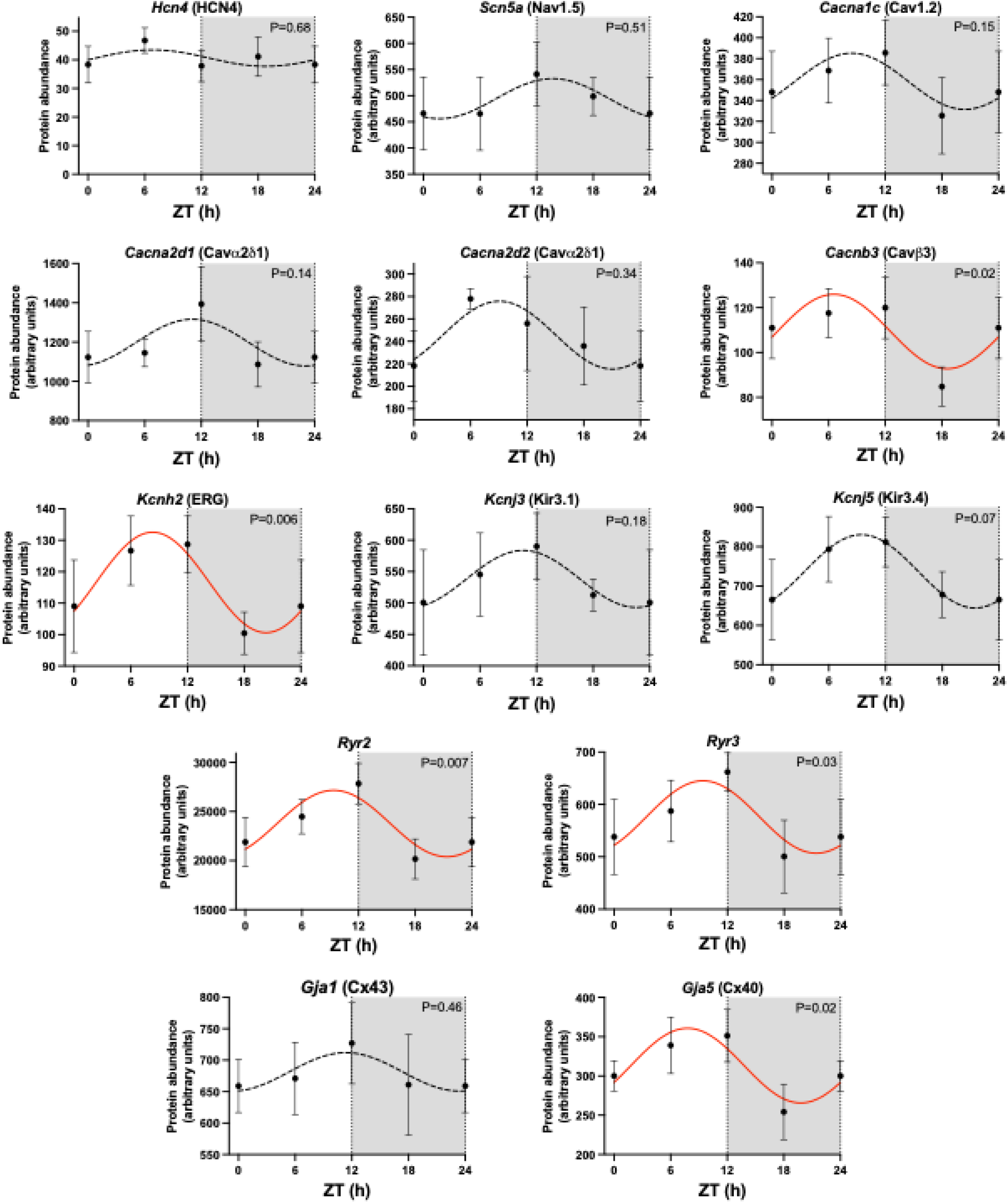
Circadian rhythm in ion channel proteins in the AV node. Data for HCN4, the cardiac Na^+^ channel, Ca^2+^ channels, K^+^ channels, ryanodine receptor channels and gap junction channels shown. See Fig. 2 legend for a general description of the figure. In this and similar figures showing protein abundance, each point is for four (ZT0, 12 and 18) or five (ZT6) biological replicates, each replicate being from the AV nodes of two mice. Therefore, 8 or 10 mice contributed to each point.

### Circadian rhythm in ionic currents

Ionic currents were recorded at ZT0, 6, 12 and 18 from isolated AV node myocytes using the patch clamp technique (Fig. 6). There was no difference in the cell capacitance over 24 h (Fig. S6). *I*_Ca,L_ was recorded during depolarizing pulses from a holding potential of -55 mV (Fig. 6A). In the example shown in Fig. 6A, *I*_Ca,L_ was greater at ZT18 than at ZT6 and this is confirmed by the mean current density-voltage relationships in Fig. 6B. Fig. 6C shows that there was a circadian rhythm in *I*_Ca,L_ density, which peaked at ZT18. *I*_Ca,L_ is known to play an important role in AV node conduction.^44^ Biophysically-detailed computer models of the AV node were used to test the role of ionic currents in the circadian rhythm in the AV node conduction velocity and consequently the PR interval. Qualitatively similar results were obtained from two independent models and examples from both models will be given. Fig. 6D shows a result from the model of the rabbit AV node of Inada *et al*.^45^ to test the role of *I*_Ca,L_. The model consists of atrial myocytes proximally, then atrio-nodal (AN) and nodal (N) myocytes, and finally nodo-His (NH) myocytes distally; it also includes parallel fast and slow pathways.^45^ The atrial myocytes were stimulated and the action potential propagated through the AV node to the NH myocytes and the conduction time was measured. Fig. 6D shows that AV node conduction time is a function of the L-type Ca^2+^ channel conductance. The data in Fig. 6B shows that there was a ∼51% increase in *I*_Ca,L_ from ZT6 to ZT18 and Fig. 6D shows that such an increase in *I*_Ca,L_ will have a substantial impact. *I*_K,r_ was recorded during depolarizing pulses from a holding potential of -40 mV. The mean current density-voltage relationships in Fig. 6E show *I*_K,r_ density was lower at ZT12 than at ZT0 and Fig. 6F shows that there was a circadian rhythm in *I*_K,r_ density, which peaked at ZT0. Fig. 6G shows a result from the model of a mouse AV node myocyte of Bartolucci *et al*.^46^ to test the combined role of the circadian rhythms in *I*_Ca,L_ and *I*_K,r_. A one-dimensional string of AV node myocytes was constructed; the coupling between myocytes was set to achieve a conduction velocity of ∼3.2 cm/s, typical of the AV node,^43^ and the length of the string was set to achieve a conduction time to match the PR interval typically seen in experiments (e.g. Fig. 1B). Fig. 6G shows the conduction time at ZT0, 6, 12 and 18 (calculated based on conductances of *I*_Ca,L_ and *I*_K,r_ extracted from Fig. 6C,F at the four time points); the data show a circadian rhythm strikingly similar to the circadian rhythm in the PR interval in experiments (e.g. Fig. 1B). *I*_K,ACh_ was recorded using two protocols. First, current was recorded during ramp clamps from -100 to +40 mV from a holding potential of -80 mV before and after the application of 100 nM ACh (Fig. S7). *I*_K,ACh_ was calculated as the difference between the two at each potential and mean current density-voltage relationships are shown in Fig. 6H. As expected, *I*_K,ACh_ reversed at or near the K^+^ equilibrium potential, E_K_. At ZT18, *I*_K,ACh_ was more outward as compared to current at ZT0 positive to E_K_ and more inward negative to E_K_ (Fig. 6H) as expected of an increase in ionic conductance at ZT18. In Fig. 6I this is confirmed by the mean data at +35 mV positive to E_K_ and -100 mV negative to E_K_. Secondly, the membrane potential was held at -45 mV and 100 nM ACh was applied (Fig. 6J). Fig. 6J shows data at ZT6 and ZT18 and, consistent with the data in Fig. 6H, *I*_K,ACh_ was greater at ZT18. Fig. 6I,K show the density of *I*_K,ACh_ at ZT0/24, 6, 12 and 18. The data do not show a simple circadian rhythm of *I*_K,ACh_ as in the case of *I*_Ca,L_ and *I*_K,r_ – the data from both protocols show two peaks, the largest at ZT18 and a smaller one at ZT6. Using the one-dimensional model of the mouse AV node based on the model of Bartolucci *et al*.,^46^ the circadian rhythm in the conduction time was computed over 24 h based on the measured conductances of *I*_Ca,L_, *I*_K,r_ and *I*_K,ACh_. This time the circadian rhythm in the calculated conduction time bore no resemblance to the circadian rhythm in the PR interval in experiments – for example the conduction time was longest at ZT18 because of the large *I*_K,ACh_ at this time (data not shown). However, *I*_K,ACh_ depends on the ACh concentration at the AV node and it is not necessarily appropriate to include the measured channel conductances measured at an ACh concentration of 100 nM.

**Fig. 6.**
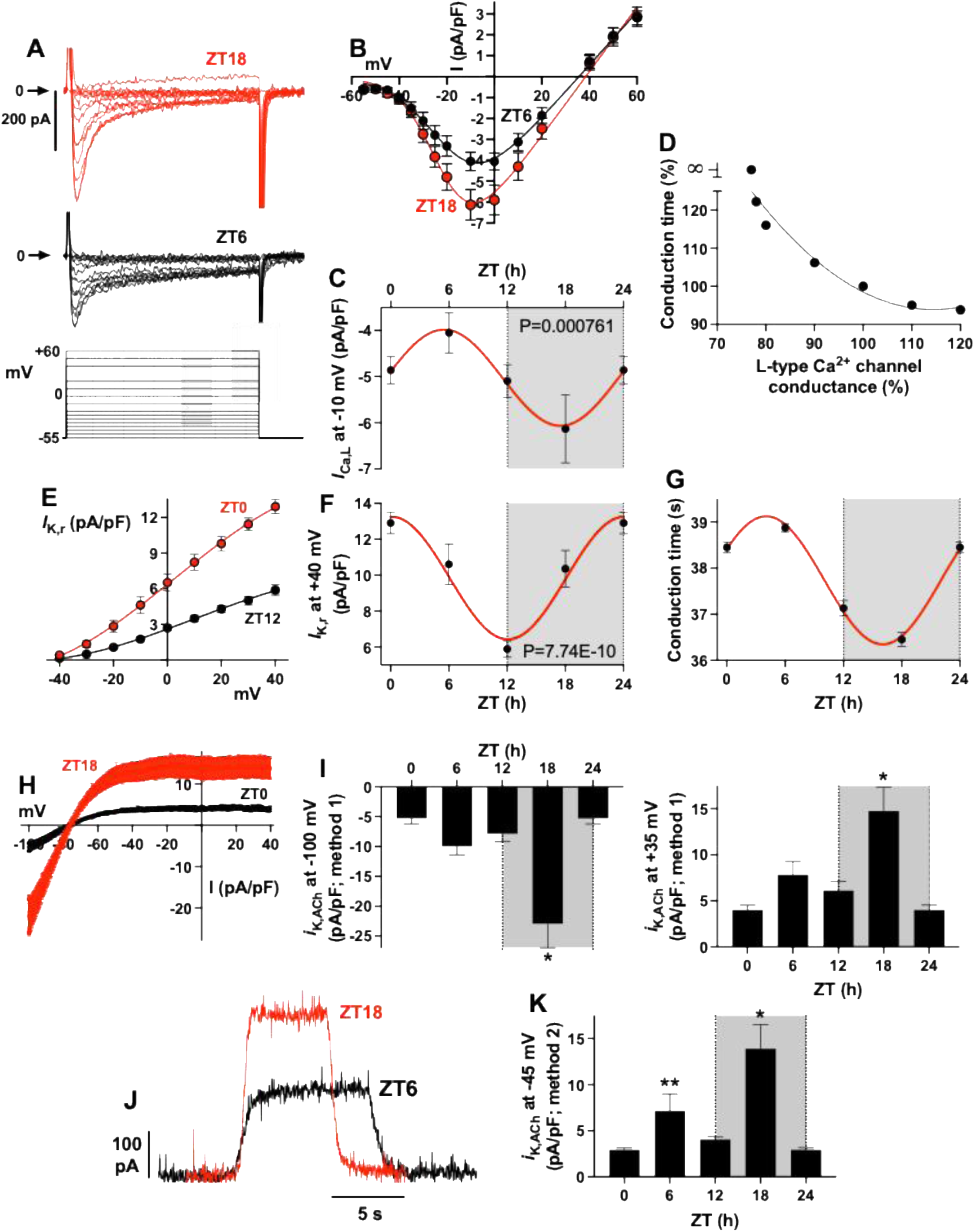
Circadian rhythm in *I*_Ca,L_, *I*_K,r_ and *I*_K,ACh_ in the AV node. **A**, families of recordings of *I*_Ca,L_ from isolated AV node myocytes at ZT6 and 18 (voltage clamp protocol shown below). **B**, current density-voltage relationships for *I*_Ca,L_ at ZT6 (n=17 myocytes) and 18 (n=14 myocytes). Means±SEM shown. **C**, circadian rhythm in *I*_Ca,L_ density at -10 mV. Means±SEM shown at ZT0/24, 6, 12 and 18 (n=14-21 myocytes at each time point). **D**, conduction time for the action potential in the one-dimensional model of the AV node of Inada *et al*.^45^ as a function of the L-type Ca^2+^ channel conductance, g_Ca,L_. The conduction time and g_Ca,L_ are expressed as a percentage of the conduction time and g_Ca,L_ in the normal configuration of the model. **E**, current density-voltage relationships for *I*_K,r_ at ZT0 (n=18 myocytes) and 12 (n=23 myocytes). Means±SEM shown. **F**, circadian rhythm in *I*_K,r_ density at +40 mV. Means±SEM shown at ZT0/24, 6, 12 and 18 (n=8-23 myocytes at each time point). **G**, calculated circadian rhythm in the conduction time though the AV node (taken to be 1.2 mm in length) in a one-dimensional model of a string of AV node myocytes modelled using the model of Bartolucci et al.^46^ Means±SEM shown at ZT0/24, 6, 12 and 18. **H**, current density-voltage relationships for *I*_K,ACh_ after application of 100 nM ACh during ramp clamps from -100 to +40 mV over 1.5 s at ZT0 (n=17 myocytes) and 18 (n=20 myocytes); data shown every ∼0.18 mV. See Fig. S7 for details of current measurement. **I**, mean±SEM *I*_K,ACh_ density at -100 mV (left panel) and +40 mV (right panel) at ZT0/24, 6, 12 and 18. Means±SEM shown (n=14-20 myocytes at each time point). *significantly different from data at ZT0, 6, 12 and 24; P<0.05; one-way ANOVA followed by Holm-Šídák’s multiple comparisons test. **J**, *I*_K,ACh_ at a holding potential of -45 mV on application of 100 nM ACh at ZT6 and 18. **K**, density of *I*_K,ACh_ at ZT0/24, 6, 12 and 18. Means±SEM shown (n=8-21 myocytes at each time point). *significantly different from the data at ZT0, 6, 12 and 24; **significantly different from the data at ZT0 and 24; P<0.01; one-way ANOVA followed by Holm-Šídák’s multiple comparisons test.

During a ramp clamp under control conditions in the absence of ACh, background current was recorded (Figs. S7A,B and S8A). Unlike *I*_K,ACh_, which is a strong inward rectifier K^+^ current and reverses at E_K_ (e.g. Fig. 6H), background current showed weak inward rectification as well as outward rectification and it reversed at ∼-30 mV and is likely to be the sum of various ionic currents (Figs. S7B and S8A). Fig. S8A shows that inward background current at ZT18 was greater than at ZT12. Once again, the current did not show a simple circadian rhythm – Fig. S8B shows background current at -100 mV at ZT0/24, 6, 12 and 18 and there appear to be two peaks, the largest at ZT18 and a smaller one at ZT6. The potential role of *I*_K,r_, *I*_K,ACh_ and background current in the circadian rhythm in the PR interval is considered in the Supplementary Discussion.

### HCN4/*I*_f_ paradox

HCN4 and the funny current, *I*_f_, already present a paradox: although evidence implicates *I*_f_ in AV node conduction^25,47^ (for example, knockout of underlying ion channel, HCN4, results in AV block and death in mouse^48,49^), computer modelling using biophysically-detailed models of the AV node AP fails to show a strong dependence of AV node conduction on *I*_f_.^50^ This study also presents paradoxical findings: *Hcn4* transcript showed a pronounced circadian rhythm (Fig. 3), whereas the HCN4 protein did not (Fig. 5) and neither did *I*_f_ (Fig. 7). *I*_f_ was recorded during pulses from a holding potential of -45 mV (Fig. 7A). Mean current density-voltage relationships in Fig. 7B shows that there was no discernible circadian rhythm in the density of *I*_f_ and this is confirmed by the mean density of *I*_f_ at -135 mV at ZT0/24, 6, 12 and 18 in Fig. 7C. The involvement of *Hcn4* in the circadian rhythm in AV node conduction was further explored using genetically modified mice. Two mouse lines, SND2A and SND2F, expressing a *conducting* HCN4 channel lacking the cAMP-dependent regulation of gating (mutant channels conduct *I*_f_, but current cannot be regulated by the cAMP-dependent pathway) and an AYA mouse line expressing a *non-conducting* dominant negative HCN4 mutant (in which GYG pore sequence is switched to AYA) were used. Consistent with earlier data,^25,47–49^ the effective knockout of HCN4 in the AYA mouse resulted in a marked slowing of conduction of AV node conduction and increase in the PR interval; there was relatively little effect in the SND2A and SND2F mouse lines (Fig. S9). There was still a circadian rhythm in the PR interval in the SND2A and SND2F mouse lines (data not shown). However, despite the lack of a circadian rhythm in HCN4 protein and *I*_f_ (Figs. 5 and 7A-C), the circadian rhythm in the PR interval was lost on effective knockout of HCN4 in the AYA mouse (Fig. 7D). How these conflicting data can possibly be reconciled are considered in the Discussion.

**Fig. 7.**
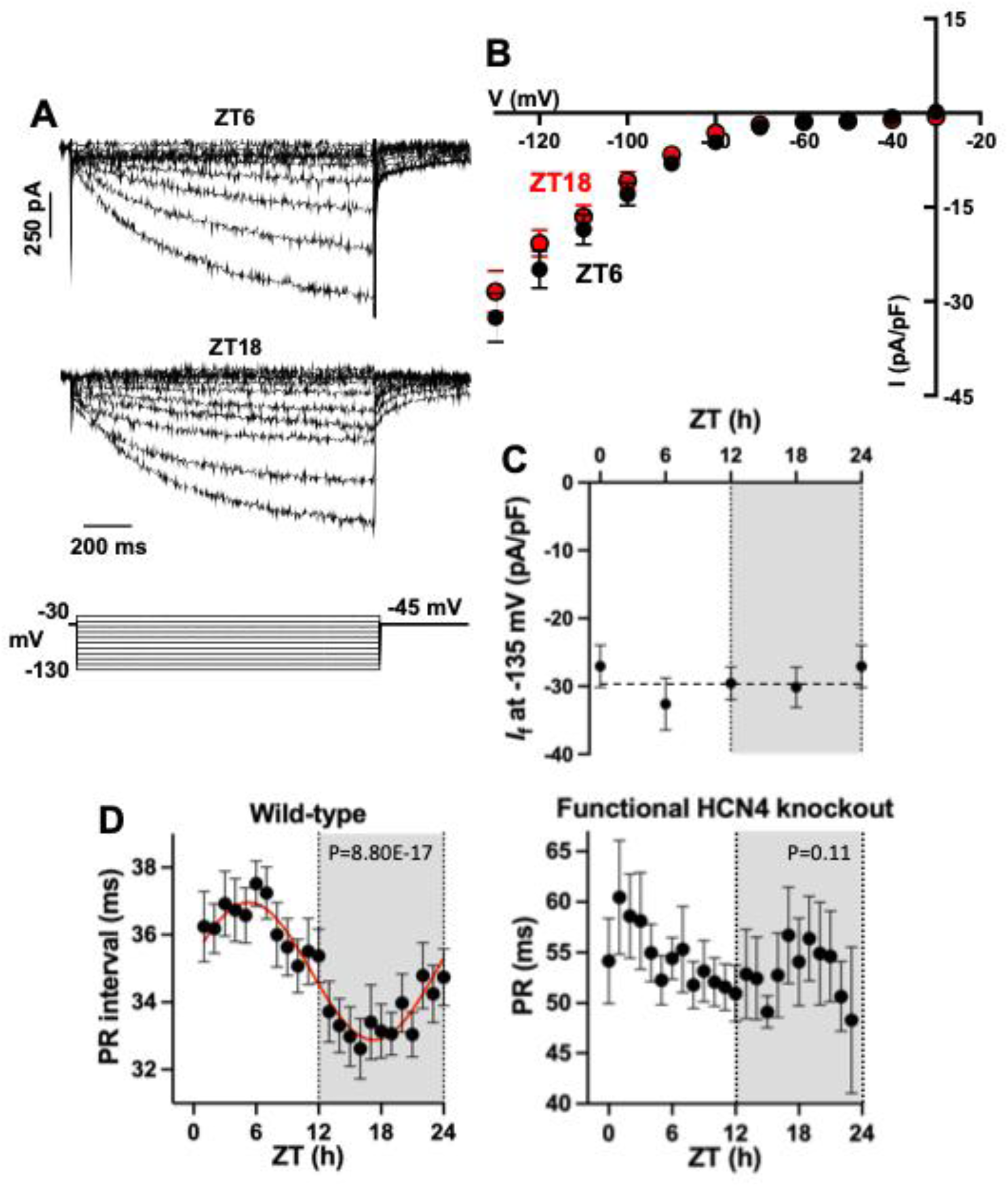
The HCN4/*I*_f_ paradox in the AV node. **A**, families of recordings of *I*_f_ at ZT6 and 18 (voltage clamp protocol shown below). **B**, current density-voltage relationships for *I*_Ca,L_ at ZT6 (n=16 or 19) and 18 (n=16-20); means±SEM shown. **C**, lack of circadian rhythm in *I*_f_ density at -135 mV. Means±SEM shown (n=19-25 myocytes at each time point). **D**, mean±SEM PR interval at 1 h intervals measured using telemetry from conscious freely moving wild-type (n=15 or 16) and functional HCN4 knockout (AYA) (n=7) mice over 24 h.

### Circadian rhythm in the contractile apparatus, metabolism and extracellular matrix

Many types of protein showed significant circadian rhythms and Fig. S10 shows the processes affected as identified by gene ontology analysis and amongst these is muscle contraction (‘Cytoskeleton in muscle cells’). The contractile proteins shown in Fig. S11 showed a significant circadian rhythm, peaked at the start of the active period, and are linked (or potentially linked) to myofilament Ca^2+^ sensitivity. These proteins could be responsible for an increase in myofilament Ca^2+^ sensitivity and muscle contractility during the active period (Fig. S12). See the Supplemental Results for details. Two proteins related to the contractile apparatus, myosin binding protein h like (*Mybphl*) and obscurin (*Obscn*), showed a significant circadian rhythm and are linked by GWAS to the PR interval (Table 1).

**Table 1.**
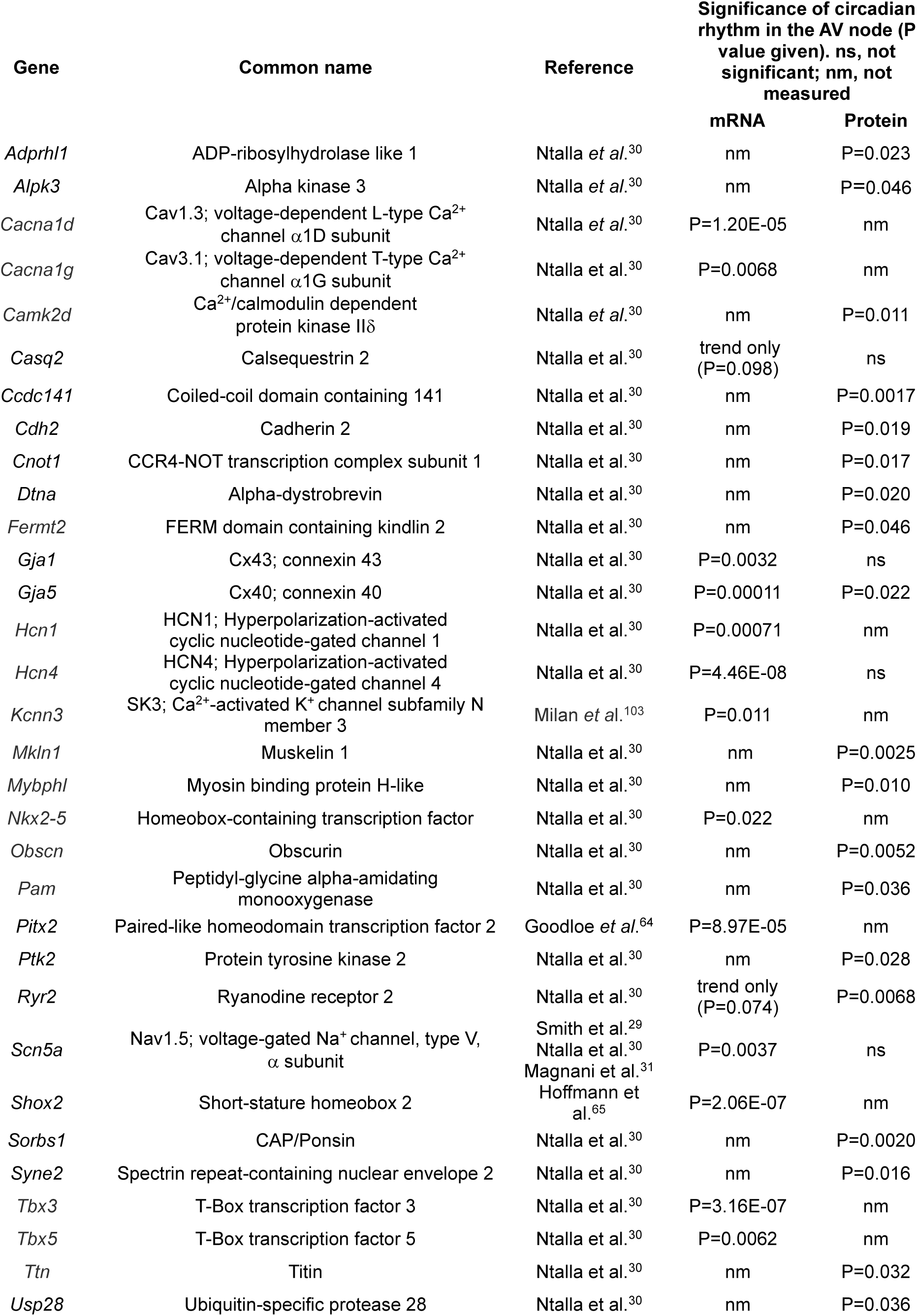
Genes identified (principally by GWAS) as influencing the PR interval and also showing a circadian rhythm.

| Gene | Common name | Reference | Significance of circadian rhythm in the AV node (P value given). ns, not significant; nm, not measured |  |
|---|---|---|---|---|
|  |  |  | mRNA | Protein |
| *Adprhl1* | ADP-ribosylhydrolase like 1 | Ntalla *et al.*<sup>30</sup> | nm | P=0.023 |
| *Alpk3* | Alpha kinase 3 | Ntalla *et al.*<sup>30</sup> | nm | P=0.046 |
| *Cacna1d* | Cav1.3; voltage-dependent L-type Ca<sup>2+</sup> channel α1D subunit | Ntalla *et al.*<sup>30</sup> | P=1.20E-05 | nm |
| *Cacna1g* | Cav3.1; voltage-dependent T-type Ca<sup>2+</sup> channel α1G subunit | Ntalla et al.<sup>30</sup> | P=0.0068 | nm |
| *Camk2d* | Ca<sup>2+</sup>/calmodulin dependent protein kinase IIδ | Ntalla *et al.*<sup>30</sup> | nm | P=0.011 |
| *Casq2* | Calsequestrin 2 | Ntalla et al.<sup>30</sup> | trend only (P=0.098) | ns |
| *Ccdc141* | Coiled-coil domain containing 141 | Ntalla et al.<sup>30</sup> | nm | P=0.0017 |
| *Cdh2* | Cadherin 2 | Ntalla et al.<sup>30</sup> | nm | P=0.019 |
| *Cnot1* | CCR4-NOT transcription complex subunit 1 | Ntalla et al.<sup>30</sup> | nm | P=0.017 |
| *Dtna* | Alpha-dystrobrevin | Ntalla et al.<sup>30</sup> | nm | P=0.020 |
| *Fermt2* | FERM domain containing kindlin 2 | Ntalla et al.<sup>30</sup> | nm | P=0.046 |
| *Gja1* | Cx43; connexin 43 | Ntalla et al.<sup>30</sup> | P=0.0032 | ns |
| *Gja5* | Cx40; connexin 40 | Ntalla et al.<sup>30</sup> | P=0.00011 | P=0.022 |
| *Hcn1* | HCN1; Hyperpolarization-activated cyclic nucleotide-gated channel 1 | Ntalla et al.<sup>30</sup> | P=0.00071 | nm |
| *Hcn4* | HCN4; Hyperpolarization-activated cyclic nucleotide-gated channel 4 | Ntalla et al.<sup>30</sup> | P=4.46E-08 | ns |
| *Kcnn3* | SK3; Ca<sup>2+</sup>-activated K<sup>+</sup> channel subfamily N member 3 | Milan *et al*.<sup>103</sup> | P=0.011 | nm |
| *Mkln1* | Muskelin 1 | Ntalla et al.<sup>30</sup> | nm | P=0.0025 |
| *Mybphl* | Myosin binding protein H-like | Ntalla et al.<sup>30</sup> | nm | P=0.010 |
| *Nkx2-5* | Homeobox-containing transcription factor | Ntalla et al.<sup>30</sup> | P=0.022 | nm |
| *Obscn* | Obscurin | Ntalla et al.<sup>30</sup> | nm | P=0.0052 |
| *Pam* | Peptidyl-glycine alpha-amidating monooxygenase | Ntalla et al.<sup>30</sup> | nm | P=0.036 |
| *Pitx2* | Paired-like homeodomain transcription factor 2 | Goodloe *et al*.<sup>64</sup> | P=8.97E-05 | nm |
| *Ptk2* | Protein tyrosine kinase 2 | Ntalla et al.<sup>30</sup> | nm | P=0.028 |
| *Ryr2* | Ryanodine receptor 2 | Ntalla et al.<sup>30</sup> | trend only (P=0.074) | P=0.0068 |
| *Scn5a* | Nav1.5; voltage-gated Na<sup>+</sup> channel, type V, α subunit | Smith et al.<sup>29</sup><br>Ntalla et al.<sup>30</sup><br>Magnani et al.<sup>31</sup> | P=0.0037 | ns |
| *Shox2* | Short-stature homeobox 2 | Hoffmann et al.<sup>65</sup> | P=2.06E-07 | nm |
| *Sorbs1* | CAP/Ponsin | Ntalla et al.<sup>30</sup> | nm | P=0.0020 |
| *Syne2* | Spectrin repeat-containing nuclear envelope 2 | Ntalla et al.<sup>30</sup> | nm | P=0.016 |
| *Tbx3* | T-Box transcription factor 3 | Ntalla et al.<sup>30</sup> | P=3.16E-07 | nm |
| *Tbx5* | T-Box transcription factor 5 | Ntalla et al.<sup>30</sup> | P=0.0062 | nm |
| *Ttn* | Titin | Ntalla et al.<sup>30</sup> | nm | P=0.032 |
| *Usp28* | Ubiquitin-specific protease 28 | Ntalla et al.<sup>30</sup> | nm | P=0.036 |

The gene ontology analysis (Fig. S10) showed that many of the proteins with significant circadian rhythms concern metabolism. Cardiac energy demand will be greater during the active period because the heart rate and potentially contractility are greater because the demand for cardiac output is greater. Glycolysis shown in Fig. 8 produces ∼5% of the ATP required by the adult human heart, while the remaining ∼95% is produced by mitochondrial oxidative phosphorylation by the electron transport chain and chemiosmosis – not shown in Fig. 8).^51^ The electron transport chain is powered by electrons, carried by NADH and FADH2, which are produced during glycolysis and the tricarboxylic acid cycle (Fig. 8). There is a hierarchy of fuels for oxidative phosphorylation: under normal conditions, fatty acids (processed by fatty acid beta-oxidation; Fig. 8) account for 60-80% of mitochondrial ATP production, and glucose (processed by glycolysis; Fig. 8) also makes a substantial contribution.^51^ Other important fuels include amino acids, including glutamine (during periods of stress such as ischaemia or anoxia; Fig. 8),^52^ lactate (during exercise; Fig.S13A)^53^ and ketone bodies (especially during fasting; Fig. S13B).^54^ Figs. 8 and S13 show that in all pathways contributing to ATP production at least one critical enzyme that potentially could act as a rate limiting step showed a significant (or near significant) circadian rhythm. In all of these cases, protein expression reached a peak at or near the start of the active period (Figs. 8 and S13). This suggests that all of the ATP producing pathways have the potential to be more active during the active period.

**Fig. 8.**
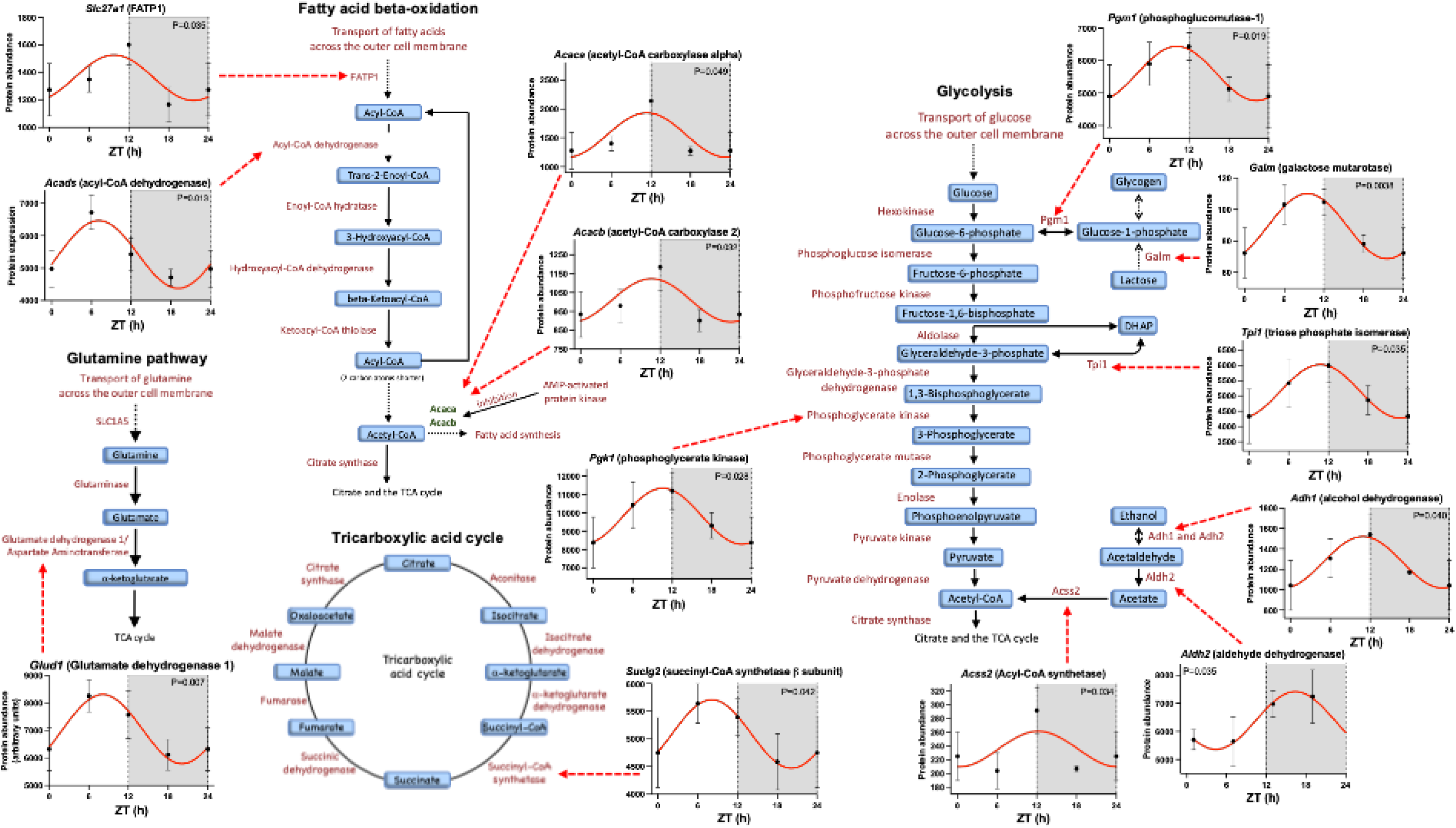
Circadian rhythm in metabolism in the AV node. Schematic diagrams of the glutamine pathway, fatty acid beta-oxidation, glycolysis and the tricarboxylic acid (TCA) cycle shown. The glutamine pathway, fatty acid beta-oxidation and glycolysis (together with lactate and ketolysis shown in Fig. S13) feed into the TCA cycle. The circadian rhythms of enzyme proteins showing significant rhythms are shown. See Fig. 2 legend for a general description of the figure and Fig. 5 legend for details of n numbers.

Gene ontology analysis showed that, amongst the proteins with significant circadian rhythms, are proteins making up the extracellular matrix (Fig. S10: ‘Cytoskeleton in muscle cells’). The major fibrillar collagens, which form cable-like fibrils and provide mechanical strength and a three-dimensional framework for tissues, are types I, II, and III. Type II collagen was not detected. *Col1a1* (collagen type I, alpha-1) was detected and is responsible for one of the two protein chains in type I collagen (Fig. S14A). It was more abundant than *Col1a2* (collagen type I, alpha-2; second of the two protein chains in type I collagen), which was also detected (Fig. S14A). Finally, *Col3a1* (collagen type III, alpha-1 responsible for type III collagen) was detected (Fig. S14A). Of these, only *Col3a1* showed a significant circadian rhythm and it reached a nadir during the active period (Fig. S14B). Type V collagen plays a role in the formation and regulation of type I collagen fibrils and *Col5a1* (collagen type V, alpha-1) showed a significant circadian rhythm and it peaked at the start of the active period (Fig. S14B). Type IV collagen is a non-fibrillar collagen that forms the network of basement membranes surrounding myocytes and is essential for structural integrity, cell adhesion and function, and collagen VI forms a microfibrillar network that acts as a bridge between the basement membrane and the surrounding interstitial matrix (containing fibrillar collagen) (Fig. S14B, inset). Type IV collagen has been repeatedly linked to cardiovascular disease.^55^ *Col4a1* (collagen type IV, alpha-1) and *Col4a2* (collagen type IV, alpha-2) are responsible for type IV collagen, and *Col4a2* showed a significant circadian rhythm peaking at the start of the active period (Fig. S14B) and *Col4a1* showed a similar pattern which bordered on significance (P=0.075; data not shown). *Col6a1* (collagen type VI, alpha-1), *Col6a2* (collagen type VI, alpha-2) and *Col6a6* (collagen type VI, alpha-6) are responsible for type VI collagen – all were highly abundant and showed a significant circadian rhythm peaking at the start of the active period (Fig. S14B). Type VI collagen plays a special role in the AV node and the rhythms may be of particular significance – see Discussion.

### Circadian rhythm in intracellular signalling

As well as a circadian rhythm in end effectors such as ion channels, metabolism, contractile proteins and the extracellular matrix, there was a circadian rhythm in intracellular signalling. There are 517 protein kinases documented for the human (kinhub.org/kinases.html) and of these 145 kinases at the protein level were detected, of which 35 (6.8%) showed a significant circadian rhythm (Table S3). The result of gene ontology analysis of the protein kinases with significant circadian rhythms is shown in Fig. S15. One relevant finding is the involvement of rhythmic protein kinases with ‘Circadian rhythm’ and ‘Circadian entrainment’ (Fig. S15). This group of kinases include *Csnk1d* (casein kinase 1 delta) and *Csnk1a1* (casein kinase 1 alpha 1). Casein kinase 1 family members fine-tune the circadian clock cycle by phosphorylating PERIOD proteins within the circadian clock mechanism (Fig. 2A).^56^ *Csnk1d* (casein kinase 1 delta) is a key regulator of the clock, phosphorylating both stabilising and destabilising sites on the *Per2* protein in a mechanism known as the phosphoswitch.^57^ Further examples will be considered. At the protein level, the alpha and delta subunits of Ca^2+^/calmodulin-dependent protein kinase II (CAMKIIα and CAMKIIδ, products of *Camk2a* and *Camk2d*) both showed significant circadian rhythms and peaked near the start of the awake period; CAMKIIδ was ∼200x more abundant than CAMKIIα (Fig. S16). The rhythms may be important, because Khoo *et al*.^58^ described CAMKII activity as being required for normal AV node conduction - genetic CaMKII inhibition by cardiac-specific expression of an inhibitory peptide prolongs the PR interval, Wenckebach cycle length and AV node effective refractory period. The effect of CaMKII has been attributed to regulation of intracellular Ca^2+^ homeostasis and the Ca^2+^ clock.^59^ Fig. S16 shows that Ca^2+^/calmodulin-dependent protein kinase I isoforms also showed a circadian rhythm. AMP-activated protein kinase (AMPK) is a key cellular energy sensor and a master regulator of metabolism. The two most abundant AMPK subunit proteins expressed (AMP-activated protein kinase subunits catalytic alpha-2 and gamma-1, products of *Prkaa2* and *Prkag1*) both showed significant circadian rhythms with expression peaking at the start of the active period (Fig. S17). AMPK has multiple actions some of which are shown in Fig. S17.^60^ For example, during times of low energy it plays a key role in maintaining cellular energy balance by stimulating energy production and also conserving energy - in part by inhibiting mTOR and thus protein synthesis, *Trim24* (TIF1A) and thus rRNA synthesis, and *Acaca*/*Acacb* (acetyl-CoA carboxylases alpha and 2) involved in fatty acid synthesis and oxidation (Fig. S17). Some, but not all, of the proteins in the AMPK pathway also showed a circadian rhythm – those showing a significant circadian rhythm are shown in Fig. S17. Fig. S17 includes other kinases which are also key energy sensors and regulators of metabolism – *Mtor* (mammalian target of rapamycin) and *Pdk1* and *Pdk4* (pyruvate dehydrogenase kinase isozymes 1 and 4). Interestingly, the AMPK pathway also promotes differentiation of human induced pluripotent stem cells into sinus node-like cells.^61^

1,482 transcription factors are listed for the mouse (https://esbl.nhlbi.nih.gov/Databases/KSBP2/Targets/Lists/TranscriptionFactors/). 51 transcription factors were detected at the protein level (limited resolution of mass spectrometry may have limited number detected). Five of the transcription factors detected (∼10%) showed significant circadian rhythms (Supplementary Table S4). A further 21 selected transcription factors were measured at the transcript level using qPCR, of which 10 (∼48%) showed significant circadian rhythms peaking at the start of or during the active period (Figs. 9 and S18; Supplementary Table S4). The 10 transcription factor transcripts include those for *Tbx3* and related transcription factors, which act together as a network (Fig. 9). Transcription factors in this pathway (*Mef2c*, *Nkx2-5*, *Shox2*,^62^ *Tbx3* and *Tbx5*^63^) are crucial for AV node development. GWAS links transcription factors in this pathway (*Nkx2-5*, Pitx2,^64^ Shox2,^65^ *Tbx3* and *Tbx5*) to the PR interval in the human (Table 1), and knockout or downregulation of transcription factors in this pathway in the mouse (*Nkx2-5*,^66^ *Pitx2*,^67^ *Tbx*3^68^ and Tbx5^69^) causes AV block. In addition, all the transcription factors in this pathway affect transcription of cardiac ion channels (Fig. 9). For example, of particular relevance here, Furtado *et al*.^70^ showed that knockout of *Nkx2-5* downregulates *Cacna1c* (Cav1.2), *Kcnh2* (ERG) and *Scn5a* (Nav1.5); in this study, *Nkx2-5*, *Cacna1c*, *Kcnh2* and *Scn5a* all showed circadian rhythms which peaked during the active period (Figs. 3, 4 and 9). Mohan et al.^71^ have argued that AV conduction is affected by subtle changes in *Tbx3* expression and showed that in a heterozygous *Tbx3* knockout mouse 409 genes are downregulated. Of particular relevance here, knockout of *Tbx3* downregulates *Kcnh2*;^71^ again both *Tbx3* and *Kcnh2* showed circadian rhythms and peaked during the active period in this study (Figs. 4 and 9). In addition, Mohan et al.^71^ showed that knockout of *Tbx3* downregulates *Hcn1* (HCN1), *Hcn4* (HCN4), *Cacna1g* (Cav3.1) and *Cacna2d2* (Cavα2δ2), all of which showed circadian rhythms (in case of *Cacna2d2*, trend only) and the transcripts peaked during the active period (Figs. 3), the same as *Tbx3* (Fig. 9). It is possible that *Nkx2-5* and *Tbx3* (and other transcription factors in the network) are responsible for the rhythms in ion channels (Figs. 3-5) and *I*_Ca,L_ and *I*_K,r_ (Fig. 6).

**Fig. 9.**
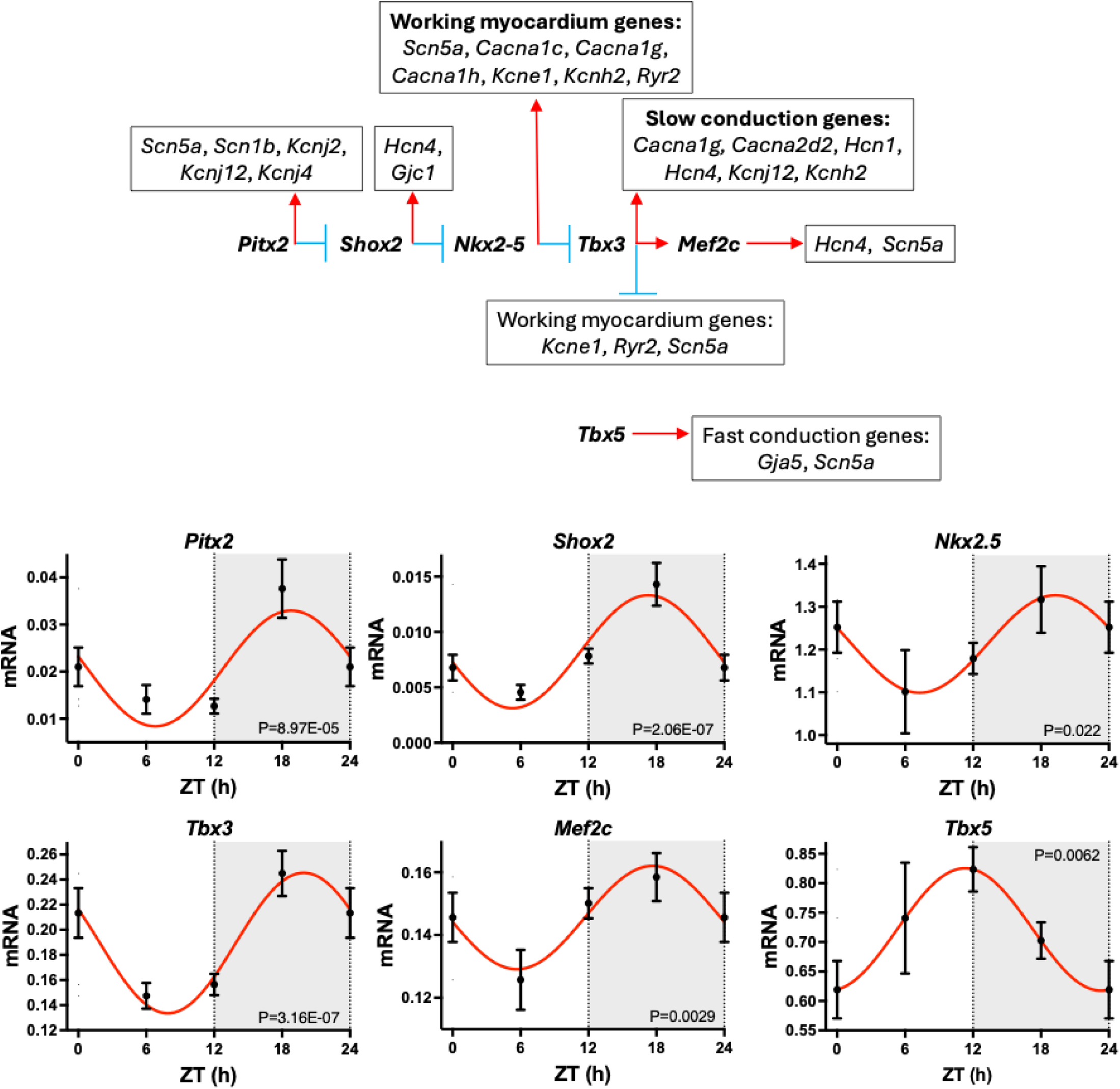
Circadian rhythm in the *Tbx3* pathway. See the legend to Fig. 2 for a general description of the figure. Data from 8-11 mice. Inset, schematic diagram of the pathway. Red arrows indicate activation of the downstream protein or process and blue lines indicate inhibition. Inset based on Espinoza-Lewis *et al*.,^99^ Mohan *et al.*,^71^ Xie *et al*.,^100^ Zhou *et al*.,^101^ Arnolds *et al*.,^102^ and Furtado *et al*.^70^

Circadian clock "slave" transcription factors are circadian clock-controlled genes (which do not in turn control the circadian clock machinery) which affect downstream output effectors. Four circadian clock slave transcription factors were observed to show significant circadian rhythms: *Klf4*, *Klf15*, *Lhx1* and *Tef* (Fig. S18). *Klf15* acts as a key transcriptional regulator of metabolism and Prosdocimo *et al*.^72^ showed *Klf15* to upregulate *Slc27a1* (FATP1) and *Pdk4* (pyruvate dehydrogenase kinase 4), both of which showed significant circadian rhythms in this study (Figs. 8 and S17), peaking approximately at the same time as *Klf15* (Fig. S18). *Klf15* is also involved in the circadian regulation of the cardiac ion channel subunit *Kcne2* (KChIP2) in the ventricles.^73^ In contrast to *Klf15*, *Klf4* is an important transcriptional regulator of mitochondrial homeostasis.

Micro-RNAs are non-coding RNAs of 19–25 nucleotides length which negatively regulate gene expression at the post-transcriptional level. They regulate expression of a wide range of cardiac ion channels.^74^ In addition, they regulate the circadian clock,^e.g.75^ contractile proteins,^76^ metabolism^77^ and fibrosis.^74^ Expression of 754 rodent micro-RNAs in AV node biopsies was determined using a TaqMan Gene Expression Assay. Expression was measured in mice at ZT0, 6, 12 and 18. 258 (34%) of the micro-RNAs showed a significant circadian rhythm. Examples are shown in Fig. S19. For the micro-RNAs with significant rhythms, QIAGEN Ingenuity Pathway Analysis (IPA) was used to identify potential targets amongst the transcripts showing significant rhythms and the results are shown in Supplementary Files S4 and S5. It is widely assumed that when a micro-RNA rises, its target transcript must fall and *vice versa* (however, this is not necessarily the case^78^) and this inverse pattern is useful for target discovery. A few selected examples of micro-RNAs and target transcripts (for ion channels or clock genes either experimentally verified or predicted) oscillating in antiphase are shown in Fig. S20. Potential target transcripts oscillating in antiphase to the micro-RNA are also highlighted in Supplementary Files S4 and S5. IPA was also used to identify potential targets of rhythmic micro-RNAs amongst the rhythmic proteins: 28 rhythmic micro-RNAs were found to potentially target 360 rhythmic proteins and Fig. S22 shows gene ontology of the potentially targeted rhythmic proteins. Of particular interest to this study, miR-1a-30 was found to be oscillating in antiphase to an experimentally verified target, *Cacna1c* (Cav1.2) responsible for *I*_Ca,L_. Other micro-RNAs were found to be in antiphase to transcripts for other experimentally verified ion channel targets (*Hcn4*, *Scn5a*, *Kcnj12* and *Kcnn3*; Fig. S20). Other potential ion channel transcript targets are listed in Supplementary Files S4 and S5. Clock transcripts are also predicted to be targeted by micro-RNAs (Fig. S20; Supplementary Files S4 and S5). Many of the potentially targeted rhythmic proteins are involved in metabolism. For example, of the proteins shown in Fig. 8, *Glud1*, *Acaca*, *Suclg2*, *Galm* and *Acss2* are potentially targeted; of the proteins in the AMP-activated protein kinase pathway involved in the regulation of metabolism shown in Fig. S17, *Prkag1*, *Tsc1*, *Trim24* and *Acaca* are potentially targeted, and *Prkaa2* and *Pdk4* are potentially targeted by multiple micro-RNAs (Supplementary File S5). See the Supplementary Results for further information concerning micro-RNAs.

## DISCUSSION

This study has investigated the circadian rhythm in the AV node. There was an analogous rhythm in the PR interval in the human and mouse. Evidence presented suggests that vagal tone is not a major factor in this. Instead, it was shown that there is an intrinsic functioning circadian clock in the AV node, and there are circadian rhythms in ion channels, ionic currents and the upstroke velocity of the action potential which is able to explain the circadian rhythm in the PR interval. In addition, the study showed rhythms in contractile proteins, metabolism and the extracellular matrix, and presumably driving all of these changes, a circadian rhythm in intracellular signalling in terms of protein kinases, transcription factors and micro-RNAs. Therefore, just as in the sinus node,^3^ the circadian rhythm in the AV node is all pervasive.

### Circadian rhythm in the PR interval

The circadian rhythms in the PR interval and heart rate are clinically important because bradyarrhythmias most commonly occur during sleep.^2^ For a long time (approaching 100 years in the case of heart rate^11,12^), both have been attributed to high vagal tone during sleep.^13,79^ In the case of the circadian rhythm in the heart rate, vagal tone plays only a minor or even no role.^5,15,16^ Now, this study has shown that knockout of the vagal tone effector gene in the AV node, *Kcnj5* (Kir3.4), only reduced the circadian rhythm in the PR interval by ∼30% (Fig. 1), suggesting that vagal tone may play only a minor role in the AV node. In addition, there is a possible alternative explanation of the decrease in the circadian rhythm in the PR interval on knockout of *Kcnj5* (Supplementary Discussion). Furthermore, this study showed that there is a functioning circadian clock in the AV node (Fig. 2) and knockout of this intrinsic clock reduced the circadian rhythm in the PR interval by ∼43-52% (Figs. 2D and S4). Ion channels are responsible for conduction through the AV node and therefore the PR interval, and there was a circadian rhythm in the expression of ion channels and the corresponding ionic currents in the AV node, in particular *Cacna1c* (Cav1.2) and *Cacna1d* (Cav1.3) and the L-type Ca^2+^ current, *I*_Ca,L_ (Figs. 3 and 6A), which could contribute to the circadian rhythm in the PR interval: the AV node has a Ca^2+^-dependent AP and the increase in *I*_Ca,L_ during the active period can explain the observed increase in the upstroke velocity of the AP at this time (Fig. 1D,E), which in turn will increase the conduction velocity and shorten the PR interval (Fig. 6D,G). A potential role for *I*_K,r_, *I*_K,ACh_, background current and even *I*_Na_ is considered in the Supplementary Discussion. The ion channel HCN4 could be playing a role, because the *Hcn4* transcript showed a circadian rhythm (Fig. 3) and knockout of *Hcn4*, identified by GWAS as a determinant of the PR interval (Table 1), slowed AV node conduction (prolonged the PR interval) and abolished the circadian rhythm in the PR interval (Figs. 7D and S9). However, how it is involved is unclear because HCN4 protein and the corresponding ionic current, *I*_f_, do not show circadian rhythms (Figs. 5 and 7); HCN4, as well as its function as an ion channel, may have a signalling role – block of HCN4 results in widespread changes in ion channel transcripts.^80^ By analogy with the sinus node,^5^ the circadian rhythm in the expression of ion channels (Figs. 3-5) could be driven by the intrinsic circadian clock in the AV node and also by factors such as plasma catecholamine and glucocorticoid – we have reported that knockout of the glucocorticoid receptor abolishes the circadian rhythm in the PR interval (as well as the circadian rhythm in heart rate).^6^ It is interesting that expression of the glucocorticoid receptor protein reached a peak at the start of the active period in this study (Fig. 23). It should be mentioned that there is another factor that could be contributing to the circadian rhythm in the PR interval *in vivo* – the core body temperature. The core body temperature shows a circadian rhythm^5,81,82^ and the lowering of the core body temperature during the sleep period is expected to slow AV node conduction and prolong the PR interval.^83,84^ Furthermore, Usai *et al*.^82^ showed that circadian rhythms in the PR interval and core body temperature occur in parallel in mice.

### GWAS

Genome-wide association studies (GWASs) have identified genes associated with the PR interval - the meta-analysis of Ntalla et al.^30^ identified 202 loci associated with the PR interval and of the genes associated with the loci, 29 of them showed a significant circadian rhythm in the AV node (Table 1). Whereas the connection of the genes with the PR interval is obscure in many cases, the connection is better understood in the cases of the ion channel genes, *Cacna1d*, *Cacna1g*, *Gja1*, *Gja5*, *Hcn1*, *Hcn4*, *Kcnn3*, *Ryr2* and *Scn5a*, the protein kinase gene, *Camk2d*, and the transcription factor genes, *Nkx2-5*, *Tbx3* and *Tbx5*; independently, this study has shown a potential role for all of these genes in the circadian rhythm in the PR interval.

### Role of intracellular signalling molecules in the circadian rhythm in the PR interval

The importance of the circadian rhythm in the AV node is shown by the plethora of circadian rhythms in intracellular signalling molecules – this study has shown significant rhythms in 35 protein kinases (Table S3), 15 transcription factors (Table S4) and 258 micro-RNAs and there are likely more that have not been detected. This study has highlighted two factors that may be responsible for the circadian rhythms in ion channels underlying the circadian rhythm in the PR interval: the Tbx3 pathway (Fig. 9) and micro-RNAs (e.g. Fig. S21). Ultimately of course, driving the circadian rhythm in intracellular signalling are extrinsic neurohumoral factors and the intrinsic circadian clock in the AV node.

### Circadian rhythm in contractile proteins, metabolism and extracellular matrix

There is limited evidence of a circadian rhythm in cardiac contractility and an increase in cardiac contractility during the active period^85,86^ – this is expected because during the active period the demand for cardiac output is increased. There is almost no data on whether cardiac contractile proteins show a circadian rhythm (although there is a report of a circadian rhythm in α-myosin heavy chain,^85^ which was not seen in the present study; P=0.57). In this study, a few contractile proteins showed significant circadian rhythms (Fig. S11), and five of them may regulate the myofilament Ca^2+^ sensitivity and this may explain why myofilament Ca^2+^ sensitivity is greater during the active period.^87,88^ Such an increase in myofilament Ca^2+^ sensitivity is expected to result in an increase in cardiac contractility and this was demonstrated at least in ventricular myocytes (Fig. S12). The heart is capable of using a wide range of substrates for energy production including fatty acids, glucose including glycogen, amino acids including glutamine^89^ (Fig. 8), lactate and ketone bodies (Fig. S13).^1^

Cardiac metabolism is well known to show a circadian rhythm.^1,90^ 70% of the heart’s energy comes from fatty acids beginning with fatty acid oxidation.^90^ In *ex vivo* (denervated) perfused hearts, fatty acid oxidation shows a circadian rhythm with a peak at ZT6 during the sleep period.^90^ About 20% of the heart’s energy comes from glucose beginning with glycolysis (glucose oxidation).^90^ Glycolysis shows a circadian rhythm and peaks at ZT18 during the active period,^90^ experiments on *ex vivo* perfused hearts suggest that glycogenolysis is higher at ZT18 during the active period,^90^ and most amino acid catabolic enzymes reach peak expression during the active period.^90^ This study shows that metabolism in the AV node also shows a circadian rhythm. Furthermore, to the best of our knowledge, this is first proteomics study of the circadian rhythm in cardiac metabolism showing the expression level of many different proteins involved in metabolism. In the present study, at least one key enzyme in the fatty acid beta-oxidation and glycolysis pathways, and the glycogen, amino acid, lactate, and ketone body utilisation pathways was upregulated at or near the start of the active period at ZT12 (Figs. 8 and S13) presumably to respond to the greater demand for cardiac output during the active period. There were also changes in the expression of metabolic regulatory proteins. The master regulator of metabolism, AMPK, was upregulated at the start of the active period as were other proteins associated with it (Fig. S17). AMPK is activated by phosphorylation and acts on downstream effectors by phosphorylation. Phospho-AMPK is known to peak at ZT18 during the active period.^90^ AMPK stimulates ATP-generating pathways including fatty acid beta-oxidation (by inhibiting ACACB; Fig. S17) and glycolysis by increasing glucose uptake and activating phosphofructokinase (not shown), while inhibiting energy-consuming processes such as protein synthesis (by suppressing mTOR, a driver of cell growth; Fig. S17) and lipid synthesis (by inhibiting ACACA; Fig. S17). In various tissues the extracellular matrix is known to show a circadian rhythm.^91^ This is also true in the heart - ∼20% of collagen undergoes daily turnover.^92^ In a review, Rana *et al*.^92^ reported circadian rhythms in collagens COL1A1, COL3A1, COL4A1, and COL4A2 in the heart. Wang *et al*.^3^ reported significant circadian rhythms in *Col1a1*, *Col3a1*, *Col6a1*, *Col6a2* and *Col15a1* transcripts in the mouse sinus node. Types I and III collagens are widely regarded as the principal collagen isoforms in the heart^93^ and, as an example, the data from Wang *et al*.^3^ in Fig. S14A shows this is the case in the mouse sinus node. However, the data from the present study of the mouse AV node shows that it is type VI collagen that is the most abundant and types I and III are poorly expressed (Fig. S14A) and the alpha-1, 2 and 6 isoforms of type VI collagen show prominent and significant circadian rhythms (Fig. S14B). This is likely to be important: embryologically, the AV node is derived from the AV canal^94^ and type VI collagen plays an important role in the embryological development of the AV canal.^95^ Type VI collagen as a key part of the extracellular matrix not only plays a role in maintaining the structural integrity of the AV node, it is also involved in regulating Ca^2+^ signalling within cardiomyocytes^96^ and patients with *Col6a1* mutations have been observed to develop bundle branch block.^97^

### Conclusions

The well-known circadian rhythm in the PR interval and the less well-known circadian rhythm in the incidence of AV block are simplistically attributed to high vagal tone during sleep, but this study shows that it is much more complex than this. There are circadian rhythms in electrophysiology, contractility, metabolism and the extracellular matrix. Probably driving all of this is a local intrinsic circadian clock in the AV node and neurohumoral factors such as glucocorticoid, and, downstream of these, a plethora of oscillating intracellular signalling pathways. Ultimately, the circadian rhythms in the PR interval and the incidence of AV block likely involve circadian rhythms in ion channel expression (and consequently ionic currents) and a circadian rhythm in the core body temperature, but it is possible that a circadian rhythm in vagal tone may still be involved.

## SUPPLEMENTARY MATERIAL

### Supplementary methods, discussion and figures

**Supplementary File S1.** Excel spreadsheet showing all transcripts detected, the P value for a significant circadian rhythm and the acrophase.

**Supplementary File S2.** Excel spreadsheet showing all proteins detected, the P value for a significant circadian rhythm and the acrophase.

**Supplementary File S3.** Excel spreadsheet showing all micro-RNAs measured, the P value for a significant circadian rhythm, and the acrophase.

**Supplementary File S4.** Potential targets of rhythmic micro-RNAs (micro-RNA focussed).

**Supplementary File S5.** Potential targets of rhythmic micro-RNAs (mRNA focussed).

## ACKNOWLEDGEMENTS

British Heart Foundation (PG/17/29/32945) to MRB, HD, AD and HP; British Heart Foundation (Control of ion channel expression in the cardiac conduction system in disease - search for potential new therapeutic targets) to MRB, HD and AD; Fondation Leducq (19CV03) to MEM, AD and MRB; Fondation pour la Recherche Médicale (EQU202403018013) to MEM; BBSRC (BB/W010801/1, BB/W018314/1, BB/Y00695X/1) to VP-V; Rosetrees Trust (CF-2021-2/133) to VP-V; EPSRC (EP/Y024168/1) to VP-V; MD UK (21GRO-PG12-0532) to VP-V; The Royal Society (RGS\R1\211432) to VP-V. Novo Nordisk Fonden (Grant NNF18OC0032728) to MBT.

## Notes

### Competing Interest Statement

The authors have declared no competing interest.

