## Supplemental figures and Tables for "Circadian rhythm in the atrioventricular node - in electrophysiology, contractility, metabolism, intracellular signalling and extracellular matrix"

### Authorship

This is a multi-disciplinary, multi-centre study. All authors played significant roles; two share first authorship and six share senior authorship.

### SUPPLEMENTARY METHODS

#### Electrocardiogram (ECG) recordings from human subjects at the Erasmus Medical Center

ECG recordings from 10 human subjects were analysed. The subjects were healthy patients who had cardiovascular screening due to sudden cardiac death in the family, and in whom no signs of cardiomyopathies or channelopathies were found. Echocardiographic data showed normal sized atria and ventricles. The sex and age of the patients is below

| Patient number | Sex | Age (years) |
|---|---|---|
| 1 | female | 59 |
| 2 | female | 20 |
| 3 | female | 33 |
| 4 | male | 25 |
| 5 | male | 24 |
| 6 | female | 29 |
| 7 | female | 34 |
| 8 | female | 52 |
| 9 | female | 44 |
| 10 | male | 32 |

Continuous three-channel bipolar Holter ECG recordings acquired at a sampling frequency of 128 Hz were analysed. Night time was from 00:00–06:00 and day time was from 08:00–22:00 and 06:00–08:00 and 22:00–00:00 were designated as transition periods. P-wave onset was detected automatically from three bipolar ECG leads after signal-quality assessment. Analyses were restricted to intervals in which both the current and preceding QRS complexes were classified as normal. Across the 10 subjects, a total of 1,017,739 P-waves were accepted, ranging from 34,004 to 209,185 per patient. The signal quality varied substantially between recordings, but there was a large number of analysable beats for all subjects. The PR interval was calculated as the time difference between consecutive annotated P-waves and QRS complexes.The RR interval was calculated as the time difference between consecutive annotated normal QRS complexes. Over 24 h, PR and RR intervals were averaged during 30 min time bins. To account for differences in intervals among subjects, for each subject, the intervals were calculated with reference to the median over the 24 h.

#### Telemetry recording of heart rate of the mouse at the University of Montpellier

All animal procedures were in accordance with the Guide for the Care and Use of Laboratory Animals published by the US National Institute of Health (NIH Publication No. 85-23, revised 1996) and European directives (2010/63/EU). Experimental procedures were approved by the Ethical Panel of the University of Montpellier and the French Ministry of Agriculture (protocol no: 2017010310594939). Telemetry recording of the electrocardiogram (ECG) was carried out as previously described[1] in freely moving wild-type and Kir3.4$^{-/-}$ (*Girk4*$^{-/-}$) mice and in double transgenic mice conditionally expressing dominant negative human HCN4 (hHCN4) subunits in a heart-specific manner by an α-MHC driven transgenic promoter and in a time-controlled way by a tetracyclinesensitive (tet-off) transgenic responder. This enabled conditional expression of hemagglutinin (HA)-tagged hHCN4 subunits carrying a silenced channel in which the GYG sequence in the channel pore was replaced by an AYA sequence (AYA mouse),[2] or cAMP-insensitive hHCN4 subunits lacking part of the c-linker and the cAMP binding domain either in the atria (SND2A), or in the atria and ventricles (SND2F).[3,4] Conditional expression of the dominant negative hHCN4 subunits was obtained by removing doxycycline from the food pellet for at least six weeks before ECG recording, as described previously.[2] All mouse lines were established on a C57Bl/6J genetic background. After local anaesthesia, a midline incision was made on the back along the spine to insert a telemetry transmitter (ETA-F10, DSI, St-Paul, NM, USA) into a subcutaneous pocket with paired wire electrodes placed over the thorax in DII derivation against the heart axis. Local anaesthesia was obtained with lidocaine (1%) injected subcutaneously at the sites of electrodes and transmitter implantation. To manage possible post-surgery pain, ibuprofen was added to the drinking water for four days after implantation. Mice were housed in individual cages with *ad libitum* access to food and water and were exposed to a standard 12 h:12 h light-dark cycle in a thermostatically controlled room (21–22°C) with 40−60% humidity. Experiments were initiated after at least 15 days of recovery from surgical implantation. ECG signals were recorded using a telemetry receiver connected to an analogue-to-digital conversion acquisition system (DSI, St-Paul, NM, USA). Recordings started at 8:30 am and were stopped at 8:30 a.m. of the following day. Heart rates were determined from ventricular interbeat intervals with Ponemah Physiology Platform 6.51 (DSI, St. Paul, NM, USA) software. PR intervals were measured by analysing a 5 min period each hour for 24 h period with ECG Auto 1.5.7 (EMKA Technologies, Paris, France) software. At the end of the protocol, animals were killed by cervical dislocation, according to the protocol approved by the ethical committee

**RNA isolation from the mouse atrioventricular (AV) node and quantitative polymerase chain reaction (qPCR) at the University of Manchester**

All animal procedures were approved by the University of Manchester in accordance with the UK Animals (Scientific Procedures) Act 1986. Experiments were performed on male C57Bl/6J mice aged 8-12 weeks and 25-30 g body weight. Mice were housed under a strict 12 h:12 h light-dark cycle in a temperature-controlled environment (22°C) with access to food and water *ad libitum*. Animals were housed under these conditions for at least seven days prior to tissue collection. Mice were sacrificed by cervical dislocation at 6 h intervals across the 24 h cycle: at zeitgeber time (ZT) 0 (n=11), 6 (n=9), 12 (n=10) and 18 (n=10) (ZT0 denotes the start of the light period and ZT12 the start of the dark period). A ~1x1 mm biopsy of AV node tissue was collected from the triangle of Koch, near the coronary sinus on the interatrial septum,[5] and snap frozen in liquid $N_2$ and stored at -80°C. Total RNA was isolated from frozen biopsies using an RNeasy Micro kit (Qiagen). RNA purity and quantity were determined using a NanoDrop ND-1000 spectrophotometer (NanoDrop Technologies). Synthesis of cDNA was performed using the High-Capacity RNA-to-cDNA Kit (Applied Biosystems, catalogue number 4387406) using the manufacturer's protocol in which 150 ng of RNA in 9 $\mu$l of nuclease-free $H_2O$ was added to 10 $\mu$l of 2x RT Buffer Mix + 1 $\mu$l of 20x RT Enzyme mix. qPCR was carried out using a QuantStudio 7 Flex Real-Time PCR System (Applied Biosystems/Life Technologies Corporation). Gene expression analysis of 96 transcripts, including putative housekeeping genes and genes involved in cardiac function, was performed using custom-designed TaqMan Low-Density Array cards (Applied Biosystems, catalogue number 4342259; format 96a) as previously described[6] and according to the manufacturer's instructions. Briefly, 150 ng of cDNA and TaqMan Fast Advanced Master Mix (Applied Biosystems) were loaded into each port of microfluidic cards containing fluorogenic TaqMan probes and primers. Thermal cycling conditions were 50°C for 2 min and 92°C for 10 min, followed by 40 cycles at 95°C for 1 s and 60°C for 20 s. Data were collected and analysed with QuantStudio Real-Time PCR software. Three housekeeping genes, *Pgk1*, *Gapdh* and *Csnk1e*, were selected as optimal (most stable) endogenous controls (low M-values). Transcript expression levels were calculated using the ΔCt method. Cosinor analysis was used to determine whether there was a circadian rhythm in the expression of a transcript. As previously described,[7,8] a cosine function (with a fixed wavelength of 24 h) was fitted to expression data at the four time points (ZT0, 6, 12 and 18) by least-squares analysis:

$$Y(t) = M + A\cos\left(\frac{2\pi t}{\tau} + \phi\right) + \varepsilon$$

where *Y(t)* is the expression at time *t* (in hours), *M* is the mesor (mean value), *A* is the amplitude, $\varphi$ is the acrophase (time of peak expression), $\tau$ is the assumed period (fixed at 24 h), and $\varepsilon$ represents residual error. A rhythm was considered statistically significant if the zero-amplitude test yielded a P value of <0.05 (Bonferroni adjusted for multiple testing). This test determines whether the fitted cosine function explains the data better than a flat line, ruling out the possibility that the amplitude is zero.

**Bioluminescence experiments on the mouse AV node at the University of Manchester**

As described above, all animal procedures were approved by the University of Manchester in accordance with the UK Animals (Scientific Procedures) Act 1986. Male transgenic knock-in mice (*Per1*::LUC), in which luciferase expression is driven by the mouse *Per1* promoter and 5'-UTR elements[9] (originally obtained from Professor G.T. van der Horst, Erasmus MC, Rotterdam, Netherlands) were used at 4-6 months of age as described previously.[10] Total bioluminescence was recorded for 96 h from an AV node preparation using photomultiplier tube assemblies (H8259/R7518P, Hamamatsu) using previously described methods.[11] 3.5 cm dishes were prepared under sterile conditions with a layer of autoclaved high-vacuum grease (Dow Corning) on the top rim, a Millicell insert (Millipore) and 1 ml recording medium consisting of DMEM (Invitrogen) supplemented with 10% foetal calf serum (FCS) and 1% penicillin/streptomycin and 0.1 mM luciferin in autoclaved Milli-Q water. A freshly dissected AV node was placed on the Millicell insert following which the dish was sealed with a glass coverslip and placed into the photomultiplier tube assembly housed in a light-tight incubator (GalaxyR+; RS Biotech) and maintained at 37°C. Photon counts were integrated for 599 s every 600 s. Bioluminescence data were detrended by subtracting a 24 h running average from the raw data and smoothed with a 3 h running average. A 3-parameter sinusoidal fit to the data was performed with a fixed wavelength of 24 h due to the strict 12:12 light:dark cycle of the laboratory.

**Bioluminescence experiments on the mouse AV node at the University of Liverpool**

All animal experiments were conducted under the aegis of the 1986 Home Office Animal Procedures Act (UK). PERIOD2::luciferase (PER2::*luc*) reporter mice were generated as described previously.[12] Mice were kept on a standard maintenance chow *ad libitum* and light-entrained under a 12 h light:12 h dark regimen. 6-7 months of age adult male and female mice were used and culled at ZT8. A LumiCycle 32-channel luminometer (Actimetrics), housed inside a 37°C incubator in 0% $CO_2$, was used to track the bioluminescence from micro-dissected AV node tissue and isolated atria from PER2::*luc* reporter mice in real-time. Photons were recorded every 10 min for 1 min and presented as counts/min. Heart tissues were placed on Millicell organotypic cell culture inserts (Merck) in Corning 35 mm dishes containing 1 ml of recording medium and placed into the LumiCycle for recording. Recording medium contained Gibco phenol-free DMEM with low glucose (1000 mg/l; Thermo Fisher Scientific), supplemented with 2 mM L-glutamine, 10 mM HEPES, 8.8 mM $NaHCO_3$, 1 mM sodium pyruvate, 5% foetal bovine serum (Thermo Fisher Scientific) and 0.1 mM beetle luciferin (Promega). All reagents were purchased from Sigma-Aldrich unless otherwise stated. Bioluminescence traces were exported as either 'Raw' data or 'Baseline Subtracted' data and circadian parameter analyses were performed on baseline subtracted traces fitted with a damped sine wave in order to determine circadian period, amplitude, and phase (excluding the first 24 h of recording) using LumiCycle analysis software. After 5-6 days of real-time bioluminescence recording in constant conditions, heart tissues were taken out of the LumiCycle in the dark and synchronised with 0.1 mM dexamethasone in recording medium for 30 min within a 37°C incubator. Dishes were re-sealed with 40 mm glass coverslips and returned to the LumiCycle for further recording over 4-5 days. Amplitude of PER2::*luc* oscillations was measured as the peak-trough expression level in photon counts of the first stable cycle (i.e. second peak) in constant conditions or following synchronisation *in vitro*. The period of PER2::*luc* oscillations was calculated as the peak-to-peak time length (h) for at least three full circadian cycles starting from the first stable cycle in constant conditions or following synchronisation *in vitro*. Phase of PER2::*luc* oscillations was measured as the projected circadian time of the peak of the first stable cycle in constant conditions or following synchronisation *in vitro* relative to the ZT12 of the light:dark entrained animals, which is denoted as a peak of PER2::luc *in vivo*.

**Experiments on *Bmal1* knockout mice at the University of Manchester**

As described above, all animal procedures were approved by the University of Manchester in accordance with the UK Animals (Scientific Procedures) Act 1986. Male 10-14 week old cardiac-specific *Bmal1* knockout mice were used at the University of Manchester. They were generated by breeding *Bmal1*$^{flox/flox}$ mice with α-myosin heavy chain (α-MHC)-Cre mice. *Bmal1*$^{flox/flox}$ mice have loxP sites flanking exon 8 of the *Bmal1* gene (B6.129S4(Cg)-Arntl$^{tm1Weit}$, Jackson Laboratory, ME, USA)[13] and the cardiomyocyte-specific α-myosin-heavy chain (*Myh6*) promoter drives expression of Cre recombinase in the α-myosin heavy chain-Cre mice.[14] In the resulting offspring, the exon encoding the *Bmal1* basic helix-loop-helix (bHLH) domain was deleted in the Cre recombinase-expressing cardiac tissues. These *Bmal1* knockout mice were used in a previous study.[15] Over 24 h, the electrocardiogram (ECG) was recorded using radiotelemetry (DSI) in conscious, freely moving mice housed under 12 h light:12 h dark conditions.

**Experiments on *Bmal1* knockout mice at the University of Copenhagen**
Experiments at the University of Copenhagen were performed under license (2019-15-0201-00333) from the Ministry of Food, Agriculture and Fisheries, Denmark and were in accordance with the European Union legislation for the protection of animals used for scientific purposes (Directive 2010/63/EU). CBK cardiac-specific *Bmal1* knockout mice were used. The CBK mice were generated in the same manner as the *Bmal1* knockout at the University of Manchester, but independently.[16] The mice were generated by crossing *Bmal1*$^{flox/flox}$ mice (B6.129S4(Cg)-Arntl$^{tm1Weit}$, Jackson Laboratory, ME, USA) with α-MHC-CRE$^{+/-}$ mice (heterozygous cardiac-specific α-myosin heavy chain-driven *Cre* recombinase transgenic mice). CBK (BMAL1$^{flox/flox}$/α-MHC-CRE$^{+/-}$) and littermate controls (BMAL1$^{flox/flox}$/α-MHC-CRE$^{-/-}$) on a C57Bl/6J background were used. Over 24 h, the ECG was recorded using radiotelemetry (Data Sciences International) in conscious, freely moving mice housed at a constant 22°C under 12 h light:12 h dark conditions and provided with food and tap water *ad libitum*. An averaged ECG was calculated every hour in every mouse and the PR interval was determined in a semi-automatic manner using LabChart (ADInstruments). The data reported here were obtained through a reanalysis of data previously reported by Gottlieb *et al*.[17]

**Proteomics of the mouse AV node at the University of Manchester and Imperial College London**
As described above, all animal procedures were approved by the University of Manchester in accordance with the UK Animals (Scientific Procedures) Act 1986. Experiments were performed on male C57BL/6J mice aged 8-12 weeks. Mice were housed in a random order on shelves under a strict 12 h:12 h light-dark cycle in a temperature-controlled environment (22°C) with access to food and water *ad libitum*. Animals were housed under these conditions for at least seven days prior to tissue collection. Mice were sacrificed by cervical dislocation at 6 h intervals across the 24 h cycle: at ZT0, 6, 12 and 18. 34 mice were randomly into divided into the ZT0 6, 12 and 18 groups using a random number generator (n=10 mice at ZT6 and n=8 for the other groups). AV node tissue from 34 mice was collected as previously described[18] and snap frozen in liquid $N_2$ and stored at -80°C.

*Quantitative proteomics*. DTU Proteomics Core and Biogenity carried out the proteomics sample analysis. To increase protein yield, two AV node preparations were pooled to obtain one biological replicate resulting in 4 or 5 biological replicates per ZT group.

*Sample preparation*. 75 µl of lysis buffer (50 mM Tris-HCl, pH 8.5, 5 mM EDTA, 150 mM NaCl, 10 mM KCl, 1% Triton X-100, 5 mM $Na^+$ fluoride, 5 mM beta-glycerophosphate, and 1 mM $Na^+$ orthovanadate) containing Roche complete protease inhibitor was added to each sample along with a 3 mm tungsten carbide bead (Qiagen). Samples were treated twice in the TissueLyser II homogenizer (Qiagen) going from 3-30 Hz in 60 s. The beads were removed and samples were transferred to clean Eppendorf LoBind tubes and incubated for 2 h at -4°C. The samples were centrifuged at 15,000 g for 20 min at 4°C and supernatant precipitated with 25% acetone and left at -20°C. The samples were centrifuged at 400 g for 1.5 min and the pellets were left to dry. Samples were added to 20 µl lysis buffer (6 M guanidinium hydrochloride, 10 mM TCEP, 40 mM CAA, 50 mM HEPES, pH 8.5) and treated twice in the TissueLyser II homogenizer (Qiagen) going from 3-30 Hz in 60 s. The beads were removed and samples were transferred to clean Eppendorf LoBind tubes and boiled at 95°C for 5 min, followed by sonication on high for 5x 60 s on and 30 s off in a Bioruptor Pico sonication water bath (Diagenode) at 4°C. Samples were centrifuged at 18,000 RCF for 10 min, and supernatants were transferred to clean Eppendorf Protein LoBind tubes. Protein concentrations were determined by BCA Rapid Gold (Thermo Scientific) and 20 µg protein was taken forward for digestion. Sample volume was normalised between samples with additional lysis buffer and diluted 3x with digestion buffer (10% acetonitrile in 50 mM HEPES pH 8.5) and 400 ng of LysC (Mass

Spectrometry Grade, Wako) was added. Samples were incubated for 3 h at 37°C, shaking at 750 rpm. After LysC digestion, samples were diluted 10x in digestion buffer and digested with 200 ng trypsin (Mass Spectrometry Grade, Sigma Aldrich) for 19 h at 37°C, shaking at 750 rpm. Digestion was stopped by adding 2% trifluoroacetic acid (TFA) to a final concentration of 1%. The resulting peptides were desalted on a SOLAµ SPE plate (HRP, Thermo). Between each application, the solvent was spun through by centrifugation at 350 RCF. For each sample, the filters were activated with 200 µl of 100% methanol, and then 200 µl of 80% acetonitrile and 0.1% formic acid. The filters were equilibrated twice with 200 µl of 1% TFA and 3% acetonitrile, after which the sample was loaded. After washing the tips twice with 200 µl of 0.1% formic acid, the peptides were eluted into clean Eppendorf Protein LoBind tubes using 40% acetonitrile and 0.1% formic acid. The eluted peptides were concentrated in an Eppendorf Speedvac and reconstituted in 50 mM HEPES (pH 8.5) for TMT labeling with 16plex tags (Thermo). Labelling was done according to the manufacturer's instructions. TFA was added to acidify and bring the acetonitrile concentration down to <5%. Prior to mass spectrometry analysis, the peptides were desalted following the same procedure as described previously and fractionated using an offline ThermoFisher Ultimate3000 liquid chromatography system using high pH fractionation (5 mM ammonium bicarbonate, pH 10) at 5 µl/min flow rate. 15 µg of each sample pool was separated over a 120-min gradient (5% to 35% acetonitrile), while collecting fractions every 130 s. The resulting 60 fractions were pooled into 30 final fractions, acidified to pH<2 with 1% TFA and loaded onto EvoSep stage tips according to the manufacturer's protocol.

*Mass spectrometry (MS) analysis*. For each fraction, peptides were analysed using the pre-set '30 samples per day' method on the EvoSep One instrument. Peptides were eluted over a 44-min gradient, and analysed with an Orbitrap EclipseTM TribridTM instrument (Thermo Fisher Scientific) with FAIMS ProTM Interface (Thermo Fisher Scientific) switched between CVs of -50 V and -70 V with cycle times of 1.5 s. Full MS spectra were collected at a resolution of 120,000, with normalised AGC target set to 'standard' or maximum injection time of 50 ms and a scan range of 375–1500 m/z. MS1 precursors with an intensity of >5×103 and charge state of 2-7 were selected for MS2 analysis. Dynamic exclusion was set to 60 s, the exclusion list was shared between CV values, and Advanced Peak Determination was set to 'off'. The precursor fit threshold was set to 70% with a fit window of 0.7 m/z for MS2. Precursors selected for MS2 were isolated in the quadrupole with a 0.7 m/z window. Ions were collected for a maximum injection time of 50 ms and normalized AGC target set to 'standard'. Fragmentation was performed with a CID normalised collision energy of 35% and MS2 spectra were acquired in the IT at scan rate rapid. Precursors were subsequently filtered with an isobaric tag loss exclusion of TMT and precursor mass exclusion set to 18 m/z low and 5 m/z high. Precursors were isolated for an MS3 scan using the quadrupole with a 2 m/z window, and ions were collected for a maximum injection time of 86 ms and normalised AGC target of 200%. Turbo TMT was deactivated and number of dependent scans set to 5. Isolated precursors were fragmented again with 63% normalised HCD collision energy, and MS3 spectra were acquired in the orbitrap at 50,000 resolution with a scan range of 100-500 m/z. MS performance was verified for consistency by running complex cell lysate quality control standard.

*Data processing*. The raw files were analysed using Proteome Discoverer 2.4 (Thermo Fisher Scientific). TMT reporter ion quantitation was enabled in the processing and consensus steps, and spectra were matched against the Mus Musculus database obtained from UniProt. Dynamic modifications were set as Oxidation (M), and Acetyl on protein N-termini. Cysteine carbamidomethyl (C) and TMT 16-plex (peptide N-termini and K) were set as static modifications. All results were filtered to a 1% FDR, and protein quantitation done using the built-in Minora Feature Detector with statistical significance testing done with the built-in t test. The peptide abundances were normalised based on total peptide amount. Thereby, the total sum of identified peptides in a channel was normalised to the channel with the highest abundance. The protein or peptide abundances were normalised to total peptide abundance. To analyse circadian rhythms in protein abundance over 24 h, data were processed with DiscoRhythm[19] (v1.26.0) in R (v4.5).

**Patch clamp recordings of ionic currents from mouse AV node myocytes at the University of Montpellier**

As described above, all animal procedures were in accordance with guidelines published by the US National Institute of Health as well as European directives, and experimental procedures were approved by the University of Montpellier and French Ministry of Agriculture. AV node myocytes were isolated at ZT6 and ZT18 from 2- to 4-month-old wild-type C57Bl/6J mice as previously described.[20,21] The AV node region was excised in warmed (35°C) Tyrode solution containing (in mM): NaCl, 140; KCl, 5.4; $CaCl_2$, 1.8; $MgCl_2$, 1; Hepes, 5; and D-glucose, 5.5 (adjusted to pH 7.4 with NaOH). AV node tissue was then transferred into a "low $Ca^{2+}$" solution containing (in mM): 140 NaCl, 5.4 KCl, 0.5 $MgCl_2$, 1.2 $KH_2PO_4$, 50 taurine, 5.5 D-glucose, 1 mg/ml BSA, and 5 Hepes (adjusted to pH 6.9 with NaOH) for 4-6 min. The tissue was then transferred into low-$Ca^{2+}$ solution containing liberase TH (0.15 mg/ml, Roche, Boulogne-Billancourt, France) and elastase (0.48 mg/ml, Worthington, Lakewood, NJ, USA). Digestion was carried out for 15-20 min at 36°C. Digestion was stopped by washing tissue strips in a modified 'Kraftbrühe' (KB) medium containing (in mM): 100 L-glutamic acid $K^+$ salt, 10 L-aspartic acid $K^+$ salt, 25 KCl, 80 KOH, 10 $KH_2PO_4$, 2 $MgSO_4$, 5 creatine, 0.5 EGTA, 20 D-glucose, 20 taurine, 1 mg/ml BSA, and 5 Hepes (adjusted to pH 7.2 4 with KOH). Single AV node myocytes were isolated from tissue by manual agitation using a flame-forged Pasteur's pipette in KB solution at 36°C. To recover the automaticity of the AV node cells, $Ca^{2+}$ was gradually reintroduced in the cell storage solution to a final concentration of 1.8 mM.[22] AV node cells were then left to rest for 1 h before recording. AV node cells were harvested in custom-made chambers with glass bottoms for cell attachment and perfused with Tyrode's solution warmed to 36°C (multi-MPRE8 heating pen, Cell Micro Controls, Norfolk, VA, USA) before recording and containing (in mM): 140 NaCl, 5.4 KCl, 1 $MgCl_2$, 1.8 $CaCl_2$, 5.5 D-glucose and 5 Hepes (adjusted to pH 7.4 with NaOH). The funny current ($I_f$), the fast delayed rectifier $K^+$ current ($I_{K,r}$) and the ACh-activated $K^+$ current ($I_{K,ACh}$) were recorded using the standard whole-cell variation of the patch-clamp technique in pre-warmed (36°C) Tyrode's solution. Whole-cell electrodes had a resistance of 3–5 MΩ when filled with $K^+$ aspartate-based (K-Asp) intracellular solution containing (in mM): 80 L-aspartic acid $K^+$ salt, 50 KCl, 1 $MgCl_2$, 5 Hepes, 2 $CaCl_2$, 5 EGTA and 3 ATP-$Na^+$ salt (adjusted to pH 7.2 with KOH). For recording the L-type $Ca^{2+}$ current, $I_{Ca,L}$, $K^+$ aspartate intracellular solution was replaced with a warmed to 36C° (multi-MPRE8 heating pen, Cell Micro Controls, Norfolk, VA, USA) solution containing (in mM): 125 CsOH, 20 TEA-Cl, 1.2 $CaCl_2$, 5 Mg-ATP, 0.1 $Li_2$-GTP, 5 EGTA, and 10 HEPES (pH adjusted to 7.2 with HCl) and Tyrode's solution was replaced with a solution containing (mM): 135 tetraethylammonium chloride (TEA-Cl), 10 4-aminopyridine (4-AP), 1 $MgCl_2$, 0.03 tetrodotoxin (TTX), 5.5 D-glucose, 2 $CaCl_2$, and 10 Hepes (adjusted to pH = 7.2 with HCl). All electrophysiological signals were recorded using a Multiclamp 700B patch clamp amplifier connected to a Digidata 1550B interface (Molecular Devices, San Jose, CA, USA).

**Measurement of rat ventricular myocyte contraction at the United Arab Emirates University**
All animal procedures were approved by the United Arab Emirates University Animal Ethics Committee (reference: ERA_2024_5443). Male Wistar rats (~300 g) were housed in environmentally controlled rooms under a 12 h light-dark cycle and randomly assigned to one of two separate rooms with opposing light schedules. The normal light cycle room had lights on at 6:00 am, whereas the reverse light cycle room had lights on at 6:00 pm. Animals were euthanised at 9:30 am by guillotine decapitation. Left ventricular myocytes were enzymatically isolated using a Langendorff perfusion system, as previously described.[23] During experiments, isolated myocytes were superfused (2-5 ml/min) with normal Tyrode's solution containing (in mM): 140.0 NaCl, 5.0 KCl, 1.0 $MgCl_2$, 10.0 glucose, 5.0 HEPES and 1.8 $CaCl_2$ (pH 7.4). Unloaded left ventricular myocyte shortening was recorded using a video edge detection system (VED-114; Crystal Biotech, Northborough, MA, USA). Resting cell length and twitch shortening amplitude (expressed as a percentage of resting cell length) were measured in electrically stimulated myocytes (1 Hz) maintained at 35°C. Data acquisition and analysis were performed using WinWCP v5.2.4 software (John Dempster, Strathclyde Institute for Pharmacy & Biomedical Sciences, University of Strathclyde, Glasgow, UK).

**Measurement of micro-RNA in the mouse AV node at the University of Manchester**
As described above, all animal procedures were approved by the University of Manchester in accordance with the UK Animals (Scientific Procedures) Act 1986. Experiments were performed on male C57BL/6J mice aged 10-12 weeks. Mice were housed under a strict 12 h:12 h light-dark cycle in a temperature-controlled environment (22°C) with access to food and water *ad libitum*. Animals were housed under these conditions for at least seven days prior to tissue collection. Mice were sacrificed by cervical dislocation at 6 h intervals across the 24 h cycle: at ZT0, 6, 12 and 18. 32 mice were randomly into divided into the ZT0 6, 12 and 18 groups (n=8 in each group). AV node tissue from the mice was collected as previously described[12] and snap frozen in liquid $N_2$ and stored

at -80°C. Total RNA was isolated from AV node biopsies using a RNeasy Micro Kit (Qiagen) according to the manufacturer's instructions. Quantity and purity of RNA was measured using a NanoDrop ND-100 spectrophotometer (NanoDrop Technologies). cDNA synthesis was performed using the Superscript VILO cDNA Synthesis kit (Thermo Fisher Scientific) using the manufacturer's protocol. qPCR was carried out using an ABI Prism 7900 HT Sequence Detection System (Applied Biosystems). TaqMan Array Rodent MicroRNA A&B Cards Set v3.0 (Thermo Fisher Scientific) were used to measure expression levels of 762 micro-RNAs and five endogenous house keepers. Briefly, 150 ng of cDNA and TaqMan Universal Master Mix (Applied Biosystems) were loaded into microfluidic card ports containing TaqMan probes and primers. Thermal cycling conditions were 50°C for 2 min and 94.5°C for 10 min, followed by 40 cycles at 97°C for 30 s and 59.7°C for 1 min. Data were uploaded onto the Thermo Fisher Cloud platform and Cq values exported. RealTime Statminer (v4.1, Integromics) was used for analysis. Housekeepers snoRNA202 and U6 snRNA were selected using the Minimum Variance approach. Transcript expression levels were calculated using the ΔCt method. Median Absolute Deviation analysis was performed and outliers removed. Circadian rhythmicity of micro-RNA expression was assessed using cosinor analysis performed using the cosinor2 R package v0.2.1[8] in R Studio v4.6.0.[24]

**Bioinformatics analysis at the University of Chester**
Ingenuity Pathway Analysis[25] (IPA; QIAGEN) was used to identify potential links between micro-RNAs with circadian rhythms and either transcripts or proteins with circadian rhythms. Circadian targets were first identified using cosinor analysis. Micro-RNAs demonstrating significant rhythmicity ($P<0.01$) were selected for downstream analysis, while transcripts and proteins meeting a threshold of $P<0.05$ were used. Protein identifiers were mapped to corresponding gene symbols using standard nomenclature to ensure compatibility with IPA. Micro-RNA-target interactions were explored using the micro-RNA Target Filter tool, integrating both experimentally validated and high-confidence predicted interactions from sources including TargetScan, TarBase, and the Ingenuity Knowledge Base. Analyses were restricted to the mouse, and both direct and indirect relationships were considered. Results (micro-RNA-target interaction tables) were exported and assessed.

**Computer modelling at Morinomiya University of Medical Sciences**
A simplified one-dimensional multicellular string model for the right atrium and AV node of the rabbit developed by Inada *et al.*[26] was used to simulate electrical conduction from the right atrium to the bundle of His. The model includes linked parallel fast and slow AV node pathways. The principal pathway, the fast pathway, consists of 100 atrial, 50 atrio-nodal, 50 nodal, and 50 nodo-His myocytes. Atrial myocytes are represented by the Lindblad *et al.*[27] model and the atrio-nodal, nodal, nodo-His myocytes are represented by models from Inada *et al.*[26] All models utilise Hodgkin-Huxley type differential equations. To simulate conduction from the right atrium to the bundle of His via the AV node, a stimulus was applied to the atrial myocytes at the start of the one-dimensional model. The pacing cycle length was 350 ms. After the 10th stimulus, the conduction time from the atrium to the bundle of His (at the end of the one-dimensional model) was measured. The computer programme was written using C++, with a message-passing interface (MPI) employed for parallel computation of action potentials using multiple CPUs and run on a personal computer with the macOS operating system.

**Computer modelling at the University of Montpellier**
The recently developed model of the mouse AV myocyte from Bartolucci et al.[21] was employed. The model accurately reproduces evoked as well as spontaneous action potentials of HCN4-expressing AV node myocytes and incorporates key ionic currents including HCN-mediated funny current ($I_f$), Cav1.2-mediated ($I_{Cav1.2}$) and Cav1.3-mediated ($I_{Cav1.3}$) L-type $Ca^{2+}$ currents, Cav3.1-mediated T-type $Ca^{2+}$ current ($I_{Ca,T}$), cardiac Nav1.5-mediated TTX-insensitive $Na^+$ current ($I_{Na}$), a TTX-sensitive $Na^+$ current, rapid delayed rectifier $K^+$ current ($I_{K,r}$) and ACh-activated $K^+$ current ($I_{K,ACh}$). To investigate the circadian rhythm in the conduction velocity of the AV node, a one-dimensional strand of 100 AV node myocytes was constructed. The length of a myocyte was assumed to be 65 μm. The time course of the action potential of myocyte *j* (j=1,….,100) was calculated by the following general partial differential equation:

$$C_m \frac{dV_{Myo(j)}}{dt} = -I_{ionMyo(j)} - I_{GapMyo(j)} + I_{st(j,t)}$$

where $C_m$ is the cell capacitance, $V_{Myo(j)}$ is the membrane potential of myocyte $j$, $t$ is the time, $I_{ionMyo(j)}$ is the sum of ionic currents of myocyte $j$, $I_{GapMyo(j)}$ is the sum of the coupling currents flowing through gap junctions between myocyte $j$ and the preceding $(j-1)$ and succeeding $(j+1)$ myocytes in the strand, and $I_{st(j,t)}$ is the myocyte-dependent and time-limited stimulus current. The $C_m$ was assumed to be 22 pF (Marger *et al.*[20] reported $C_m$ to be ~20 pF in mouse AV node myocytes). Myocytes 1-5 in the strand were paced at 350 beats/min with a 5 ms stimulus pulse of 200 pA amplitude. $I_{GapMyo(j)}$ was calculated according to the following equation:

$$I_{GapMyo} = D \frac{\left(V_{Myo(j+1)} + V_{Myo(j-1)} - V_{Myo(j)}\right)}{\Delta x^2}$$

where $D$ is the diffusion coefficient (assumed to be 7.68182E-05). The following boundary conditions were applied:

$$V_{Myo(2)} = V_{Myo(1)};\ V_{Myo(100)} = V_{Myo(99)}$$

The conductances for $I_{Cav1.2}$, $I_{Cav1.3}$, $I_{K,r}$, and $I_{K,ACh}$ were scaled at ZT0, 6, 12 and 18 according to the experimental data in Fig. 6. It was assumed that the conductances in the Bartolucci et al.[21] model correspond to ZT18 and the conductances were scaled according to this. Partial differential equations were integrated using the Method of Lines (http://www.scholarpedia.org/article/Method_of_lines; Dokos[28]). Simulations were run using C++ code first implemented by Michael Colman and David Conesa at the University of Leeds. The code was modified to allow the maximal conductance for an ionic current to change during the circadian period. Simulations were run for 5 s with a 0.01 ms time step. Simulation results were analysed using custom made Matlab (2025B, Mathworks Inc.) script automatically detecting and reporting the action potential frequency, action potential threshold, action potential peak, action potential duration at 90% repolarization, and maximum diastolic potential. The conduction velocity of the action potential along the strand was calculated by determining the distance (in cm) and time delay (in seconds) between the action potentials of myocytes 10 and 80. Simulation results were plotted using GraphPad Prism (version 10) or Matlab. The model code is available for download at Github public server with link: https://github.com/CARDIAC-RHYTHM-PHYSIOPATHOLOGY/Circadian_AVN_1D_Strand.

### SUPPLEMENTARY RESULTS AND DISCUSSION

#### Sensitivity and resolution of proteomics

In work on the mouse sinus node and AV node, we measured ~55,450[29] and 55,385[30] transcripts, respectively, using RNAseq. For comparison, a meta-analysis of gene expression in the whole normal human heart identified 43,360 transcripts.[31] The transcriptome consistently exceeds the proteome in complexity and size. In this study, using mass spectrometry-based proteomics we measured 4,186 proteins in the mouse AV node and previously we have measured 7,248 proteins in the mouse sinus node and atrial muscle[32] and in another study 6,849 proteins in the mouse sinus node.[33] 10,700 proteins have been measured in the human heart.[34] Part of the difference between the transcriptome and proteome is that not all transcripts are translated into unique proteins.[35] However, technical limitations in mass spectrometry-based proteomics also hinders detection of low-abundance or highly similar protein isoforms.[36] The abundance of cellular proteins spans many orders of magnitude from the least abundant to the most abundant and mass spectrometric instrumentation is unable to detect proteins over such a large range of abundances – whereas the dynamic range of the cellular proteome may approach seven orders of magnitude, the dynamic range of mass spectrometric instrumentation may exhibit only four orders of magnitude. This is a central reason why low-abundance proteins are missed.[36] It is likely that this is one reason why only 4,186 proteins were detected in the mouse AV node in this study. For example, it is known that certain proteins are present in the AV node and yet they were not detected in this study. This includes all circadian clock proteins, some ion channel proteins (e.g. Cav1.3) and most transcription factors.

**The role of ionic currents in the circadian rhythm in the PR interval**

In the main text, attention is focussed on the L-type $Ca^{2+}$ current, $I_{Ca,L}$, but other ionic currents are likely to be involved in the circadian rhythm in the PR interval as well. The increase in $I_{K,ACh}$ during the active period (Fig. 6H-K) is counter-intuitive if the increase in the AV node conduction velocity during the active period and consequently the circadian rhythm in the PR interval is the result of a decrease in vagal tone during the active period. If a circadian rhythm in vagal tone is responsible for the circadian rhythm in the PR interval, it would perhaps make more sense for $I_{K,ACh}$ to be reduced rather than increased during the active period.

It is interesting that $I_{K,ACh}$ was at a peak at ZT18 (Fig. 6H-K) at the same time as the AV node conduction velocity was at its maximum (and therefore the PR interval was at its shortest; e.g. Fig. 1B). The *Kcnj2* (Kir2.1) and *Kcnj12* (Kir2.2) transcripts responsible for the background inward rectifier $K^+$ current, $I_{K,1}$, were also at a maximum at about the same time (Fig. 4; which could explain the increase in the inward rectifying component of background current at ZT18 shown in Fig. S8). It is likely that it is the combination of $I_{K,1}$ and $I_{K,ACh}$ which is responsible for the resting potential (or maximum diastolic potential) of the AV node – the reversal potential of background current was ~-30 mV (Fig. S8A) whereas the reversal potential of $I_{K,ACh}$ was ~-80 mV (Fig. 6H) and the resting potential of the AV node is in between (~-60 mV[37]). Increases in $I_{K,1}$ and $I_{K,ACh}$ during the active period could increase the resting potential, and a more negative resting potential may be more suitable for fast conduction because there could be less inactivation of the inward currents, such as the $Na^+$ current, $I_{Na}$, responsible for conduction. Therefore, an increase in $K^+$ currents is expected to have two opposing effects: *per se* an increase is expected to slow conduction, but an increase is also expected to speed up conduction by hyperpolarizing the membrane and reducing inactivation of the inward currents underlying conduction. Could this interplay between the two opposing effects be responsible for the decrease in the amplitude of the circadian rhythm in the PR interval after knockout of *Kcnj5* and thus $I_{K,ACh}$ as shown in Fig. 1B.

$I_{Na}$ is a major determinant of the conduction velocity in the heart. Although the $Na^+$ channel, *Scn5a* (Nav1.5), is not expressed in the compact part of the AV node it is expressed in the transitional AN and NH regions,[18,38] conduction through which will help determine the PR interval. GWAS studies shows *Scn5a* to be associated with the PR interval (Table 1). It is possible, therefore, that $I_{Na}$ is involved with the circadian rhythm in the PR interval. In this study there was a significant circadian rhythm in the Scn5a transcript (Fig. 3), which peaked at the start of the active period, but not in the protein (Fig. 5) – however, the protein data have to be treated cautiously, because the heterogeneity of *Scn5a* expression in the AV node may cloud the picture. In the mouse ventricle, we have recently shown a circadian rhythm in *Scn5a* transcript expression (peaking at the start of the active period as in the mouse AV node – Fig. 3) and the density of $I_{Na}$ (albeit a complex one).[39]

Both the transcript and protein for *Kcnh2*, responsible for the rapid delayed rectifier $K^+$ current, $I_{K,r}$, showed a significant circadian rhythm peaking at the start of the active period (Figs. 4C and 5) - the same as we have shown for *Kcnh2* transcript in mouse ventricle.[39] Curiously though, $I_{K,r}$ in the mouse AV node showed a significant circadian rhythm but reached a nadir at the start of the active period (Fig. 6F), whereas in the mouse ventricle $I_{K,r}$ peaked at the start of the active period.[39] $I_{K,r}$ along with $I_{Ca,L}$ and $I_{K,ACh}$ (all of which showed significant circadian rhythms - Fig. 6) are important determinants of action potential duration. There is a circadian rhythm in the AV node refractory period in the human (measured during atrial fibrillation) and the refractory period is longest at night (sleep period).[40-42] This is likely to reflect a rhythm in action potential duration. $I_{K,r}$, $I_{Ca,L}$ and $I_{K,ACh}$ could be responsible, but an investigation of action potential duration was outside of the scope of the present study.

**Circadian rhythm in contractile proteins**

Myosin light chain kinase 3, also known as cardiac myosin light chain kinase (*Mylk3*), showed a significant rhythm and peaked at the start of the active period (Fig. S11). Cardiac myosin light chain kinase enhances myofilament $Ca^{2+}$ sensitivity by phosphorylating the myosin regulatory light chain.[43,44] Titin (*Ttn*), telethonin (*Tcap*) and alpha actinin 2 (*Actn2*) also showed significant circadian rhythms and again peaked at the start of the active period (Fig. S11; a circadian rhythm in *Tcap* in mouse heart has previously been reported[45]). Titin is a molecular spring that gives muscle its elasticity and provides structural support and stability within the sarcomere (Fig. 11). It interacts with actin and myosin and links the Z disk and M line of the sarcomere. Telethonin and alpha actinin 2 are in a complex anchoring titin at the Z disc (Fig. S11). There is evidence that alpha actinin 2 affects myofilament $Ca^{2+}$ sensitivity[46] and there is speculative evidence that telethonin and titin also influence myofilament $Ca^{2+}$ sensitivity.[47,48] Troponin I interacting kinase (*Tnni3k*) shows a significant circadian rhythm peaking at the start of the active period (Fig. S11) and it may again regulate myofilament $Ca^{2+}$ sensitivity (it phosphorylates cardiac troponin I and increases contractility[49] and phosphorylation is known to regulate myofilament $Ca^{2+}$ sensitivity[50]). Interestingly, *Tnni3k* is associated with AV node conduction.[51] Although the AV node is not primarily a contractile tissue, the findings above are novel and have implications for the whole heart. To investigate this, in mouse ventricle it was confirmed using RNAseq that there is an upregulation of *Actn2*, *Mylk3*, *Ttn* and *Tcap* (but not *Tnni3k*) transcripts during the active period (Fig. S12A) consistent with the protein findings in the AV node (Fig. S11). Podobed *et al.*[52,53] have shown that cardiac myofilament $Ca^{2+}$ sensitivity in the mouse heart does indeed increase during the active period and this effect is absent following disruption of the cardiomyocyte circadian clock by the expression of a cardiomyocyte-specific *Clock* mutant. An increase in myofilament $Ca^{2+}$ sensitivity during the active period is expected to have two effects: first, the greater binding of $Ca^{2+}$ to the myofilaments means the myofilaments will be a more effective $Ca^{2+}$ buffer and this should reduce the amplitude of the intracellular $Ca^{2+}$ transient; the intracellular $Ca^{2+}$ transient in ventricular myocytes (rat) has already been shown to be reduced during the active period.[54] Secondly, the greater binding of $Ca^{2+}$ to the myofilaments should result in an increase in the contraction and this has not been examined and rat ventricular myocytes were used to test this prediction. Myocytes isolated at ZT3.5 or ZT15.5 were stimulated at 1 Hz. As predicted, Fig. S12B shows that the mean twitch shortening was greater during the active period and Fig. S12C shows that the distribution of twitch shortening amongst myocytes was shifted to the left (to greater twitch shortening) during the active period. Although an increase in the contraction at a time when the intracellular $Ca^{2+}$ transient is decreased may be surprising, such a situation has already been reported in the case of a drug-induced increase in myofilament $Ca^{2+}$ sensitivity.[55] Stroke volume is enhanced at the start of the active period[56] (presumably to meet the expected demand for greater cardiac output) and it is logical for cardiac contractility to be greater at this time as shown here.

**Circadian rhythms in micro-RNAs**

A large number of micro-RNAs showed significant circadian rhythms (examples are shown in Fig. S19) and they could contribute to or be responsible for circadian rhythms in their target transcripts. It has already been stated that it is widely assumed that, when a micro-RNA rises, its target transcript must fall and *vice versa* and this inverse pattern is useful for target discovery (however, this is not necessarily the case[57] - it could be the result of a false positive prediction by IPA or even genuine and the result of another type of micro-RNA-mRNA interaction). miR-211-5p and miR-432 have been shown to regulate *Hcn4*[58] and both oscillated in antiphase to *Hcn4* as expected if they are involved in the oscillation of *Hcn4* (Fig. S20). Overexpression of miR-1 causes AV block[59] and miR-1 is known to regulate both *Scn5a*[60] and *Cacna1c* (Cav1.2)[59] and Fig. S20 shows that miR-1a-3p oscillated in antiphase to both transcripts. miR-29 and miR-652-3p have been implicated in regulating *Kcnj12* (Kir2.2)[61] and *Kcnn3* (SK3),[62] respectively, and both oscillated in antiphase to the respective transcripts (Fig. S20). Micro-RNAs were not only potentially linked to ion channel transcripts, they were also potentially linked to the circadian clock. *Per2* is a key regulatory point in the circadian clock[63] and miR-25-3p has been shown to regulate *Per2*[64] and miR-138-1-3p and let-7i-3p are predicted to target *Per2* (Supplementary files S4 and S5) and Fig. S20 shows that all the micro-RNAs oscillated roughly in antiphase to *Per2*. Another clock transcript, *Rora*, is a candidate regulated by miR-99a-3p[65] and the two oscillated in antiphase (Fig. S20). Fig. S20 shows that micro-RNAs could contribute to the circadian rhythms in transcripts.

Supplementary Files S4 and S5 summarise the potential ion channel, clock and transcription factor target transcripts (as identified by QIAGEN Ingenuity Pathway Analysis, IPA) shown to be rhythmic in this study by micro-RNAs also shown to be rhythmic in this study. The micro-RNAs generally potentially targeted more than one transcript (some potentially targeted seven) and a single micro-RNA generally potentially targeted different types of transcript (clock, ion channel and transcription factor) (Supplementary File S4). A single transcript was generally potentially targeted by more than one micro-RNA: two clock transcripts *Clock* and *Per2* were potentially targeted by 12 and seven micro-RNAs, respectively, while *Kcnj3* (Kir3.1) was potentially targeted by eight micro-RNAs (Supplementary File S5). Could the number of micro-RNAs targeting a transcript reflect the importance of the transcript?

IPA was also used to identify micro-RNAs potentially targeting the proteins shown in this study to have a circadian rhythm (in this case, temporal relationship of micro-RNAs and proteins was not considered because proteins are downstream of transcripts targeted by micro-RNAs). As already discussed, many of the potentially targeted rhythmic proteins are involved in metabolism. Micro-RNAs potentially target other types of protein – for example, rhythmic miR-327 is predicted to regulate rhythmic *Mylk3* (myosin light chain kinase 3) discussed above in relation to contractile proteins and illustrated in Fig. S11. Whereas the majority of the proteins with significant circadian rhythms peaked at the start of the active period, the micro-RNAs with significant rhythms peaked at different times (Fig. S21).

### SUPPLEMENTARY FILES

**Supplementary File S1.** Excel spreadsheet showing all transcripts detected, the P value for a significant circadian rhythm and the acrophase.

**Supplementary File S2.** Excel spreadsheet showing all proteins detected, the P value for a significant circadian rhythm and the acrophase.

**Supplementary File S3.** Excel spreadsheet showing all micro-RNAs measured, the P value for a significant circadian rhythm and the acrophase.

**Supplementary File S4.** Potential targets of rhythmic micro-RNAs (micro-RNA focussed).

**Supplementary File S5.** Potential targets of rhythmic micro-RNAs (mRNA focussed).

53 Podobed, P., Pyle, W. G., Ackloo, S., Alibhai, F. J., Tsimakouridze, E. V., Ratcliffe, W. F., Mackay, A., Simpson, J. A., Wright, D. C., Kirby, G., Young, M. E. & Martino, T. A. The

day/night proteome in the murine heart. *American Journal of Physiology-Regulatory, Integrative and Comparative Physiology* **307**, R121–R137 (2014). https://doi.org/10.1152/ajpregu.00011.2014
54 Collins, H. E. & Rodrigo, G. C. Inotropic response of cardiac ventricular myocytes to beta-adrenergic stimulation with isoproterenol exhibits diurnal variation: involvement of nitric oxide. *Circulation Research* **106**, 1244–1252 (2010). https://doi.org/10.1161/CIRCRESAHA.109.213942
55 White, J., Lee, J. A., Shah, N. & Orchard, C. H. Differential effects of the optical isomers of EMD 53998 on contraction and cytoplasmic $Ca^{2+}$ in isolated ferret cardiac muscle. *Circulation Research* **73**, 61–70 (1993). https://doi.org/10.1161/01.res.73.1.61
56 Veerman, D. P., Imholz, B. P. M., Wieling, W., Wesseling, K. H. & van Montfrans, G. A. Circadian Profile of Systemic Hemodynamics. *Hypertension* **26**, 55–59 (1995). https://doi.org/10.1161/01.HYP.26.1.55
57 Lopez-Gomollon, S., Mohorianu, I., Szittya, G., Moulton, V. & Dalmay, T. Diverse correlation patterns between microRNAs and their targets during tomato fruit development indicates different modes of microRNA actions. *Planta* **236**, 1875–1887 (2012). https://doi.org/10.1007/s00425-012-1734-7
58 Mesirca, P., Nakao, S., Nissen, S. D., Forte, G., Anderson, C., Trussell, T., Li, J., Cox, C., Zi, M., Logantha, S., Yaar, S., Cartensen, H., Bidaud, I., Stuart, L., Soattin, L., Morris, G. M., da Costa Martins, P. A., Cartwright, E. J., Oceandy, D., Mangoni, M. E., Jespersen, T., Buhl, R., Dobrzynski, H., Boyett, M. R. & D'Souza, A. Intrinsic electrical remodeling underlies atrioventricular block in athletes. *Circulation Research* **129**, e1–e20 (2021). https://doi.org/10.1161/circresaha.119.316386
59 Zhang, Y., Sun, L., Zhang, Y., Liang, H., Li, X., Cai, R., Wang, L., Du, W., Zhang, R., Li, J., Wang, Z., Ma, N., Wang, X., Du, Z., Yang, B., Gao, X. & Shan, H. Overexpression of microRNA-1 causes atrioventricular block in rodents. *Int J Biol Sci* **9**, 455–462 (2013). https://doi.org/10.7150/ijbs.4630
60 Boxhammer, E., Dienhart, C., Rezar, R., Hoppe, U. C. & Lichtenauer, M. Deciphering the Role of microRNAs: Unveiling Clinical Biomarkers and Therapeutic Avenues in Atrial Fibrillation and Associated Stroke-A Systematic Review. *Int J Mol Sci* **25** (2024). https://doi.org/10.3390/ijms25105568
61 Xiao-Jun, Z., Cai-Xiu, L., Kai-Jun, S., Lei-Ling, L. & Dan-Yan, X. A soluble epoxide hydrolase inhibitor upregulated KCNJ12 and KCNIP2 by downregulating microRNA-29 in a mouse model of myocardial infarction. *Heart Surgery Forum* **23**, 579–585 (2020).
62 Zhu, Q. L., Zhan, D. M., Chong, Y. K., Ding, L. & Yang, Y. G. MiR-652-3p promotes bladder cancer migration and invasion by targeting KCNN3. *European Review for Medical and Pharmacological Sciences* **23**, 8806–8812 (2019). https://doi.org/10.26355/eurrev_201910_19275
63 Wilkins, A. K., Barton, P. I. & Tidor, B. The Per2 negative feedback loop sets the period in the mammalian circadian clock mechanism. *PLOS Computational Biology* **3**, e242 (2007). https://doi.org/10.1371/journal.pcbi.0030242
64 Park, I., Kim, D., Kim, J., Jang, S., Choi, M., Choe, H., Choe, Y. & Kim, K. microRNA-25 as a novel modulator of circadian Period2 gene oscillation. *Exp Mol Med* **52**, 1614–1626 (2020). https://doi.org/10.1038/s12276-020-00496-5
65 Okada, R., Koshizuka, K., Yamada, Y., Moriya, S., Kikkawa, N., Kinoshita, T., Hanazawa, T. & Seki, N. Regulation of oncogenic targets by miR-99a-3p (passenger strand of miR-99a-duplex) in head and neck squamous cell carcinoma. *Cells* **8** (2019). https://doi.org/10.3390/cells8121535
66 Hayano, J., Ishihara, S., Fukuta, H., Sakata, S., Mukai, S., Ohte, N. & Kimura, G. Circadian rhythm of atrioventricular conduction predicts long-term survival in patients with chronic atrial fibrillation. *Chronobiol Int* **19**, 633–648 (2002). https://doi.org/10.1081/cbi-120004223
67 Yagita, K. & Okamura, H. Forskolin induces circadian gene expression of rPer1, rPer2 and dbp in mammalian rat-1 fibroblasts. *FEBS Letters* **465**, 79–82 (2000). https://doi.org/https://doi.org/10.1016/S0014-5793(99)01724-X
68 Hayashi, T., Arimura, T., Itoh-Satoh, M., Ueda, K., Hohda, S., Inagaki, N., Takahashi, M., Hori, H., Yasunami, M., Nishi, H., Koga, Y., Nakamura, H., Matsuzaki, M., Choi, B. Y., Bae, S. W., You, C. W., Han, K. H., Park, J. E., Knöll, R., Hoshijima, M., Chien, K. R. & Kimura,

A. Tcap gene mutations in hypertrophic cardiomyopathy and dilated cardiomyopathy. *Journal of the American College of Cardiology* **44**, 2192–2201 (2004). https://doi.org/https://doi.org/10.1016/j.jacc.2004.08.058

**Table S1. List of the ion channel transcripts investigated and their significance of a circadian rhythm**. The transcripts are listed according to the P value. Pink, P<0.05; orange, 0.1>P>0.05; blue, P>0.1.

| Transcript | Common name | P value | Acrophase (ZT, h) |
|---|---|---|---|
| *Kcnh2* | ERG | 1.84706E-08 | 13.9 |
| *Hcn4* | HCN4 | 4.46047E-08 | 20.3 |
| *Kcnj12* | Kir2.2 | 9.50905E-07 | 15.2 |
| *Cacna1d* | Cav1.3 | 1.19533E-05 | 16.7 |
| *Gja5* | Cx40 | 0.00011 | 5.8 |
| *Hcn1* | HCN1 | 0.00017 | 17.1 |
| *Kcnd2* | Kv4.2 | 0.00031 | 6.1 |
| *Kcnj5* | Kir3.4 | 0.00056 | 16.0 |
| *Abcc8* | SUR1 | 0.0012 | 9.6 |
| *Clcn2* | $Cl^-$ channel 2 | 0.0013 | 13.6 |
| *Kcna5* | Kv1.5 | 0.0019 | 9.9 |
| *Kcna2* | Kv1.2 | 0.0020 | 22.4 |
| *Gja1* | Cx43 | 0.0032 | 4.2 |
| *Kcnj3* | Kir3.1 | 0.0034 | 11.4 |
| *Scn5a* | Na1.5 | 0.0037 | 11.0 |
| *Scn1a* | Nav1.1 | 0.0044 | 13.4 |
| *Cacna1g* | Cav3.1 | 0.0068 | 21.6 |
| *Kcnn3* | SK3 | 0.011 | 21.4 |
| *Kcnq1* | KvLQT1 | 0.014 | 13.0 |
| *Cacna1c* | Cav1.2 | 0.025 | 14.0 |
| *Kcna4* | Kv1.4 | 0.025 | 17.0 |
| *Kcnj2* | Kir2.1 | 0.038 | 20.0 |
| *Hcn2* | HCN2 | 0.062 | 10.1 |
| *Kcnn2* | SK2 | 0.070 | 13.2 |
| *Cacna2d1* | Cavα2δ1 | 0.075 | 18.3 |
| *Cacna2d2* | Cavα2δ2 | 0.081 | 14.7 |
| *Kcnn1* | SK1 | 0.091 | 19.2 |
| *Kcnj8* | Kir6.1 | 0.099 | 11.2 |
| *Kcnj11* | Kir6.2 | 0.12 | - |
| *Kcnk3* | TASK-1 | 0.12 | - |
| *Cftr* | CFTR | 0.16 | - |
| *Kcnd3* | Kv4.3 | 0.16 | - |
| *Trpc3* | TRPC3 | 0.21 | - |
| *Cacnb2* | Cavβ2 | 0.22 | - |
| *Abcc9* | SUR2 | 0.37 | - |
| *Kcnb1* | Kv2.1 | 0.36 | - |
| *Gjc3* | Cx29; Cx30.2; Cx31.3 | 0.39 | - |
| *Scn1b* | Navβ1 | 0.49 | - |
| *Gjc1* | Cx45 | 0.55 | - |
| *Kcnj4* | Kir2.3 | 0.63 | - |
| *Kcnj14* | Kir2.4 | 0.66 | - |
| *Kcnip2* | KChIP2 | 0.69 | - |
| *Cacna1h* | Cav3.2 | 0.86 | - |

**Table S2. List of the intracellular $Ca^{2+}$-handling transcripts investigated and their significance of a circadian rhythm.** The transcripts are listed according to the P value. Pink, P<0.05; orange, 0.1>P>0.05; blue, P>0.1.

| Transcript | Common name | P value | Acrophase (ZT, h) |
|---|---|---|---|
| *Pln* | Phospholamban | 0.0039 | 9.9 |
| *Ryr3* | Ryanodine receptor 3 | 0.013 | 13.3 |
| *Sln* | Sarcolipin | 0.020 | 12.0 |
| *Ryr2* | Ryanodine receptor 2 | 0.074 | 10.5 |
| *Casq2* | Calsequestrin 2 | 0.098 | 17.2 |
| *Atp1a1* | $Na^+$-$K^+$ pump-$\alpha$1 | 0.12 | - |
| *Atp1a2* | $Na^+$-$K^+$ pump-$\alpha$2 | 0.32 | - |
| *Atp2a2* | SERCA2a | 0.34 | - |
| *Atp1a3* | $Na^+$-$K^+$ pump-$\alpha$3 | 0.40 | - |
| *Slc8a1* | NCX1; $Na^+$-$Ca^{2+}$ exchanger | 0.53 | - |

**Table S3 (over the page). List of protein kinases (proteins) detected showing a significant circadian rhythm.** *Prkag1* and *Prkar2* are genes for regulatory subunits of AMP-activated protein kinase and protein kinase A, respectively, rather than catalytic subunits, but they are still considered here. The protein kinases are listed according to the P value.

**Table S3 continued.**

| Protein | Comon name | P value | Acrophase (ZT, h) |
|---|---|---|---|
| PDK4 | Pyruvate dehydrogenase kinase, isozyme 4 | 0.0019 | 10.3 |
| PDPK1 | 3-phosphoinositide dependent protein kinase 1 | 0.0022 | 9.1 |
| CAMK2A | $Ca^{2+}$/calmodulin dependent protein kinase II$\alpha$ | 0.0036 | 9.9 |
| OBSCN | Obscurin | 0.0052 | 9.3 |
| EPHA4 | EPH receptor A4 | 0.0072 | 9.2 |
| ALPK2 | Alpha Kinase 2 | 0.0079 | 10.4 |
| SPEG | Striated muscle enriched protein kinase | 0.0087 | 9.8 |
| CDK9 | Cyclin dependent kinase 9 | 0.0095 | 8.9 |
| CAMK2D | $Ca^{2+}$/calmodulin dependent protein kinase II$\delta$ | 0.011 | 10.2 |
| DYRK1A | Dual specificity tyrosine-phosphorylation-regulated kinase 1A | 0.011 | 9.9 |
| TRIM24 | Tripartite Motif Containing 24 | 0.012 | 7.0 |
| MYLK3 | Myosin light chain kinase 3 | 0.013 | 10.2 |
| CSNK1D | Casein kinase 1$\delta$ | 0.017 | 9.3 |
| CDC42BPA | CDC42 binding protein kinase $\alpha$ | 0.018 | 9.4 |
| PRKAA2 | 5'-AMP-activated protein kinase catalytic subunit $\alpha$2 | 0.019 | 10.1 |
| CAMK1D | $Ca^{2+}$/calmodulin dependent protein kinase ID | 0.020 | 12.0 |
| MAP4K3 | Mitogen-activated protein kinase kinase kinase kinase 3 | 0.020 | 11.2 |
| MAP4K5 | Mitogen-activated protein kinase kinase kinase kinase 5 | 0.021 | 13.6 |
| ILK | Integrin linked kinase | 0.025 | 10.7 |
| EPHA7 | EPH receptor A7 | 0.027 | 9.0 |
| PTK2 | Protein tyrosine kinase 2 | 0.028 | 10.6 |
| PAK4 | P21 (RAC1)-activated kinase 4 | 0.030 | 12.1 |
| CAMK1 | $Ca^{2+}$-calmodulin dependent protein kinase I | 0.032 | 12.2 |
| SBK2 | SH3 domain binding kinase family member 2 | 0.032 | 9.4 |
| TTN | Titin | 0.032 | 9.1 |
| NPR2 | Natriuretic peptide receptor 2 | 0.037 | 8.4 |
| PDK1 | Pyruvate dehydrogenase kinase, isozyme 1 | 0.038 | 10.1 |
| PRKAG1 | 5'-AMP-activated protein kinase subunit $\gamma$1 | 0.039 | 9.1 |
| RIOK3 | RIO kinase 3 | 0.039 | 10.0 |
| TNNI3K | TNNI3 interacting kinase | 0.039 | 11.1 |
| PRKAR2B | cAMP-dependent protein kinase type II, $\beta$ regulatory subunit | 0.040 | 11.1 |
| PRKG1 | cGMP-dependent protein kinase 1, $\alpha$ isozyme | 0.042 | 12.8 |
| TBCK | TBC1 domain containing kinase | 0.042 | 8.3 |
| CSNK1A1 | Casein kinase 1 $\alpha$1 | 0.045 | 11.2 |
| ALPK3 | Alpha kinase 3 | 0.046 | 9.5 |
| TIE1 | Tyrosine kinase with immunoglobulin like and EGF like domains 1 | 0.048 | 11.4 |
| MAPK12 | 1. p38$\gamma$ 2. Mitogen-activated protein kinase 12 | 0.049 | 11.4 |

**Table S4. List of transcription factors (either transcripts or proteins) detected and their significance of a circadian rhythm.** Transcripts and proteins are listed according to the P value. There is no overlap between the transcripts and proteins because of the limited resolution of mass spectrometry and only selected transcription factors were measured at the transcript level. The transcripts and proteins are listed according to the P value. Pink, P<0.05; orange, 0.1>P>0.05; blue, P>0.1.

| Transcript or protein | Common name | P value | Acrophase (ZT, h) |
|---|---|---|---|
| | **TRANSCRIPT** | | |
| *Tef* | Thyrotroph embryonic factor | 2.3068E-34 | 12.6 |
| *Klf15* | Krüppel-like factor 15 | 2.87E-11 | 23.5 |
| *Shox2* | Short-stature homeobox 2 | 2.06E-07 | 17.5 |
| *Tbx3* | T-Box transcription factor 3 | 3.15727E-07 | 20.0 |
| *Pitx2* | Paired-like homeodomain transcription factor 2 | 8.97E-05 | 19.2 |
| *Mef2c* | Myocyte enhancer factor 2C | 0.0029 | 5.5 |
| *Klf4* | Krüppel-like factor 4 | 0.0057 | 20.0 |
| *Tbx5* | T-Box transcription factor 5 | 0.0062 | 11.3 |
| *Nkx2-5* | Homeobox-containing transcription factor | 0.022 | 19.3 |
| *Lhx1* | LIM homeobox 1 | 0.025 | 19.3 |
| *Srf* | Serum response factor | 0.056 | 16.9 |
| *Tbp* | TATA-binding protein | 0.13 | - |
| *Hlf* | Hepatic leukemia factor | 0.2 | - |
| *Isl1* | Insulin gene enhancer protein | 0.23 | - |
| *Rest* | RE1-Silencing Transcription factor | 0.34 | - |
| *Gata6* | GATA-binding factor 6 | 0.36 | - |
| *Bhlhe40* | BHLHE40 | 0.41 | - |
| *Tbx18* | T-box transcription factor 18 | 0.52 | - |
| *Dbp* | D site of albumin promoter (albumin D-box) binding protein | 0.59 | |
| *Hand2* | Heart- and neural crest derivatives-expressed protein 2 | 0.59 | - |
| *Bhlhe41* | BHLHE41 | 0.63 | - |
| | **PROTEIN** | | |
| ZBTB20 | Zinc finger and BTB domain-containing protein 20 | 0.00039 | 6.6 |
| PBRM1 | Protein polybromo-1 | 0.022 | 21.3 |
| ESRRG | Estrogen-related receptor $\gamma$ | 0.025 | 9.4 |
| MTA2 | Metastasis associated 1 family member 2 | 0.031 | 21.3 |
| CARHSP1 | $Ca^{2+}$-regulated heat stable protein 1 | 0.039 | 12.3 |
| SMAD5 | SMAD family member 5 | 0.052 | 22.4 |
| NFIB | Nuclear Factor IB | 0.061 | 22.1 |
| NR3C1 | Nuclear receptor subfamily 3 group C member 1 | 0.073 | 21.6 |
| REL | REL proto-oncogene, NF-kB subunit | 0.10 | - |
| NFKB2 | Nuclear factor kappa B subunit 2 | 0.11 | - |
| TSC22D2 | TSC22 domain family member 2 | 0.11 | - |
| GATAD2B | GATA zinc finger domain containing 2B | 0.12 | - |
| RELA | RELA proto-oncogene, NF-kB subunit | 0.12 | - |
| STAT5B | Signal transducer and activator of transcription 5B | 0.13 | - |

**Table S4 continued.**

| Transcript or protein | Common name | P value | Acrophase (ZT, h) |
|---|---|---|---|
| | **PROTEIN** | | |
| ADNP | Activity-dependent neuroprotector homeobox | 0.14 | - |
| MBD2 | Methyl-CpG binding domain protein 2 | 0.14 | - |
| DNAJC1 | DnaJ heat shock protein family (Hsp40) member C1 | 0.16 | |
| BZW1 | Basic leucine zipper and W2 domains 1 | 0.17 | - |
| MECP2 | Methyl-CpG-binding protein 2 | 0.19 | - |
| MBD1 | Methyl-CpG binding domain protein 1 | 0.20 | - |
| NOTCH2 | Notch receptor 2 | 0.20 | |
| HMGB1 | High mobility group box 1 | 0.21 | - |
| STAT3 | Signal transducer and activator of transcription 3 | 0.21 | - |
| CSDE1 | Cold shock domain containing E1 | 0.22 | - |
| MTA1 | Metastasis associated 1 | 0.23 | - |
| CUX1 | Cut-like homeobox 1 | 0.24 | - |
| SP100 | SP100 nuclear antigen/nuclear body protein | 0.24 | - |
| ATF1 | Activating transcription factor 1 | 0.27 | - |
| GTF2I | General transcription factor IIi | 0.28 | - |
| NR2F2 | Nuclear receptor subfamily 2 group F member 2 | 0.32 | - |
| FOXK1 | Forkhead box K1 | 0.34 | - |
| STAT1 | Signal transducer and activator of transcription 1 | 0.34 | - |
| TSC22D4 | TSC22 domain family member 4 | 0.35 | - |
| UBTF | Upstream binding transcription factor | 0.38 | - |
| YBX1 | Y-box binding protein 1 | 0.41 | - |
| SMARCE1 | SWI/SNF related BAF chromatin remodeling complex subunit E1 | 0.42 | - |
| NFKB1 | Nuclear factor kappa B subunit 1 | 0.43 | - |
| SSRP1 | Structure-specific recognition protein 1 | 0.43 | - |
| STAT6 | Signal transducer and activator of transcription 6 | 0.45 | - |
| MTA3 | Metastasis associated 1 family member 3 | 0.54 | - |
| SUB1 | SUB1 regulator of transcription | 0.57 | - |
| STAT5A | Signal transducer and activator of transcription 5A | 0.60 | - |
| GABPA | GA binding protein transcription factor subunit alpha | 0.67 | - |
| CTCF | CCCTC-binding factor | 0.71 | - |
| HMGB3 | High mobility group box 3 | 0.72 | - |
| SETDB1 | SET domain bifurcated histone lysine methyltransferase 1 | 0.72 | - |
| NFIX | Nuclear factor IX | 0.76 | - |
| TFAM | Transcription factor A, mitochondrial | 0.80 | - |
| HOPX | HOP homeobox | 0.88 | - |
| SMARCC2 | SWI/SNF related, matrix associated, actin dependent regulator of chromatin, subfamily a, member 2 | 0.98 | - |
| NFIC | Nuclear factor IC | 0.99 | - |

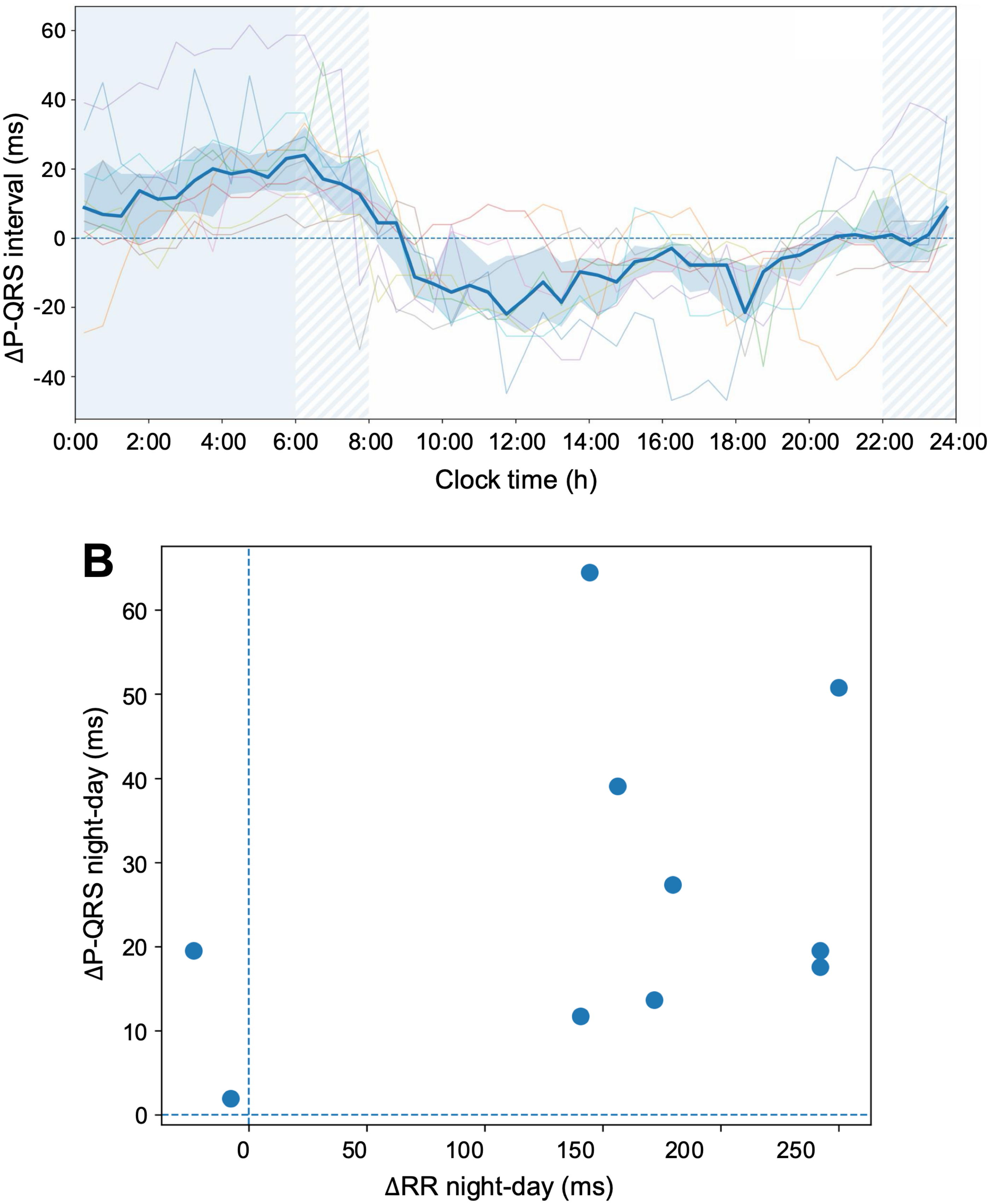


**Fig. S1. Circadian rhythm in the AV node in the human. A**, PR interval (shown relative to the mesor) of individual subjects over 24 h as well as the cohort median (dark blue line) and interquartile range (blue shaded area). Light blue background, sleep period (night); hatched blue background, transition between sleep and awake periods; colourless background, awake period (day). **B**, relationship between the nocturnal change in the PR interval and the nocturnal change in the RR interval in each subject. Most subjects are in the upper-right quadrant, but PR interval prolongation was also present in subjects without nocturnal RR prolongation.

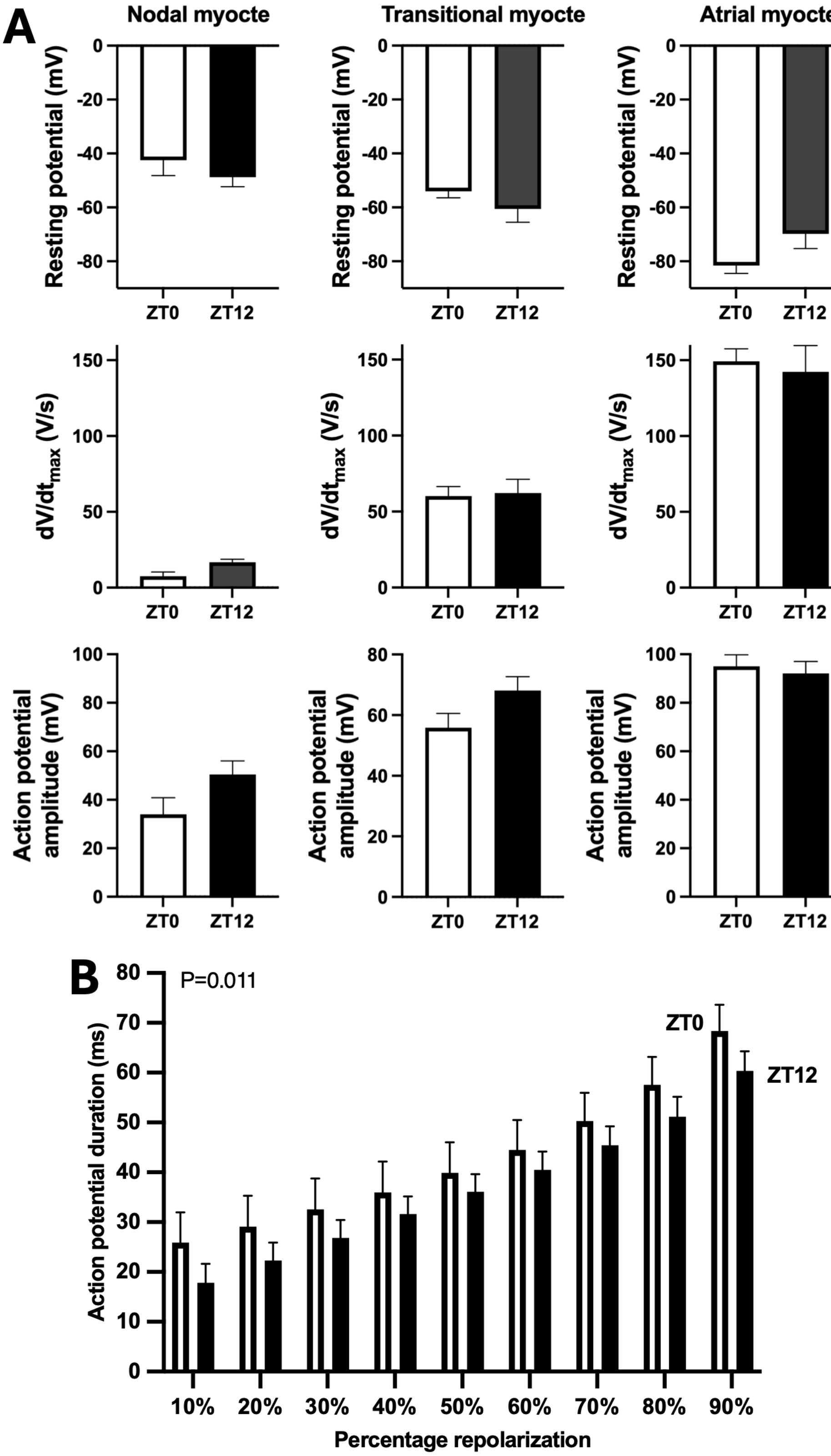


**Fig. S2. Intracellular AP recording from the mouse AV node. A**, classification of intracellular APs recorded from the right atrial preparation. If a microelectrode impalment appeared complete and the AP recording stable, APs were classified according to the maximum upstroke velocity, $dV/dt_{max}$, of the AP: <25 V/s, nodal AP; 25-100 V/s, transitional AP; >100 V/s, atrial AP. The graphs show the mean (±SEM) maximum diastolic potential, $dV/dt_{max}$ and the AP amplitude in the three AP types at ZT0 and 12 (n=5-9). As well as an increase in $dV/dt_{max}$, there is an increase in the maximum diastolic potential and AP amplitude from the nodal to the transitional to the atrial AP type as expected.[26] **B**, day-night difference in the duration of the nodal AP (mean±SEM, n=6 at ZT0 and n=8 at ZT12). AP duration is shown at 10-90% repolarization at ZT0 and 12. The P value is shown (two-way ANOVA). The AV node AP duration is longer during the sleep period (beginning at ZT0) and this is consistent with evidence of a longer AV node refractory period during the sleep period (night) in the human.[66]

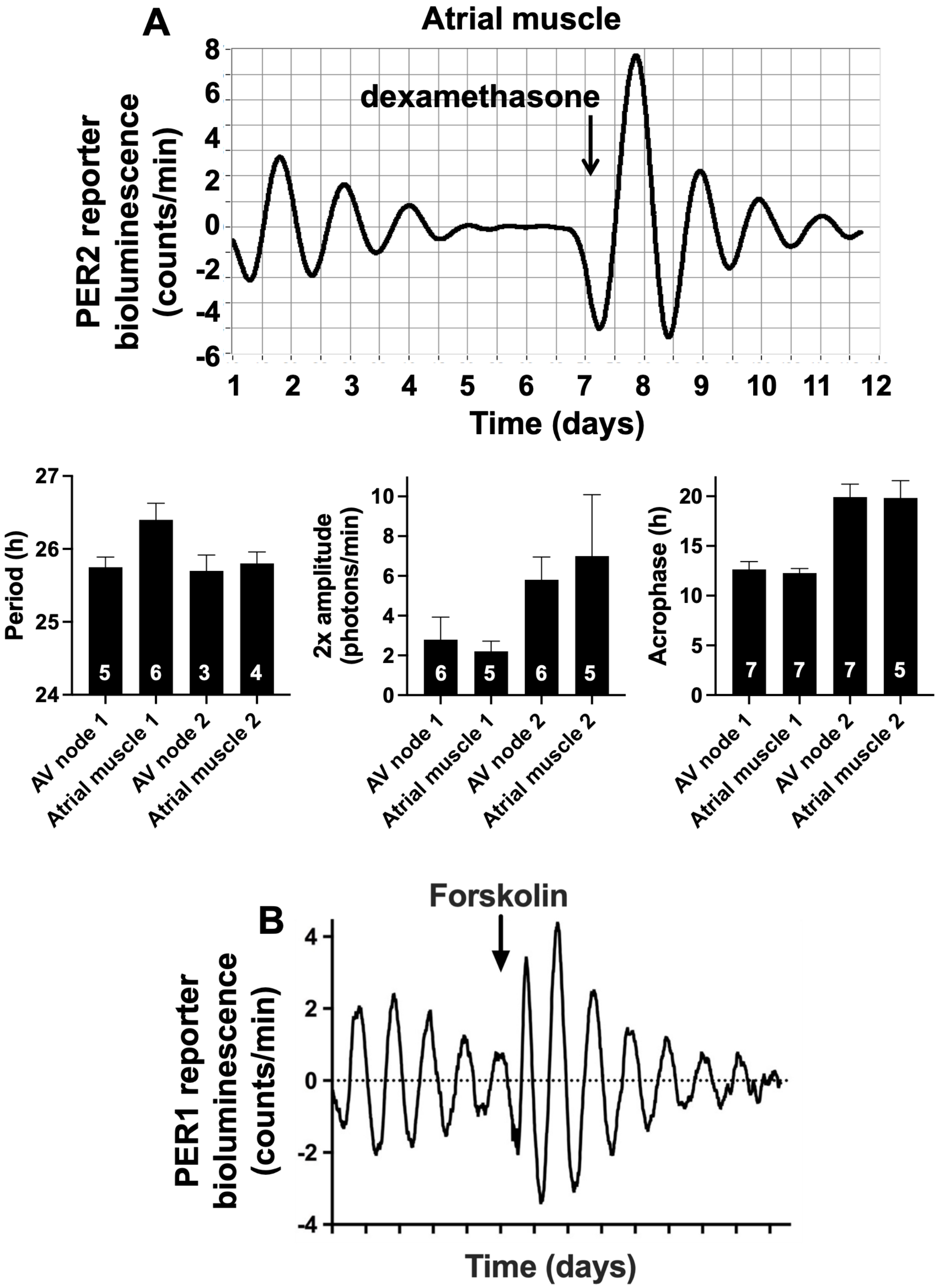


**Fig. S3. Evidence of a functioning circadian clock. A**, PER2 expression (reported by luciferase bioluminescence). Top, example of PER2 reporter bioluminescence in atrial muscle from a male PER2::LUCIFERASE mouse. PER2 expression varied in a circadian manner and when the circadian rhythm had run down it was reinstated by 0.1 mM dexamethasone. Bottom, characteristics of the circadian rhythm in PER2 (reported by luciferase bioluminescence) both in the AV node (Fig. 2C shows an example trace) and atrial muscle before (AV node 1 and atrial muscle 1) and after (AV node 2 and atrial muscle 2) application of dexamethasone; data from a mixture of male and female mice. The graphs show the mean±SEM period (peak to peak time), 2x amplitude (trough to peak) and acrophase (ZT at peak expression) of the circadian rhythm (n within bars). **B**, PER1 expression (reported by luciferase bioluminescence) in the isolated AV node from a male PER1::LUCIFERASE mouse. PER1 activity varied in a circadian manner. When the circadian rhythm had run down, it was reinstated by 10 μM forskolin. Forskolin is known to amplify circadian rhythms via the clock gene, *Per2* (e.g. Yagita and Okamura[67]). Data in A obtained at the University of Liverpool and data in B obtained at the University of Manchester.

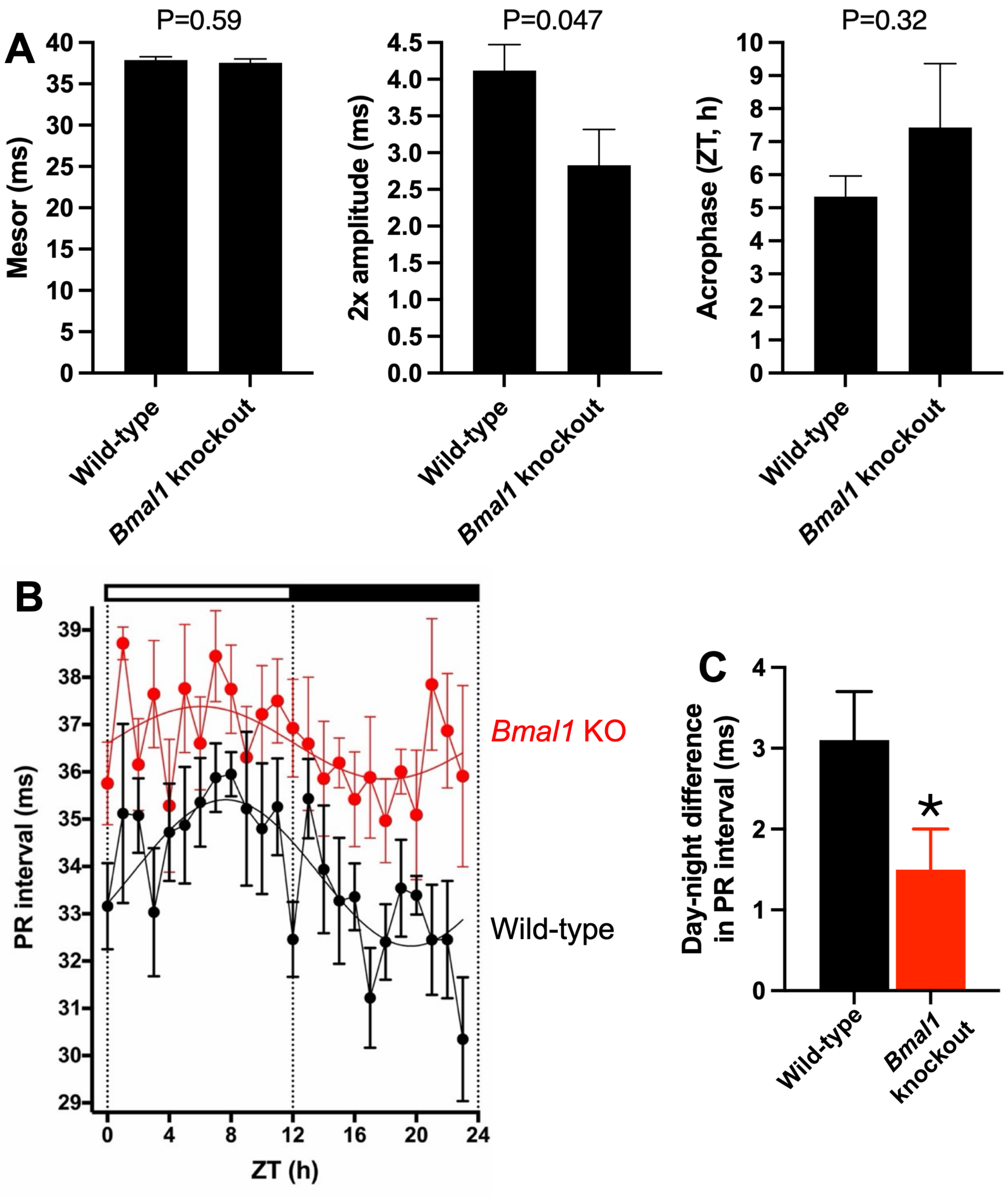


**Fig. S4. Knockout of the clock gene, *Bmal1*, reduces the amplitude of the circadian rhythm in the PR interval. A**, mean±SEM mesor, 2x amplitude and acrophase of the circadian rhythms in the PR interval in nine wild-type and 11 *Bmal1* knockout mice (based on cosinor analysis of individual mice). The difference between the wild-type and knockout mice was tested using an unpaired t test and the P value is shown. **B**, mean±SEM PR interval measured using telemetry from three wild-type mice and three *Bmal1* knockout mice. **C**, mean±SEM day-night difference in the PR interval in the wild-type and *Bmal1* knockout mice in panel B. *P<0.05 (unpaired t test).

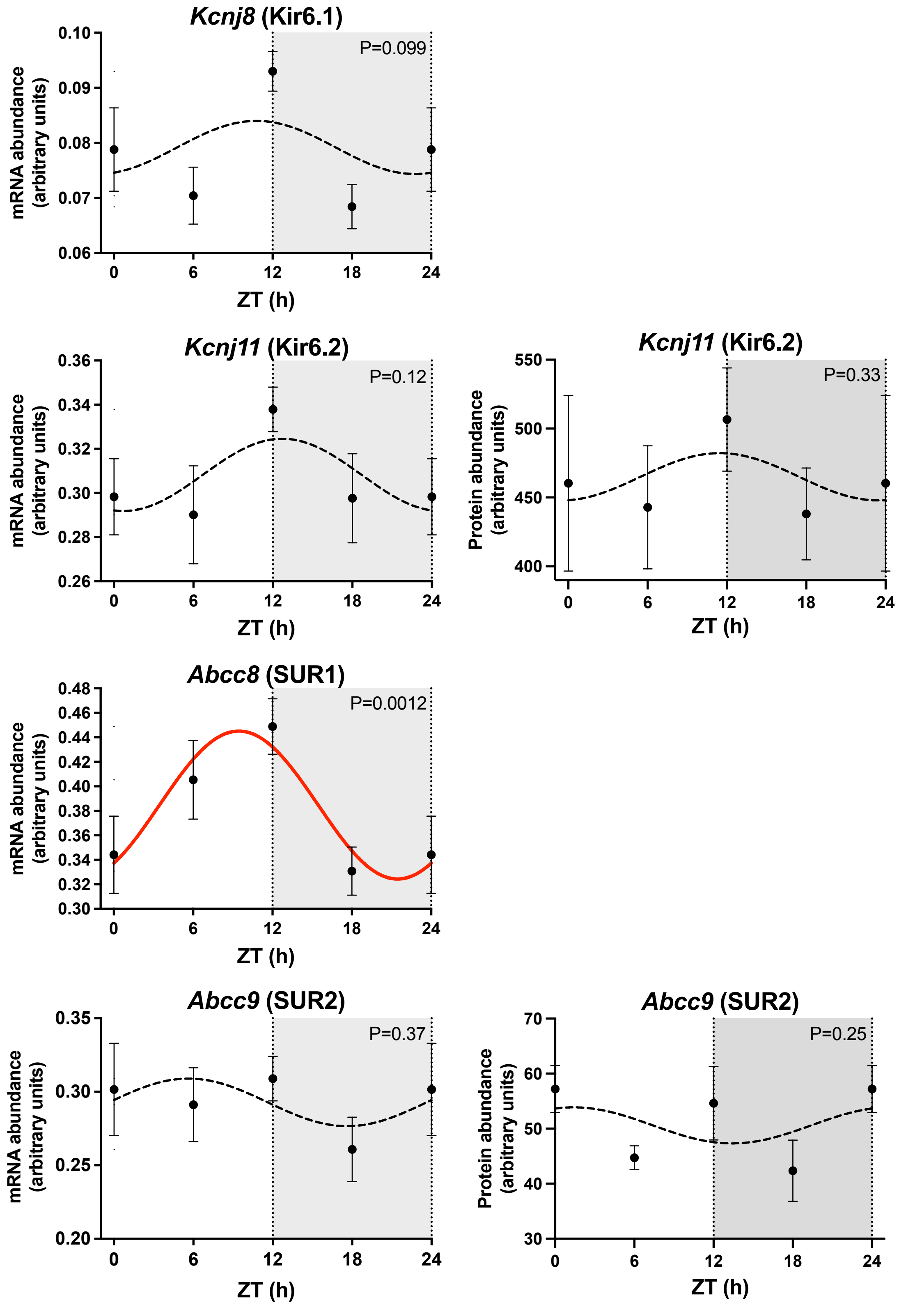


**Fig. S5. Circadian rhythm in the ATP-sensitive $K^+$ channel.** The graphs show the abundance of four transcripts and two proteins (remaining two proteins not detected) responsible for the ATP-sensitive $K^+$ channel measured at four time points over 24 h. Data from 9-11 mice and 4 or 5 biological replicates (from 8 or 10 mice) in the case of transcript and protein measurements, respectively. In this and similar figures: gene name and common name (in parenthesis) given; mean±SEM transcript or protein abundance shown at four ZT time points over 24 h (ZT24 data repeat of ZT0 data); hours of darkness shaded; P value from cosinor analysis for significance of a day-night rhythm given; significant transcripts or proteins (P<0.05) fitted with a cosine wave shown as a red line by a least squares fitting method; non-significant transcripts fitted with a cosine wave shown as a black dashed line.

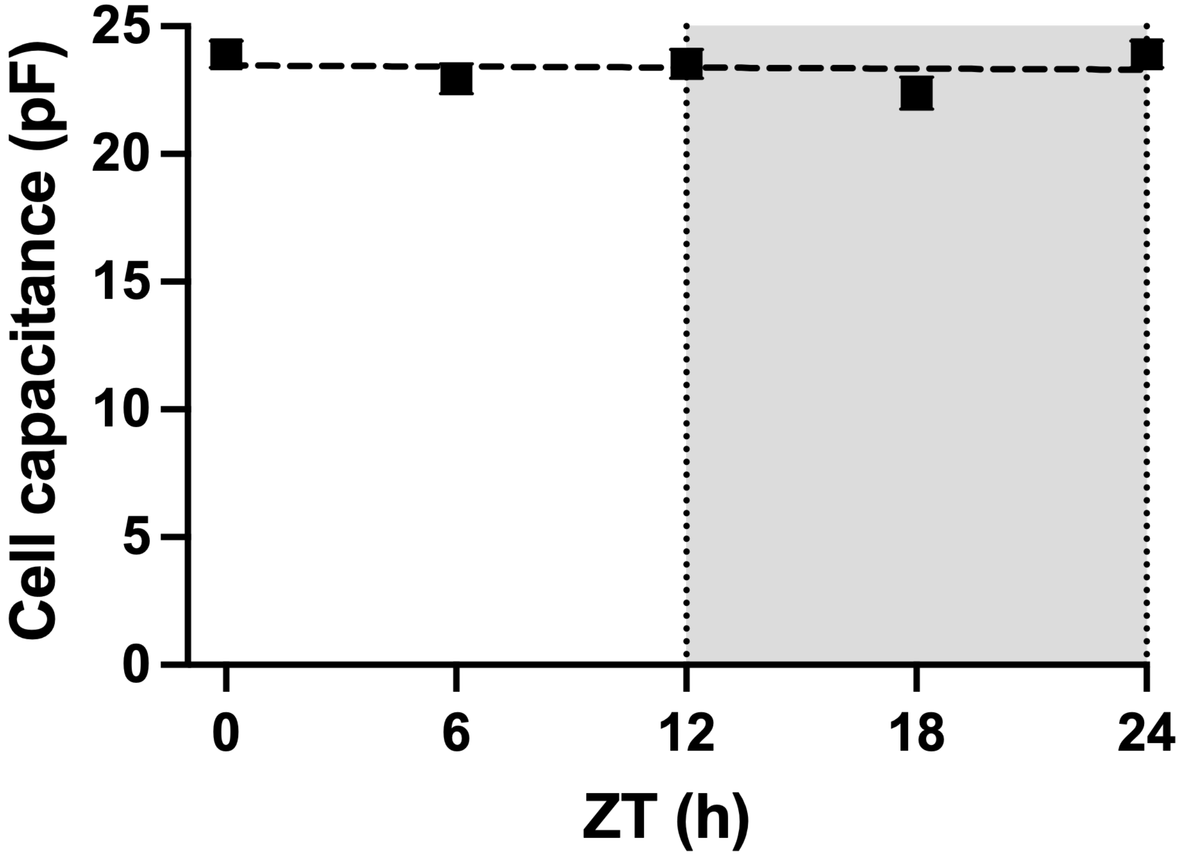


**Fig. S6. No day-night change in the cell capacitance of AV node myocytes.** The mean±SEM capacitance of AV node myocytes at ZT0, 6, 12 and 18 shown (n=54-69 myocytes at each time point; ZT24 data repeat of ZT0 data; standard error bars fall within the plotted points). No significant differences found (Kruskal-Wallis test).

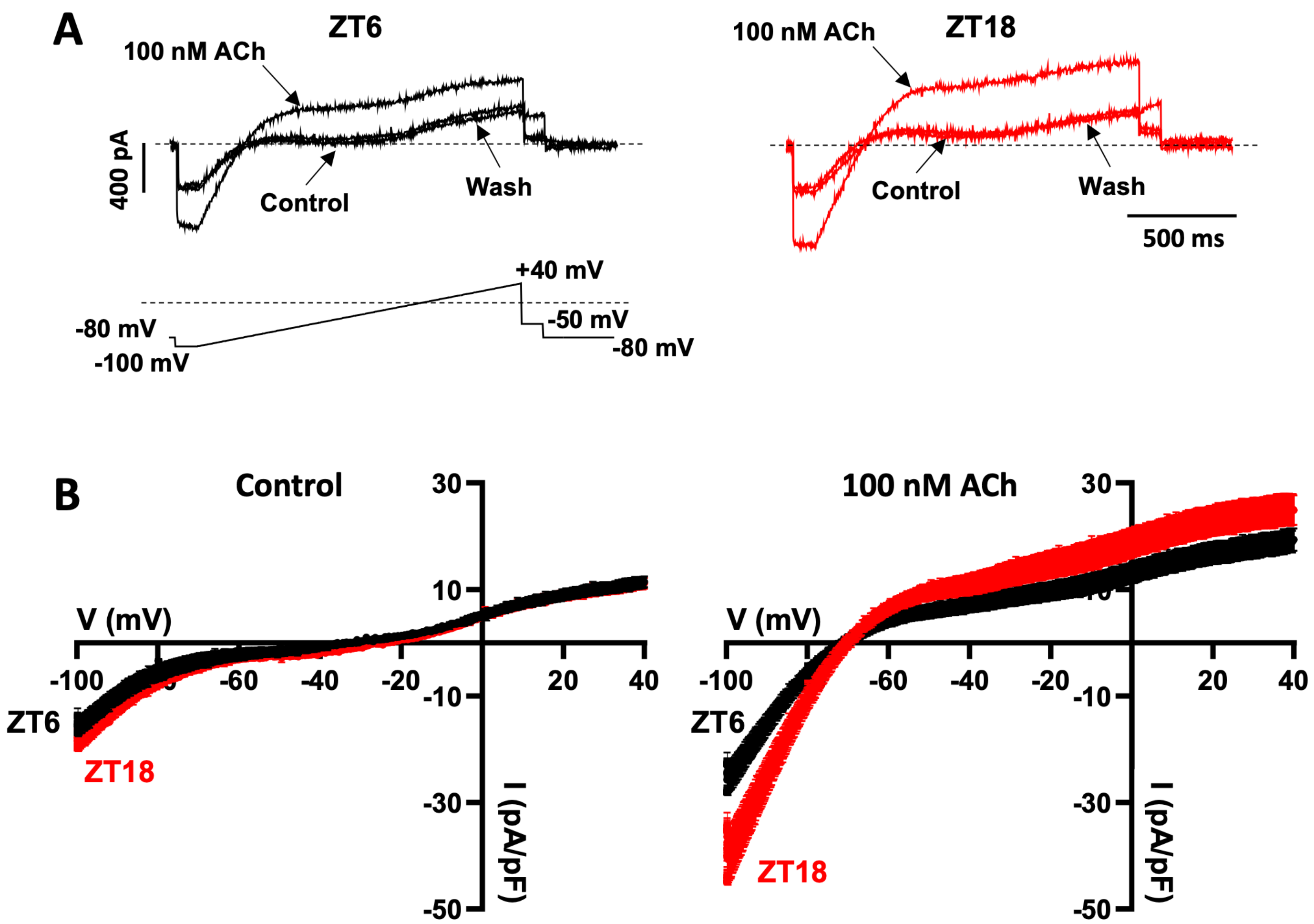


**Fig. S7. Measurement of the ACh-dependent K$^+$ current, $I_{K,ACh}$.** **A**, examples of current recorded during a ramp clamp before and after the application of 100 nM ACh (at ZT6 and ZT18 in this case). **B**, mean current-voltage relationships under control conditions and in the presence of 100 nM ACh (at ZT6 and ZT18 in this case). Mean±SEM current is plotted against the membrane potential during the ramp clamp (ZT6, n=14; ZT18, n=20). $I_{K,ACh}$ was calculated by subtracting the current under control conditions from the current in the presence of ACh at each membrane potential during the ramp clamp and is shown in Fig. 6H.

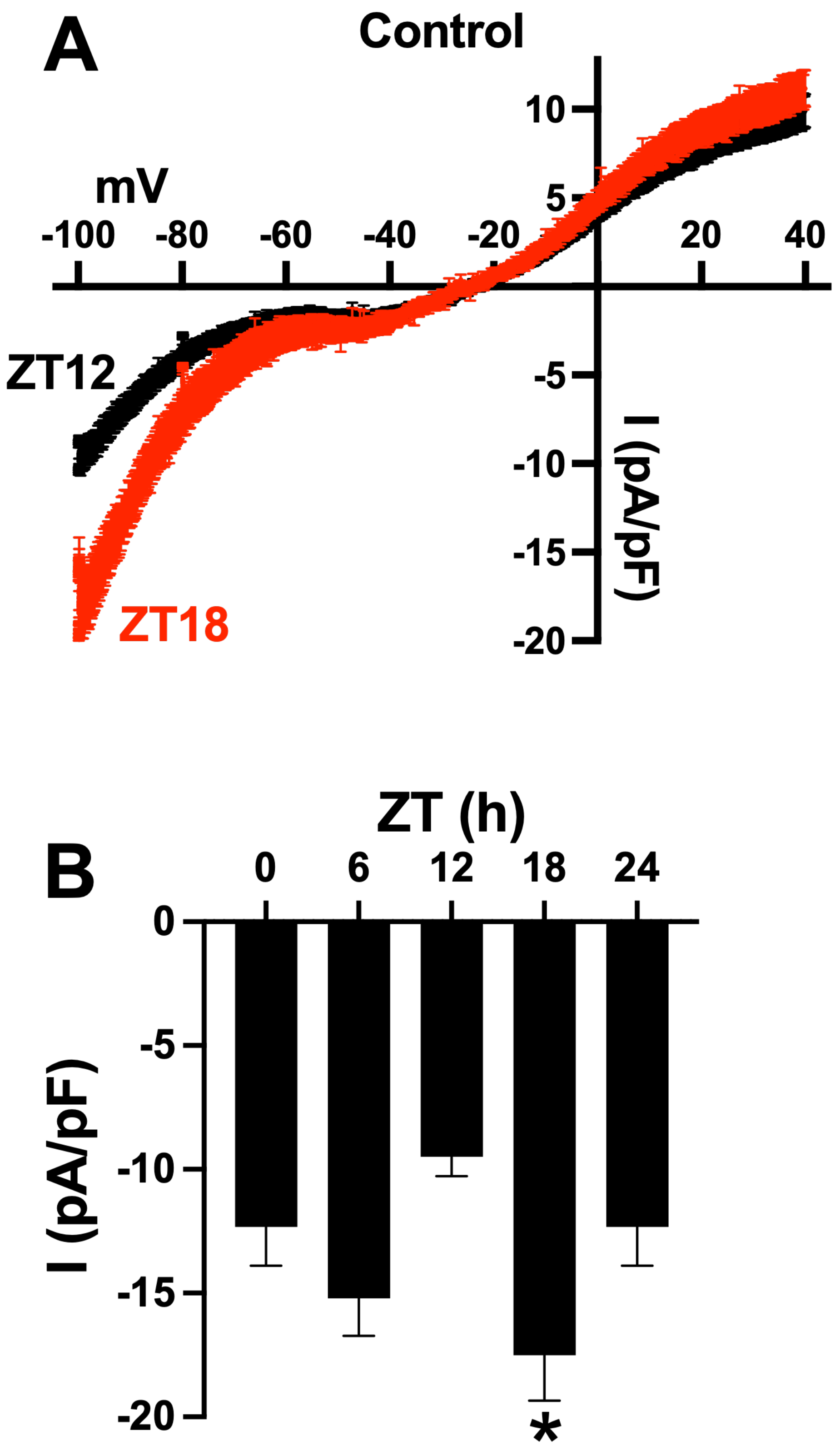


**Fig. S8. Circadian rhythm in background current. A**, mean current-voltage relationships under control conditions at ZT12 and ZT18 obtained from ramp clamps as shown in Fig. S7. Mean±SEM current is plotted against the membrane potential during the ramp clamp (ZT12, n=16; ZT18, n=20). **B**, mean±SEM current at -100 mv at ZT0, 6, 12 and 18 (data at ZT0 is duplicated at ZT24); n=14-20 AV node myocytes at each time point. *significantly different from the data at ZT12; P<0.005; one-way ANOVA followed by Tukey's multiple comparisons test.

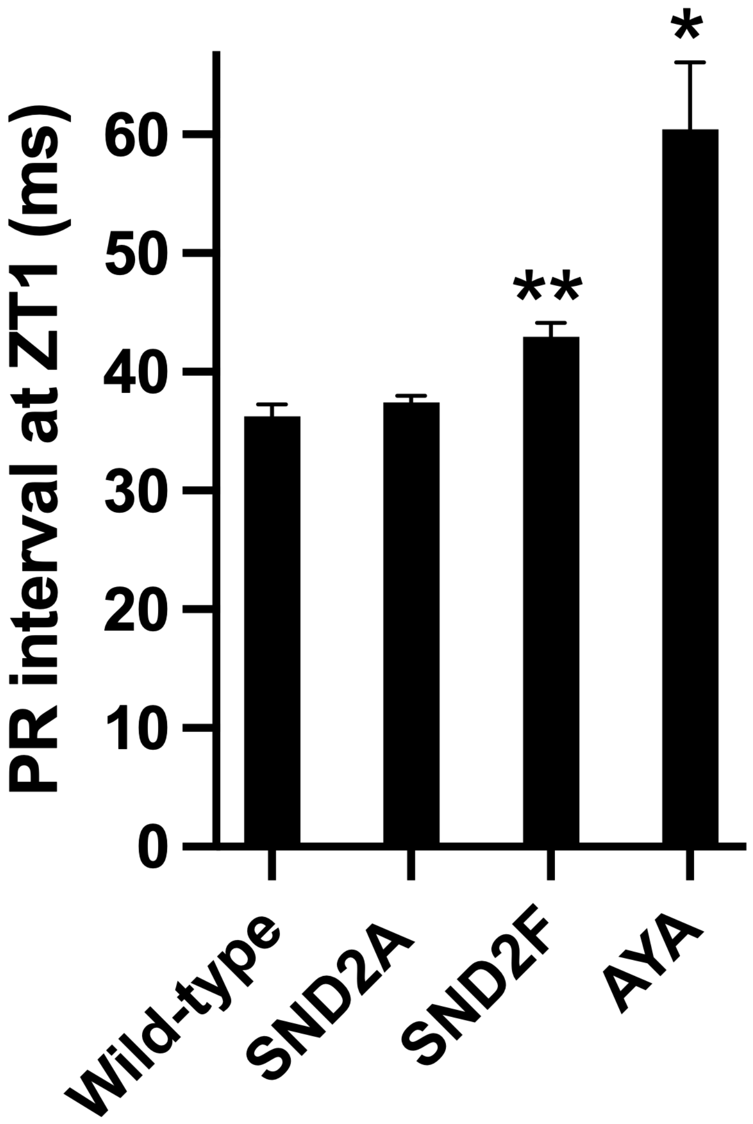


**Fig. S9. Mean±SEM PR interval measured in wild-type and mutant mice.** Wild-type mouse, n=16; SND2A mouse, n=19; SND2F mouse, n=14; AYA mouse, n=7. The SND2A and SND2F mice expressed dominant negative conducting mutants of *Hcn4* lacking the cAMP-dependent regulation of gating. The AYA mouse expressed a non-conducting dominant negative mutant of *Hcn4*. *significantly different from the data from wild-type, SND2A and SND2F mice, $P<0.0001$; **significantly different from the data from wild-type mice, $P<0.05$; one-way ANOVA followed by Tukey's multiple comparisons test.

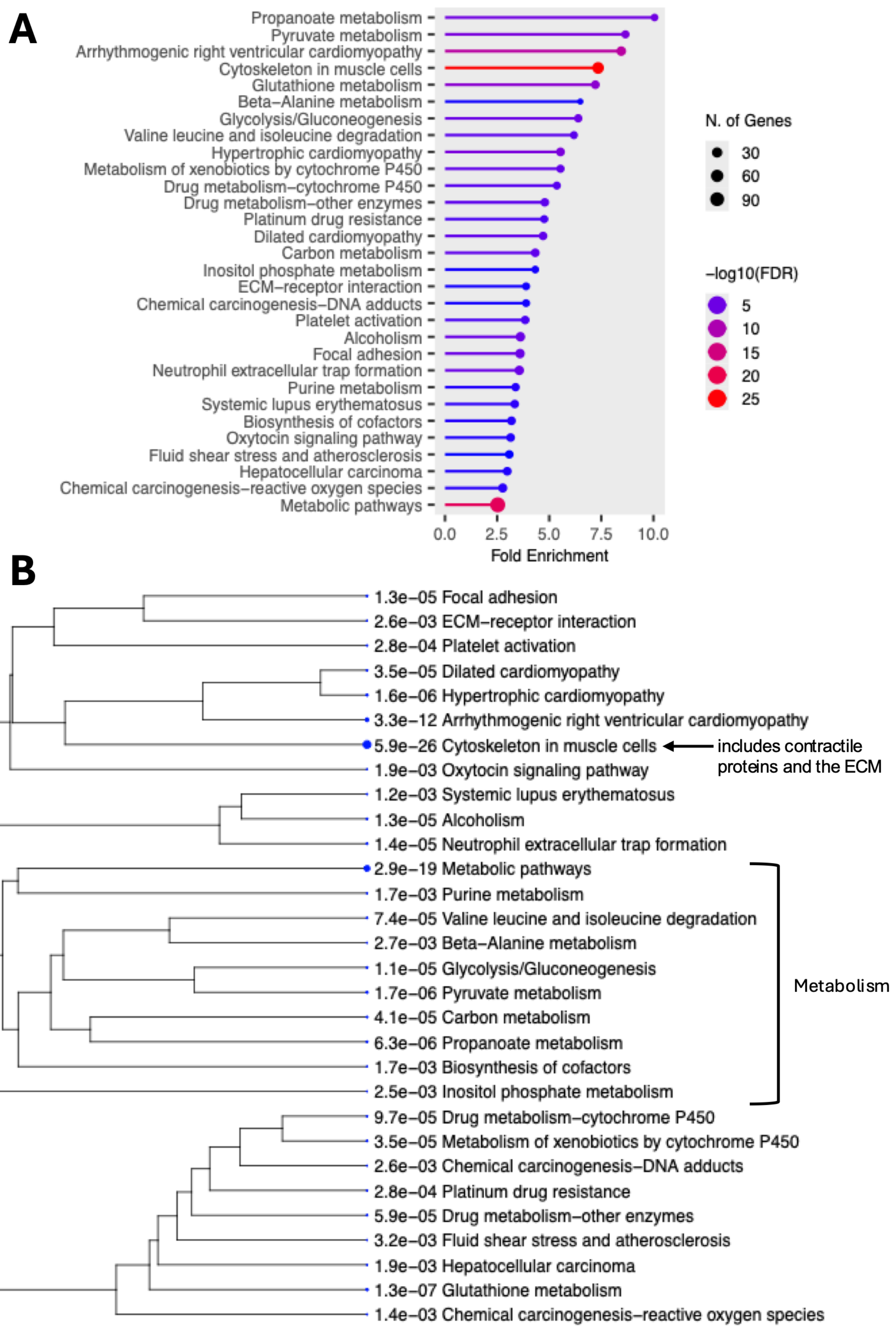


**Fig. S10. Gene ontology analysis of proteins showing a significant circadian rhythm in the AV node.** Generated using ShinyGO 0.85.1. **A**, bar chart of pathways showing a significant circadian rhythm listed according to the fold enrichment of the pathways. The fold enrichment of a pathway is the ratio of the observed number of proteins in a specific pathway showing a circadian rhythm, compared to what is expected by random chance. The number of proteins involved is shown by the size of the spots and the statistical significance is shown by colour. **B**, hierarchical clustering tree summarising the correlation among pathways showing a significant circadian rhythm. Pathways with many shared genes are clustered together. Bigger dots indicate more significant P-values.

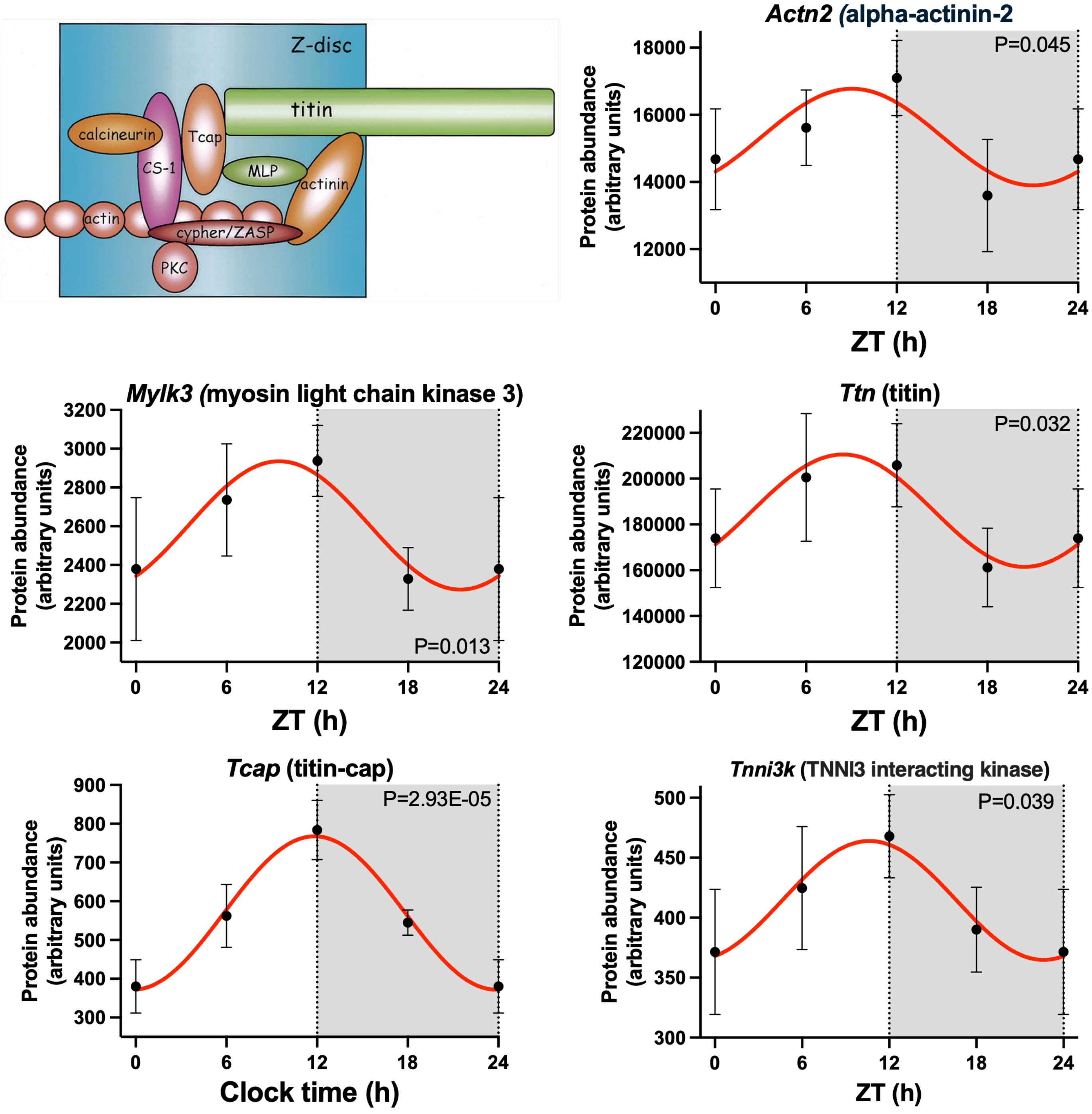


**Fig. S11. Circadian rhythm in contractile proteins.** Inset, schematic diagram showing the arrangement of alpha-actinin-2, titin and titin-cap from Hyashi *et al.*[68] See the legend to Fig. S5 for a general description of the figure. Each point is for four (ZT0, 12 and 18) or five (ZT6) biological replicates, each replicate being from the AV nodes of two mice. Therefore, 8 or 10 mice contributed to each point.

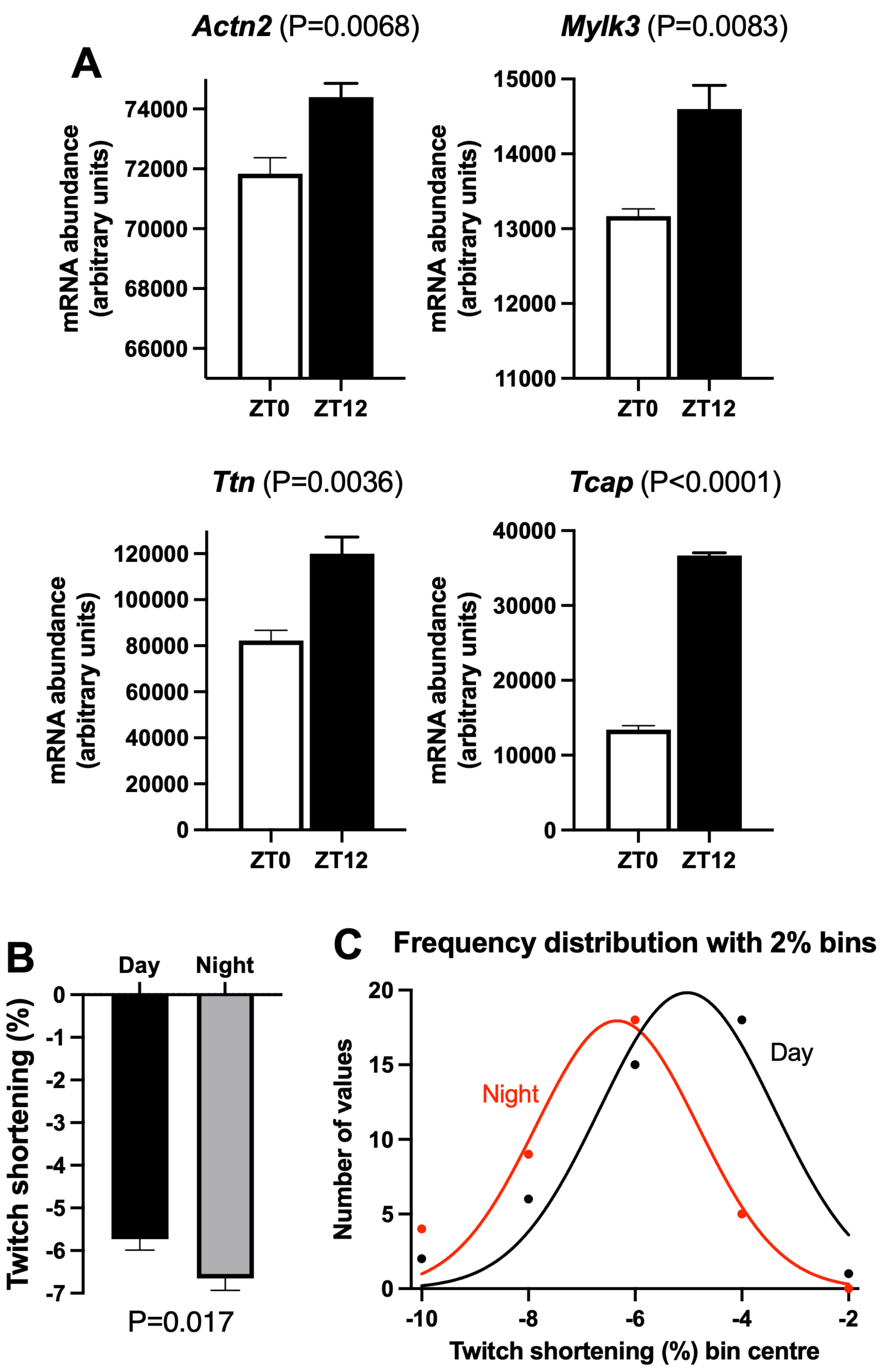


**Fig. S12. Transcripts for four of the contractile proteins showing a significant circadian rhythm in the AV node show a similar circadian rhythm in the ventricles and the potential consequence of this.** **A**, mean±SEM abundance of transcripts responsible for four of the proteins shown in Fig. S11 (data for *Tnni3k* not significantly different) at ZT0 and ZT12 in mouse ventricle; n=5 mice at both timepoints. Data from Tikhomirov *et al.*[39] **B**, mean±SEM twitch shortening of rat ventricular myocytes during the day and at night (day, n=42 myocytes; night, 36 myocytes). P values shown (unpaired t test with Welch's correction). **C**, distribution of twitch shortening magnitudes during the day and night; same data as panel B.

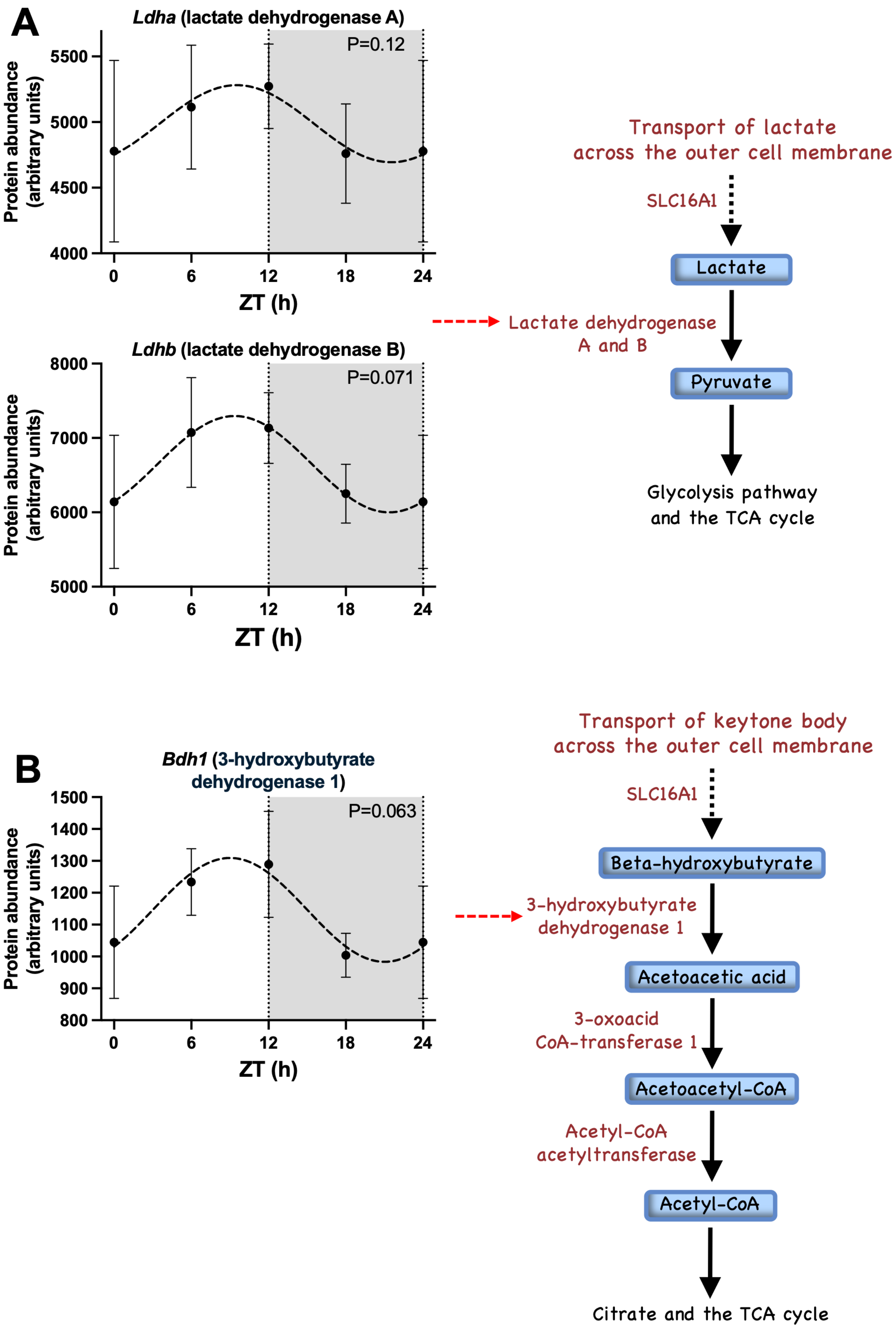


**Fig. S13. Potential circadian rhythm in the lactate (A) and ketolysis (B) pathways.** See the legend to Fig. S5 for a general description of the figure. Each point is for four (ZT0, 12 and 18) or five (ZT6) biological replicates, each replicate being from the AV nodes of two mice. Therefore, 8 or 10 mice contributed to each point. Insets, schematic diagrams of the two pathways.

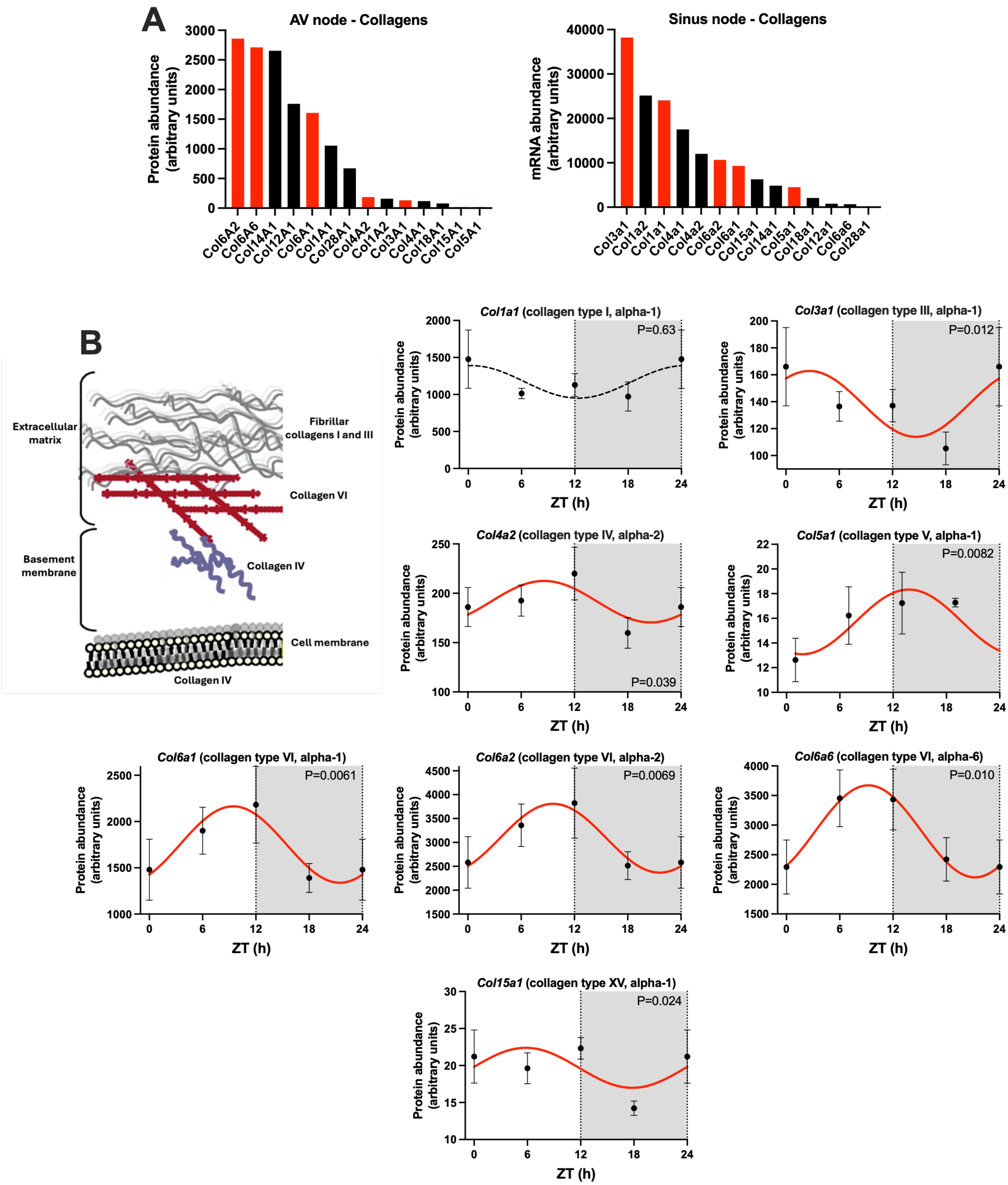


**Fig. S14. Circadian rhythm in extracellular matrix proteins. A**, 24 h mean abundance of collagen isoforms in the mouse AV node and sinus node. Data for the sinus node from Wang *et al.*[29] Red, transcripts showing a significant circadian rhythm; black, transcripts not showing a significant circadian rhythm. **B**, circadian rhythms in collagen isoforms in the mouse AV node; see the legend to Fig. S5 for a general description of the graphs. Each point is for four (ZT0, 12 and 18) or five (ZT6) biological replicates, each replicate being from the AV nodes of two mice. Therefore, 8 or 10 mice contributed to each point. Inset, arrangement of key collagens in the extracellular matrix.

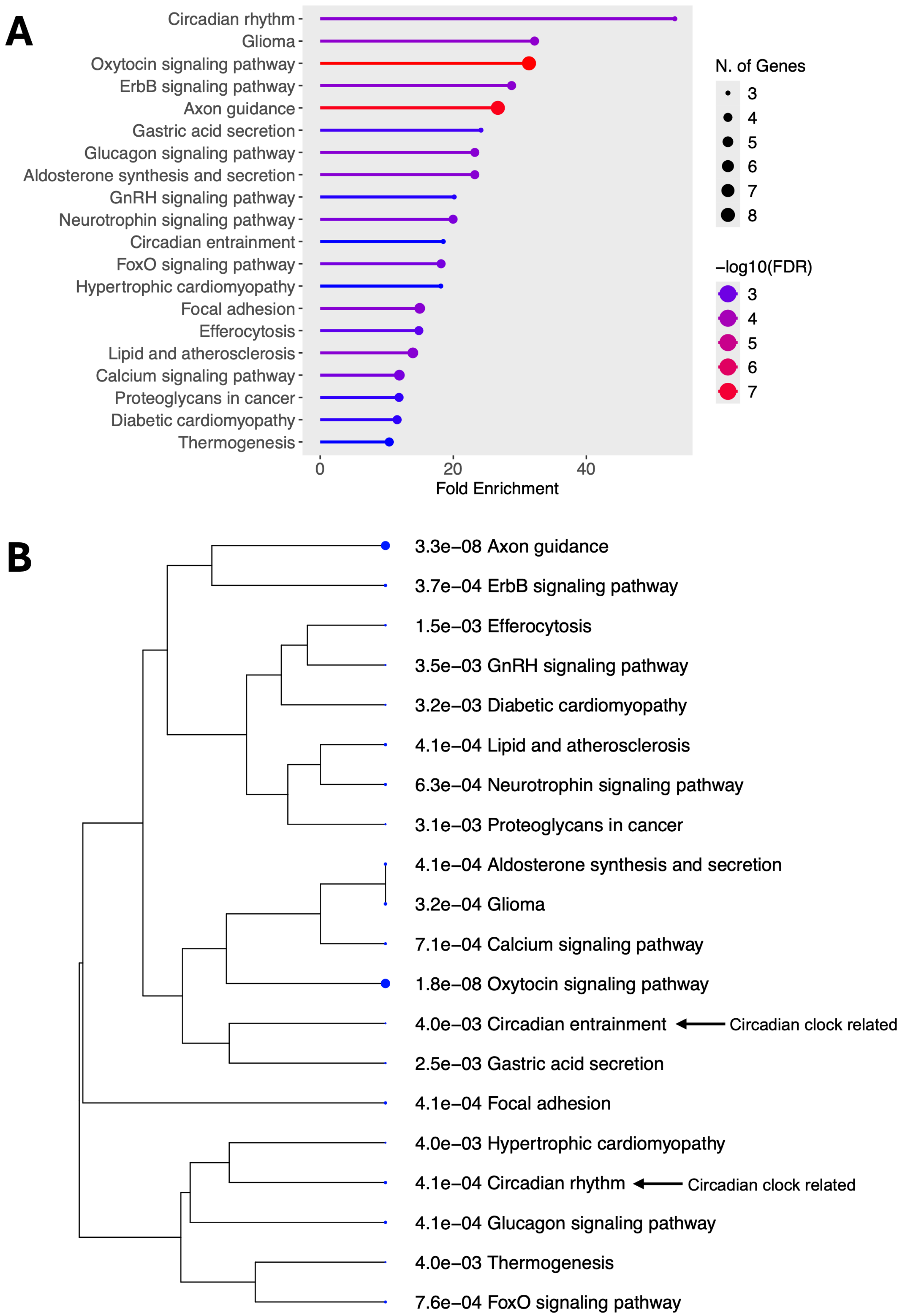


**Fig. S15. Gene ontology analysis of protein kinases showing a significant circadian rhythm.** See the legend to Fig. S10 for a general description of the figure.

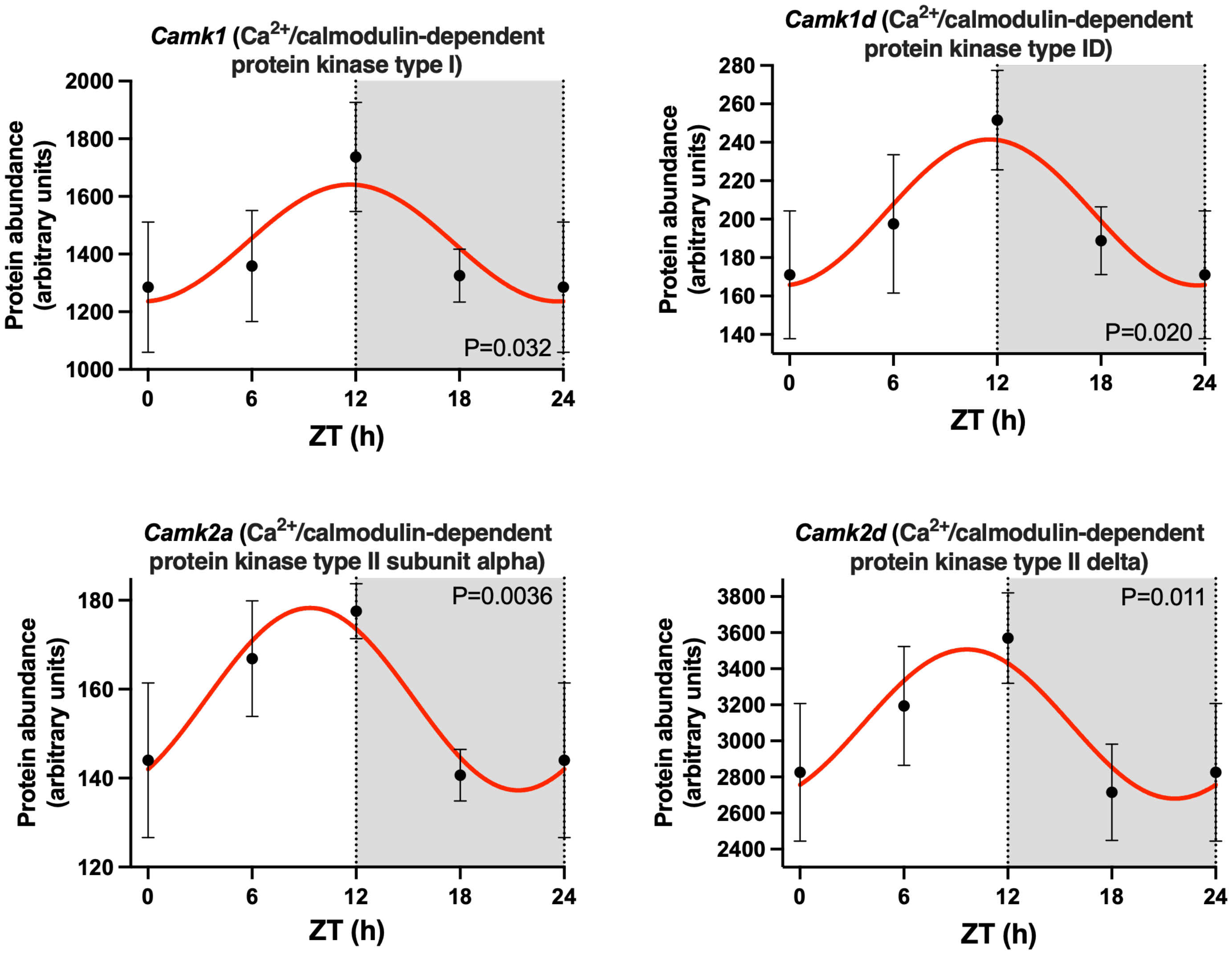


**Fig. S16. Circadian rhythm in $Ca^{2+}$/calmodulin-dependent protein kinases at the protein level.** See the legend to Fig. S5 for a general description of the figure. Each point is for four (ZT0, 12 and 18) or five (ZT6) biological replicates, each replicate being from the AV nodes of two mice. Therefore, 8 or 10 mice contributed to each point.

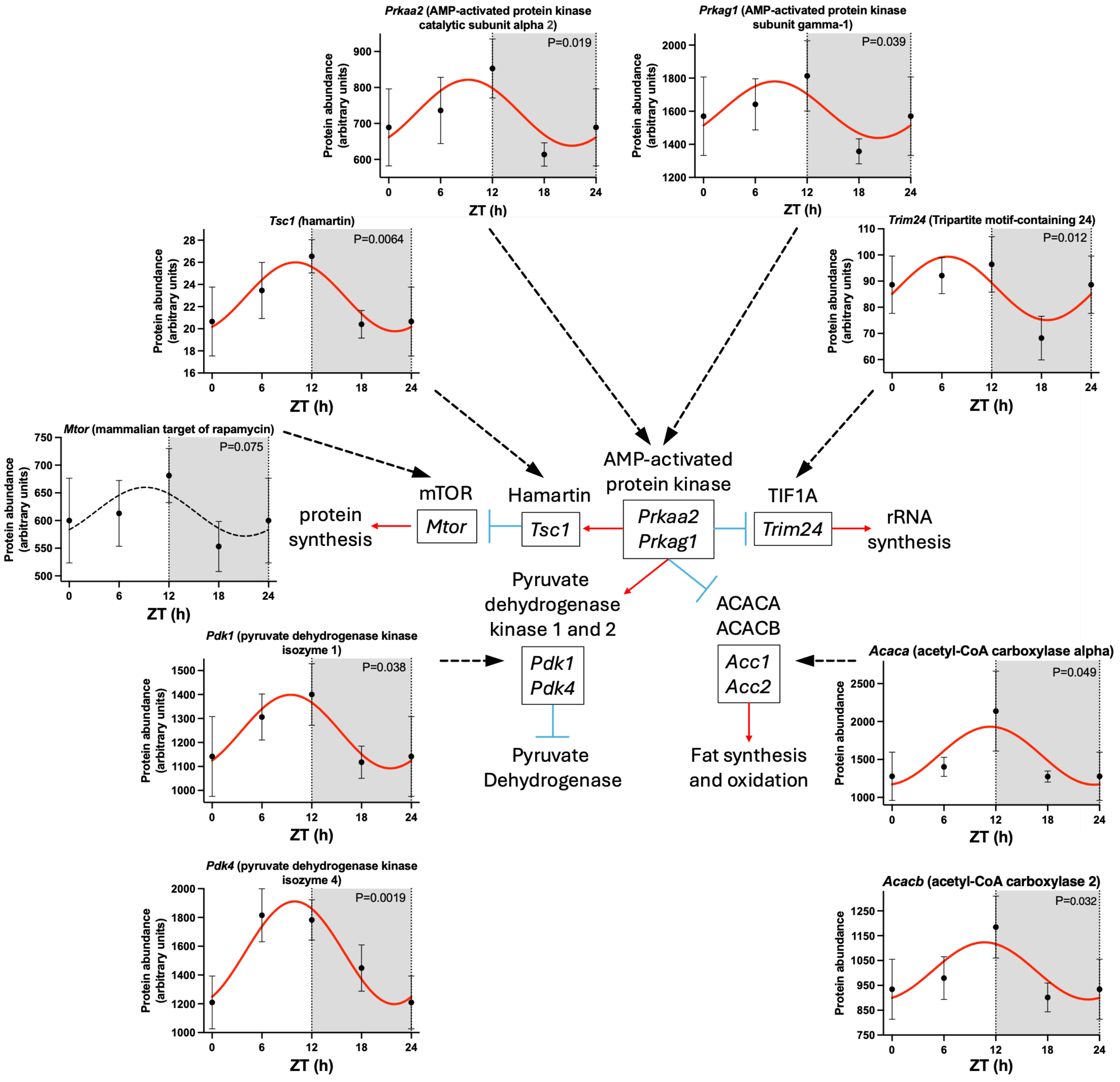


**Fig. S17. AMP-activated protein kinase pathway.** See the legend to Fig. S5 for a general description of the figure. Each point is for four (ZT0, 12 and 18) or five (ZT6) biological replicates, each replicate being from the AV nodes of two mice. Therefore, 8 or 10 mice contributed to each point. Inset, schematic diagram of the pathway. Red arrows indicate activation of the downstream protein or process and blue lines indicate inhibition.

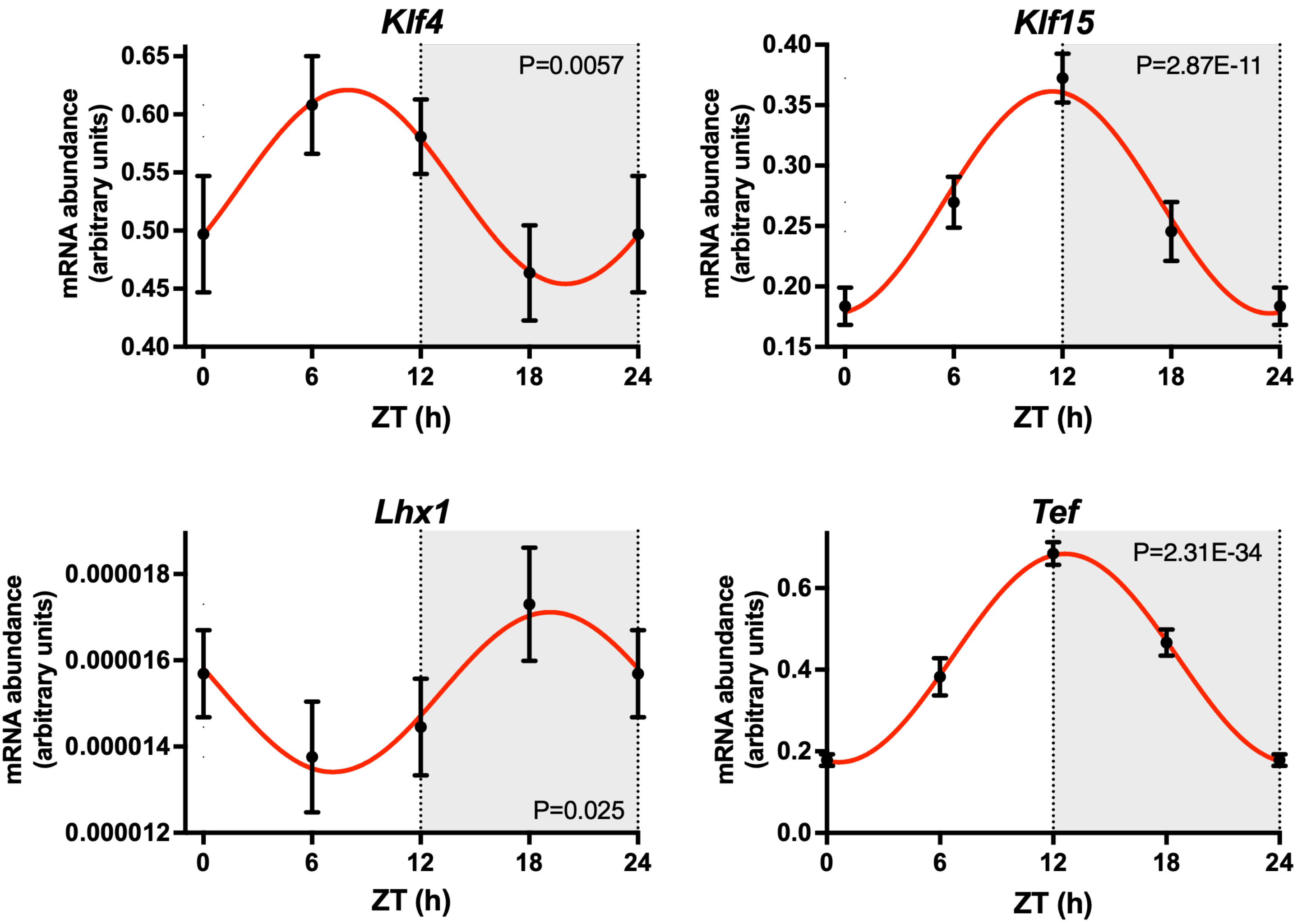


**Fig. S18. Circadian rhythm in four circadian clock "slave" transcription factor transcripts.** See the legend to Fig. S5 for a general description of the figure. Data from 8-11 mice.

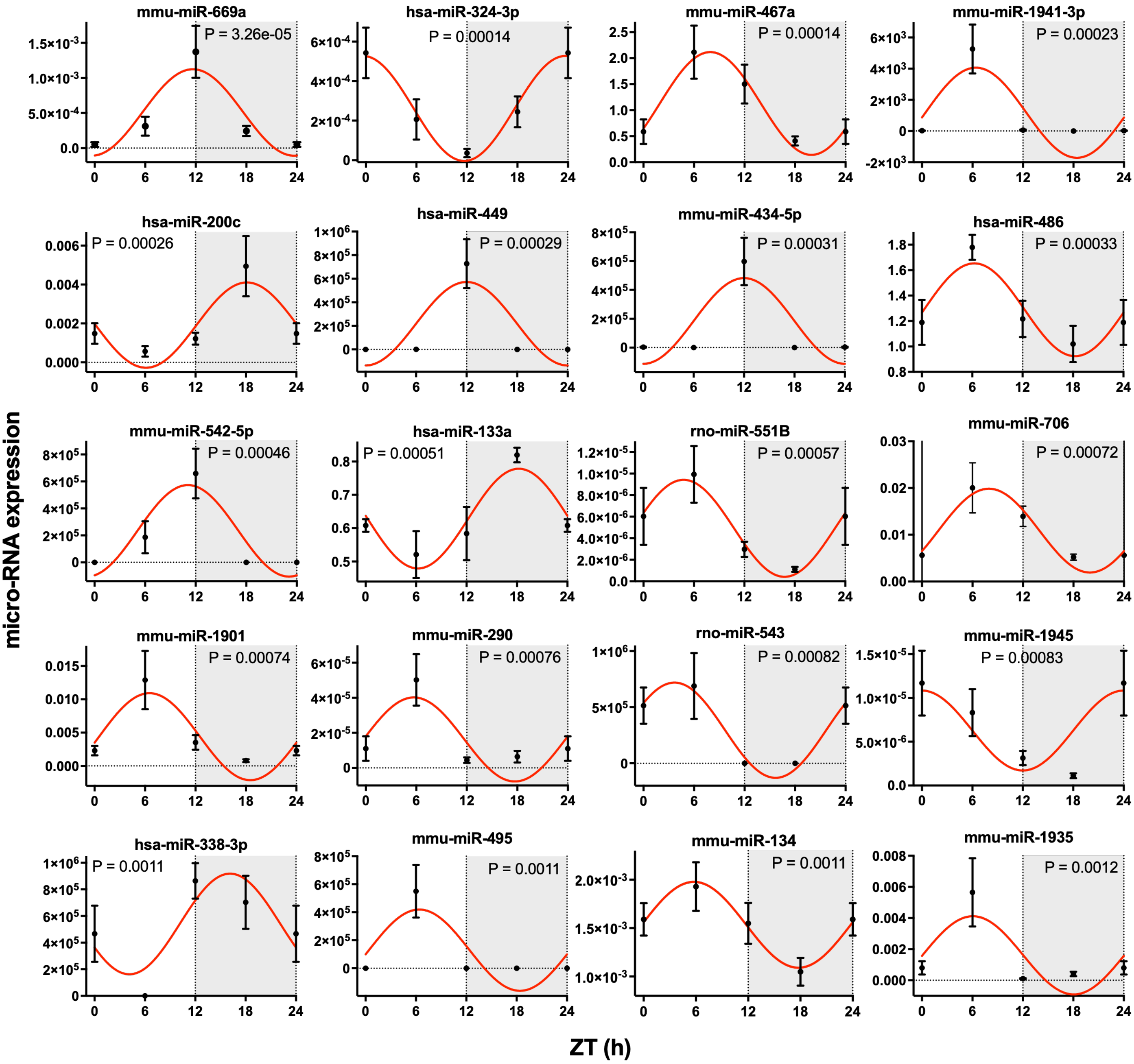


**Fig. S19. Examples of circadian rhythms in micro-RNAs (micro-RNAs showing the most significant circadian rhythms chosen for display).** See the legend to Fig. S5 for a general description of the figure. Mean±SEM micro-RNA abundance shown (n=4-8) at four time points over 24 h (24 h data repeat of 0 h data).

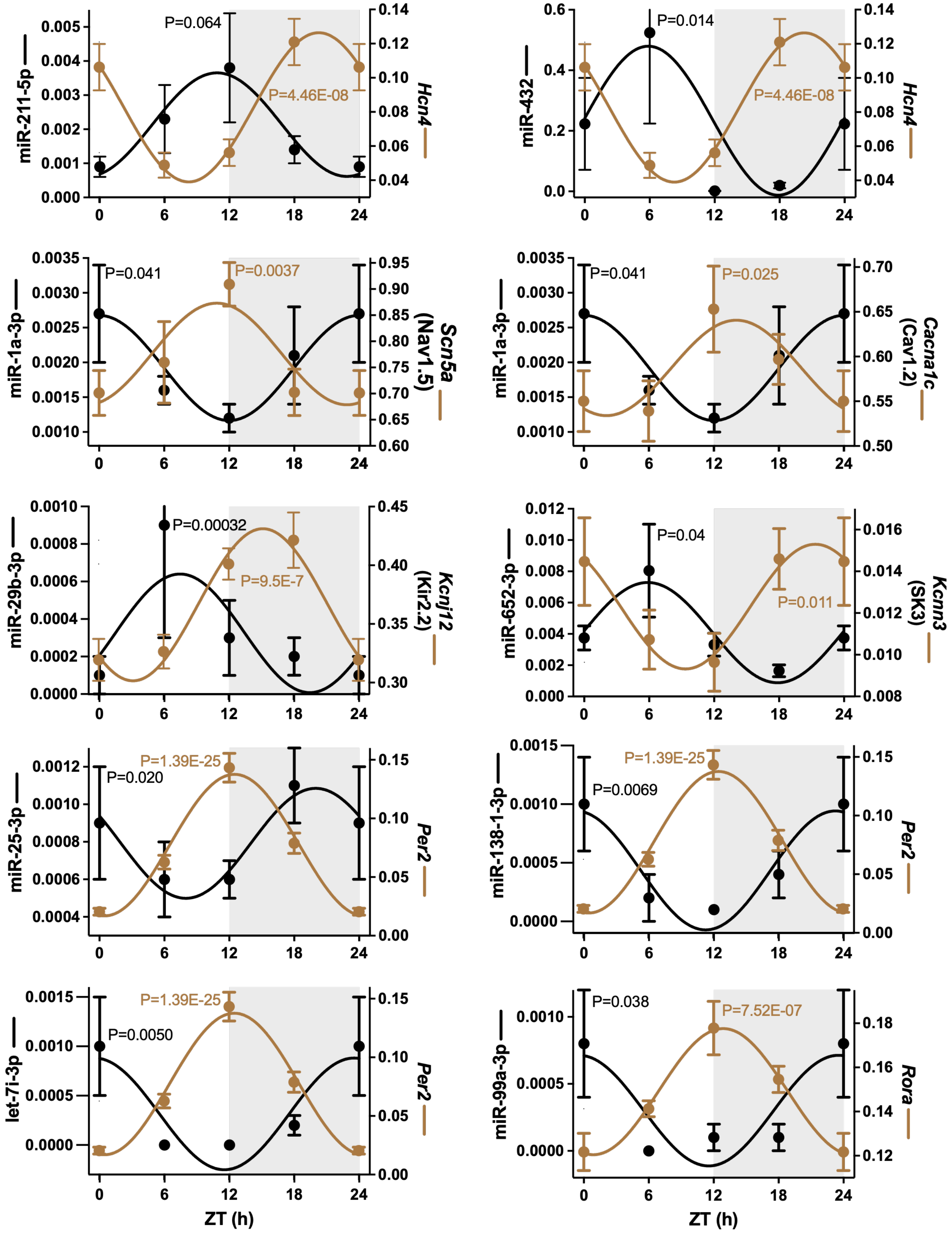


**Fig. S20. Examples of circadian rhythms in micro-RNAs and their proven or potential target transcripts.** Examples are shown of micro-RNAs with significant circadian rhythms (data shown in black) and the significant circadian rhythms of their proven or known target transcripts (data shown in brown). Examples have been chosen where the micro-RNA and target transcript are approximately in antiphase. See Fig. S5 legend for a general description of the figure. Mean±SEM micro-RNA abundance shown (n=3-8) at ZT 0/24, 6, 12 and 18.

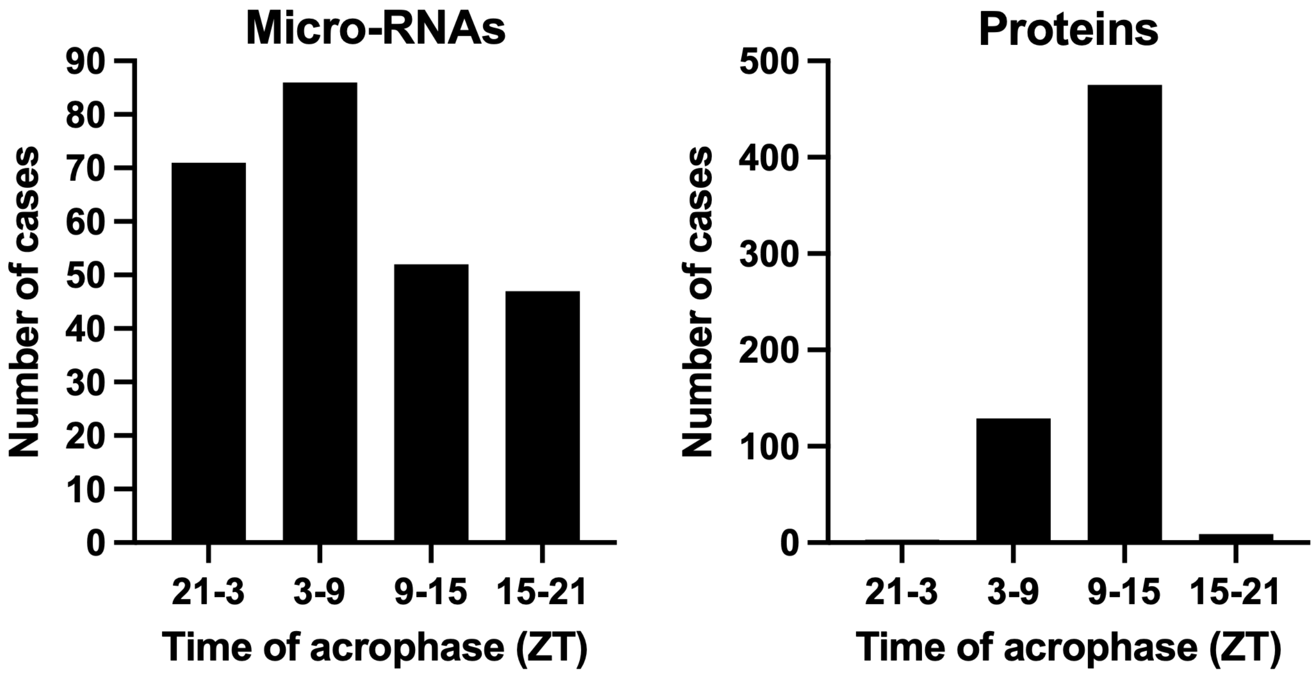


**Fig. S21. Histogram of the time of acrophase (time of peak expression) of micro-RNAs and proteins showing a significant circadian rhythm in the AV node.**

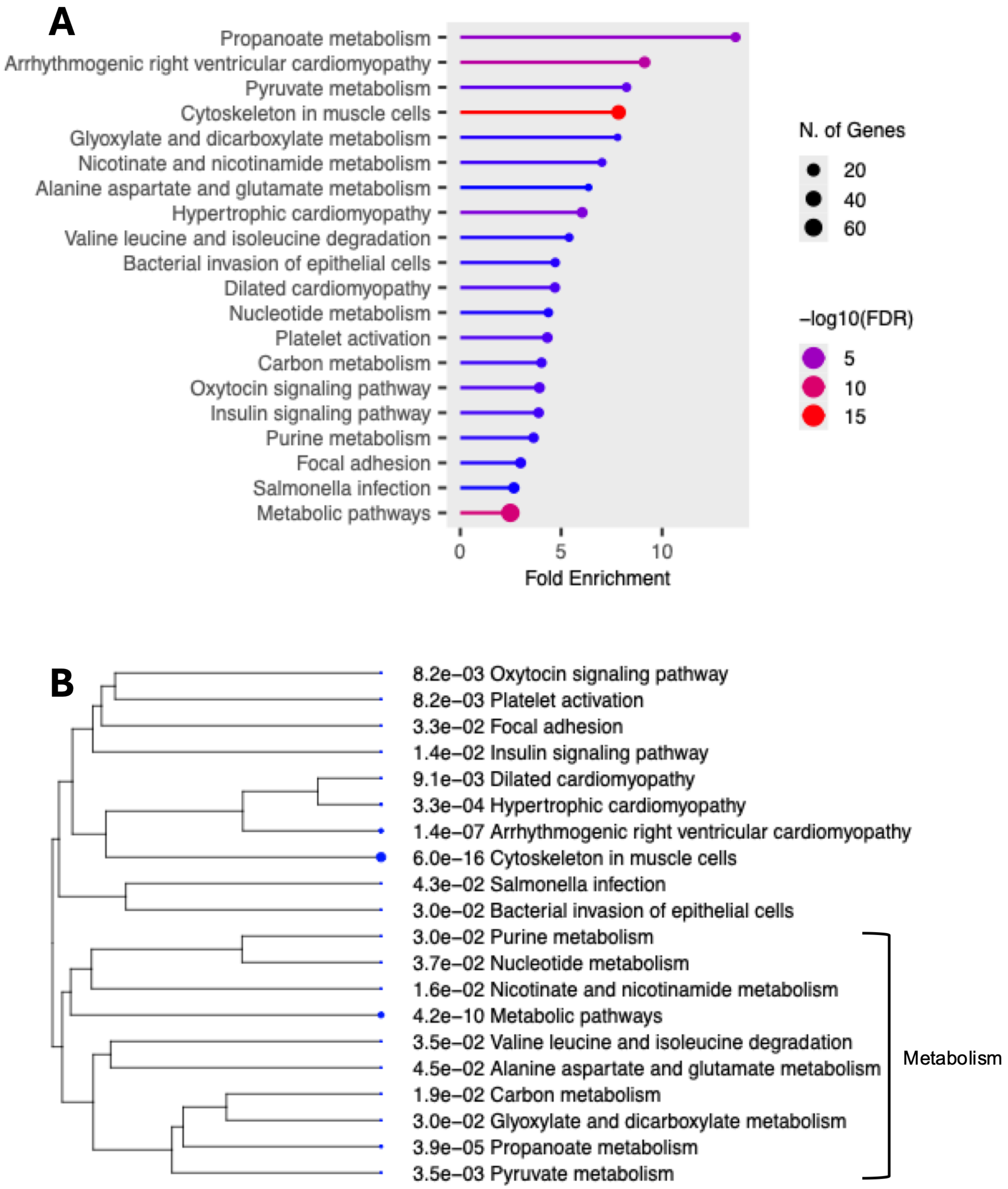


**Fig. S22. Gene ontology analysis of potential target proteins showing a significant circadian rhythm of micro-RNAs showing a significant circadian rhythm.** See the legend to Fig. S10 for a general description of the figure.

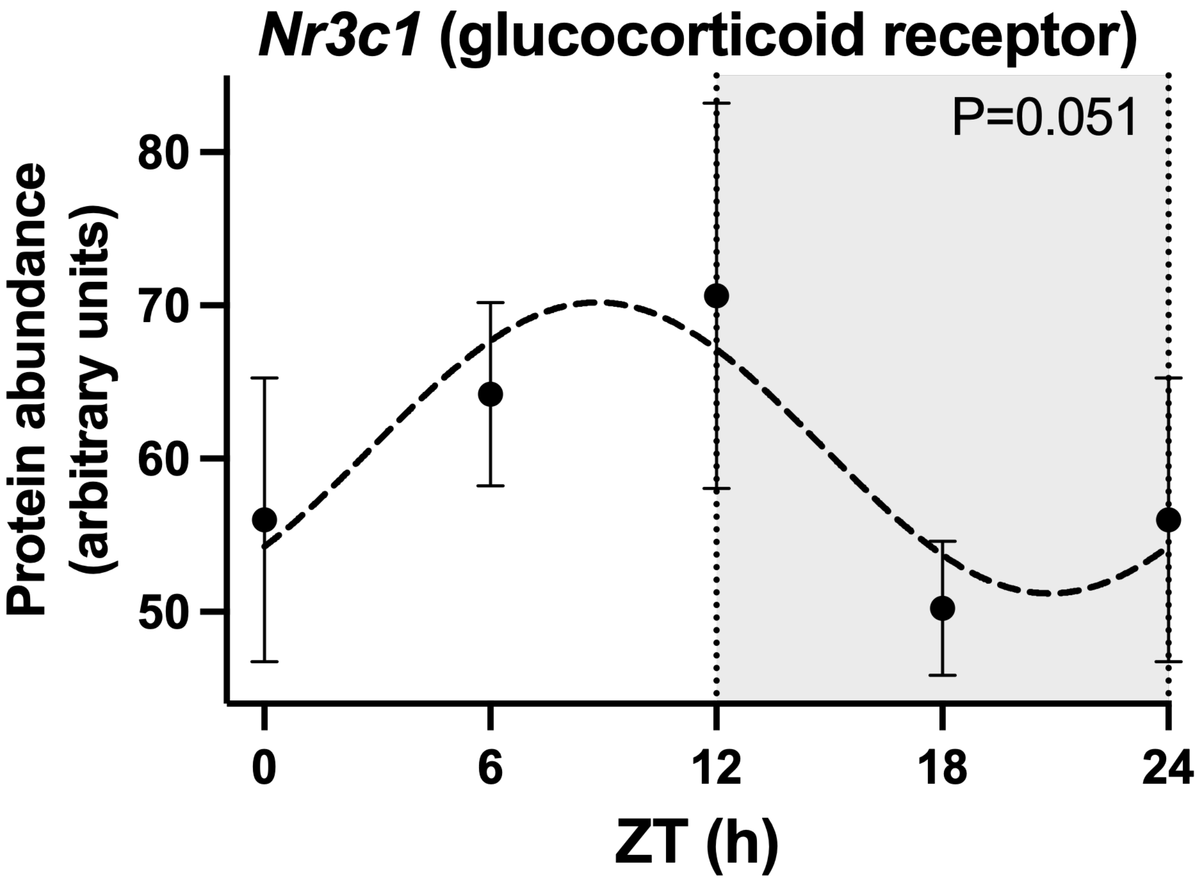


**Fig. S23. Circadian rhythm in glucocorticoid receptor protein.** See the legend to Fig. S3 for a general description of the figure. Each point is for four (ZT0, 12 and 18) or five (ZT6) biological replicates, each replicate being from the AV nodes of two mice. Therefore, 8 or 10 mice contributed to each point.
